# qBiCo: A method to assess global DNA conversion performance in epigenetics via single-copy genes and repetitive elements

**DOI:** 10.1101/2024.08.29.610354

**Authors:** Faidra Karkala, Roy B. Simons, Floor Claessens, Vivian Kalamara, Manfred Kayser, Athina Vidaki

## Abstract

Human DNA methylation profiling offers great promises in various biomedical applications, including ageing, cancer and even forensics. So far, most DNA methylation techniques are based on a chemical process called sodium bisulfite conversion, which specifically converts non-methylated cytosines into uracils. However, despite the popularity of this approach, it is known to cause DNA fragmentation and loss affecting standardization, while incomplete conversion may result in potential misinterpretation of methylation-based outcomes. To offer the community a solution, we developed qBiCo - a novel quality-control method to address the quantity and quality of bisulfite-converted DNA. qBiCo is a 5-plex, TaqMan^®^ probe-based, quantitative (q)PCR assay that amplifies single- and multi-copy DNA fragments of converted and non-converted nature. It estimates four parameters: converted DNA concentration, fragmentation, global conversion efficiency, and potential PCR inhibition. We optimized qBiCo using synthetic DNA standards and assessed it using standard developmental validation criteria, showcasing that qBiCo is reliable, robust and sensitive down to picogram level. We also evaluated its performance by testing decreasing DNA amounts using several commercial bisulfite conversion kits. Depending on the starting DNA quantity, bisulfite-converted DNA recoveries ranged from 8.5 – 100 %, conversion efficiencies from 78 - 99.9 %, while certain kits highly fragment DNA, demonstrating large variability in their performance. Towards building a prototype tool, we further optimized key functionalities, for example, by replacing the poorest performing single-plex assay and creating a more representative DNA standard. Aiming to scale-up and move towards implementation, we successfully transferred and validated our novel method in six different qPCR platforms from different major manufacturers. Overall, with the present study, we offer researchers in the epigenetic field a novel long-awaited QC tool that for the first time allows them to measure key quality and quantity parameters of the most popular DNA conversion process. The tool also enables standardization to prevent inconsistent data and false outcomes in the future, regardless of the downstream experimental analysis of DNA methylation-based research and applications across different fields of biology and biomedicine.

## Introduction

Epigenetics involves heritable modifications in gene expression that are not depicted by changes in DNA sequence, but rather appear as base and histone modifications [1]. DNA methylation is the most studied epigenetic modification, that chemically affects DNA by covalently adding a methyl group to the 5’ carbon of cytosine. In humans, the majority of DNA methylation appears on cytosine-guanine dinucleotides (5’-CpG-3’) [2]. As response to internal and external stimuli [3], DNA methylation regulates gene transcription by usually silencing a gene, while a non-methylated gene remains active and accessible to the transcription machinery [4]. Profusion of emerging evidence indicates the importance of DNA methylation as a molecular biomarker for detecting and monitoring phenotypes associated to lifestyle choices [5], ageing and age-related diseases [6] including cancer [7, 8] and cardiovascular disorders [9] as well as in neurodevelopmental disorders [10]. Furthermore, DNA methylation profiling can also be useful in more specialized applications, such as forensics [5, 11, 12].

To analyse genome-wide or targeted methylation, several high-throughput methods have been developed depending on the research question and downstream analysis. However, the vast majority require chemical treatment of DNA via the so-called bisulfite conversion (BC), a process where during DNA treatment with sodium bisulfite (NaHSO_3_), non-methylated cytosines are converted to uracils, whereas methylated cytosines remain unchanged [13]. Hence, follow-up PCR amplification of bisulfite-converted DNA (BC-DNA) replaces uracils with thymines, allowing the detection of methylation as an introduced C/T variant using standard downstream technologies, including microarrays, pyrosequencing and massively parallel sequencing [14]. Thus, due to its simplicity, cost-effectiveness and compatibility with various downstream genetic pipelines, BC has dominated the field over the last decades over enzymatic or immunoprecipitation approaches and is currently considered as the gold standard to study DNA methylation [14]. Additionally, it is the preferred method as it enables the analysis of any CpG of interest, without restrictions to specific enzyme recognition motifs and required enzyme-specific optimization [15].

Despite its popularity, however, BC requires aggressive chemical conditions such as low pH and high temperature, which can affect the integrity of DNA, resulting in damage and fragmentation. At the same time, choosing less aggressive conditions can affect the success and efficiency of conversion, resulting in follow-up overestimation of methylation and data misinterpretation [16]. When BC-DNA is too fragmented or ‘contaminated’ with carryover chemicals from the treatment, its detection and interpretation is also affected. Specifically, DNA incubation with sodium bisulfite results in depyrimidination and generation of abasic sites that are prone to DNA strand breaks, especially under heating or alkaline conditions [17]. Increased DNA fragmentation decreases the number of DNA molecules available for PCR amplification, longer targets are less likely to amplify, and eventually, the PCR results are no longer reproducible due to the large variability driven by stochastic events early during PCR [18].

Nowadays, epigenetic researchers perform BC via commercially available kits from various manufacturers, each coming with a slightly different treatment protocol. Generally, according to the manufacturers statements, commercial BC kits promise > 99 % DNA conversion rate, at least 80 % recovery rate of BC-DNA, and eluted DNA fragments of up to 2,000 bp in length. Thus far, researchers have attempted to empirically evaluate these parameters using small-scale and/or non-specific approaches. Specifically, the quantity of BC-DNA is usually assessed via Nanodrop spectrophotometer (Thermo Fisher Scientific - TFS), Qubit fluorometer (TFS), or Bioanalyzer (Agilent), while the degree of fragmentation of BC-DNA is via Bioanalyzer or agarose gel electrophoresis. When comparing the performance of different commercial BC kits using these methods, researchers concluded that key quantity and quality parameters strongly depend on the initial input amount of genomic DNA (gDNA) and the incubation time during BC [18–22]. Importantly, none of these methods is specific to BC-DNA, which is short, single/double-stranded hybrid, and contains ‘contaminating’ compounds and secondary structures that affect downstream analysis. Overall, lacking quality- and quantity-related indications does not only affect the accuracy, reproducibility and robustness of DNA methylation detection, but also the evaluation of epigenetic-based outcomes [23].

In an early effort, Ehrich *et al.* [24] developed multiple single-locus PCR assays for assessing post-conversion DNA quality in terms of fragmentation under increasing incubation temperature during bisulfite treatment. While this study showcased the impact of high temperature (> 65 °C) on degrading BC-DNA fragments (< 500 bp), no indication on BC efficiency or BC-DNA recovery could be devised. On the other hand, Sriraska *et al.* [25] aimed to verify the BC rate using a set of genomic and converted-specific primers, with incomplete conversion being detected when lower DNA amounts were used for conversion (< 37.5 ng). However, their approach was based only on a single locus, therefore highly biased and unable to provide with a global DNA conversion measure.

More recently, Kint *et al.* (2018) used a combination of agarose gel electrophoresis, quantitative PCR and digital PCR to measure BC-DNA fragmentation, spectroscopic measurements and digital PCR to evaluate BC-DNA recovery, as well as sequencing to assess BC efficiency [19]. While this study revealed large performance differences among BC kits (for example, recovery ranging from 26.6 to 88.3 %), their multi-level approach is not easy and practical to apply as a simple QC tool in experimental pipelines, as it also consumes large amounts of the often-precious BC-DNA sample.

So far, no BC-DNA-specific method exists that can thoroughly assess the BC process from both the qualitative and quantitative side, leading researchers to simply assume successful performance as stated per manufacturer to sell their products. Overall, the epigenetics community has largely overestimated the method’s capabilities despite the warnings of the inventor of BC [26], Hikoya Hayatsu, who in 2008 wrote: “I now feel that the presently available data for DNA methylation need carefully be looked at, and that a scientifically sound, assured methodology free from any false positive or false negative assignments be established as soon as possible” [27].

In the present study, to offer the epigenetic community with a solution, we developed a novel QC method to simultaneously address the quantity and quality of BC-DNA (qBiCo). It is a 5-plex, TaqMan^®^ probe-based quantitative (q)PCR assay, which amplifies single- and multi-copy fragments of converted and non-converted nature to estimate a sample’s: i) BC-DNA concentration, ii) global BC efficiency, iii) BC-DNA fragmentation and iv) potential PCR inhibition. Here, we describe the initial development of the qBiCo assay, demonstrate its applicability by evaluating several BC kits and describe our strategy moving towards a small-scale qBiCo technology prototype, which we validated using multiple qPCR platforms (technology readiness levels, TRL 1-4).

## Results

### Method formulation based on single- and multi-copy targets

Our envisioned method is based on co-amplifying five DNA regions with five sets of primers and probes labelled with different fluorophores based on TaqMan^®^-based qPCR technology. The five assays were designed to assess the conversion efficiency, quantity, fragmentation and inhibition of a BC-DNA sample (**S1 Table**). The first assay targets a single-copy gene (*hTERT*) via a short PCR fragment corresponding to the BC-DNA concentration index. The second assay targets a longer version of the same single-copy gene (*hTERT*) allowing us to calculate the BC-DNA fragmentation index. The third and fourth assays target the genomic and converted versions of a multi-copy repetitive element (*LINE1*) allowing to estimate a representative, genome-wide conversion efficiency index. The fifth assay targets a spiked-in, short, artificial DNA fragment acting as an internal positive control (IPC) that can give clues on potential PCR inhibition. Importantly, given the difficulty and uncertainty involved with using BC-DNA as standards, we opted for synthesizing artificial DNA fragments for this purpose.

### Development and evaluation using synthetic standards

During the first phase of development, we aimed to create the proof-of-concept version of our novel method (qBiCo-v1). First, we optimized each qPCR assay separately, which resulted in achieving similar performance for all five assays (**Fig 1A**). Moving into a multiplex qPCR reaction, **Fig 1B** indicates successful achievement of harmonized amplification of all assays in a single reaction, where single-copy assays (*hTERT* Short & Long) were detected in the same quantification cycle (Cq), as did the multi-copy assays (*LINE1* Converted & Genomic). Finally, we showed that a maximum of 32 cycles were preferable to avoid false-positive detection driven by low-level background signal, particularly for the very sensitive *LINE1 Genomic* assay that can also amplified in low levels of contaminating non-human DNA (**Fig 1E**).

**Fig 1.**
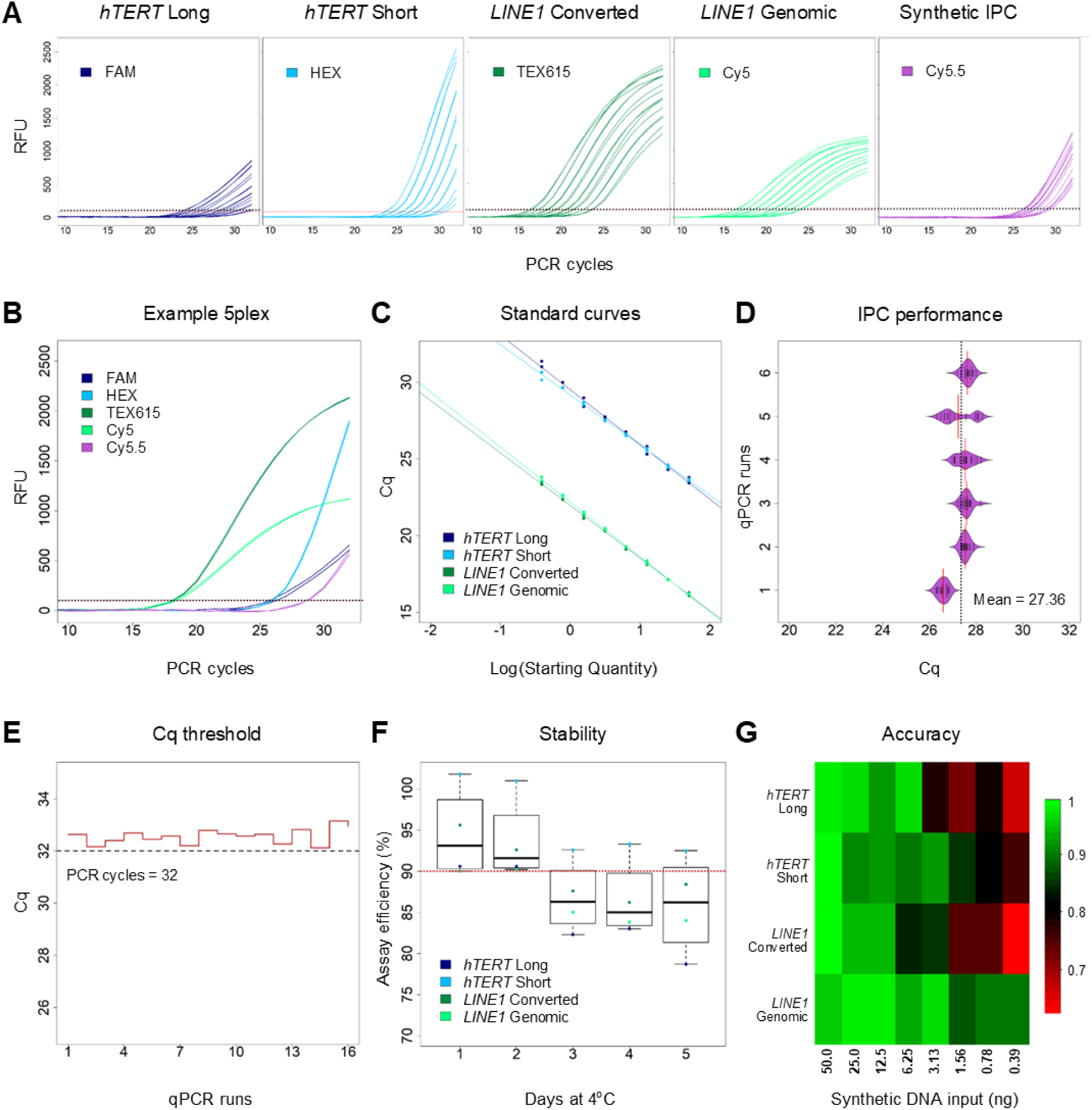
qBiCo-v1 assay development using synthetic DNA standards. (A) Amplification curves of the single qPCR assays for the eight serially diluted DNA standards, each labelled with a different fluorescent dye; (B) Amplification curves of an example 5-plex qBiCo assay (STD3); (C) Standard curves of the four assays (50-0,39 ng/µl) used to calculate the concentration of each fragment in a sample; (D) Spike-in control performance in the serially diluted DNA standards during six qPCR runs; (E) Performance of negative controls during 16 qPCR runs; (F) Assay efficiency using the same DNA standard dilution series within five days; (G) Heatmap of the detection limit of each individual assay based on the serially diluted DNA standards. RFU: Relative fluorescent units; STD: Synthetic standard; IPC: Internal positive control; Cq: Quantification cycle.

Subsequently, the performance of qBiCo was evaluated by studying the PCR efficiency and accuracy of each individual qPCR assay depending on initial input DNA template. First, to assess amplification efficiency, the pooled synthetic DNA standard was serially diluted by a factor of two, covering a resulting qPCR input DNA range from 50 down to 0.39 ng. Based on each assay’s standard curve (triplicate analysis), we obtained a well-performing PCR efficiency (98 % - *hTERT* Short, 99 % - *hTERT* Long, 94 % - *LINE1* Converted, 90 % - *LINE1* Genomic) and R^2^ > 0.98 in all assays (**Fig 1C**). Importantly, the single- and multi-copy assays were performing comparatively, even at the lowest input. Yet, as expected the accuracy of DNA copy quantification decreased with decreasing DNA input (**Fig 1G**). More specifically, we provide the first indication that quantification is sensitive and possible even down to 0.78 ng input DNA without remarkably sacrificing accuracy (81 % - *hTERT* Short, 80 % - *hTERT* Long, 75 % - *LINE1* Converted, 90 % - *LINE1* Genomic). Additionally, we assessed the performance of the IPC using data from six experiments, demonstrating that its detection is expected after the 27^th^ PCR cycle in the absence of any inhibition (**Fig 1D**). Finally, we showed that the synthetic DNA is stable and can be used for up to two days after preparation without compromising performance (**Fig 1F**), which can be practical for high-throughput analysis.

### Initial method validation using standard criteria

During the next phase of development, we aimed to test the above described qBiCo-v1 assay in terms of standard method performance criteria based on the three main parameters it measures: i) global BC efficiency (**S1 Fig**), ii) BC-DNA concentration (**S2 Fig**), and iii) BC-DNA fragmentation (**S3 Fig**). To this end, we employed one of the most popular BC kits (EZ DNA methylation kit, Zymo Research) based on optimal conditions (200 ng gDNA input) to avoid introducing further bias. All analyses were performed at the qPCR quantification level and the associated data can be found in **S2 Table**.

Global BC efficiency measurements were highly repeatable based on six replicates (average SD = 0.37 %) at least for highly converted samples (BC > 85 %) (**S1 FigA**). When measurements were replicated in two additional qPCR experiments, the reproducibility was very high, but we noticed possible small run-to-run bias (3 – 4 % lower for one sample) (**S1 FigB**). Additionally, we could successfully measure the expected BC efficiency even down to at least 0.24 ng of BC-DNA input (**S1 FigC**). Strikingly, the quantification of this parameter was very robust and did not significantly change even in the presence of PCR inhibitors at extremely high levels (800 µM hematin) (**S1 FigD**). On the other hand, artificial initial gDNA degradation using UV exposure, even for 1 min, strongly impacted BC efficiency measurement (> 10 % lower) (**S1 FigE**). Importantly, we can accurately and reproducibly determine the right portion of BC-DNA in converted/non-converted mixtures, indicating strong linearity of our detection (R^2^ = 0.994) (**S1 FigF**).

Results were very similar regarding these performance parameters for the measurement of BC-DNA concentration (**S2 Fig**). Our approach seems to be able to relatively accurately detect BC-DNA down to 0.195 ng/µl (**S2 FigC**), based on a linear fashion (R^2^ = 0.989, **S2 FigF**). Additionally, given that this is the only parameter that other researchers have previously attempted to quantify based on existing methods such as fragment analysis (Bioanalyzer), spectrophotometry (Nanodrop), fluorometry (Qubit) (see details in Materials and Methods), we were also interested in comparing qBiCo’s performance against these previously used three methods, particularly at two different gDNA amounts (optimal: 200 ng and suboptimal 16 times less: 12.5 ng). As shown in **Fig 2**, our qBiCo-v1 assay surpasses in performance and quantification abilities all three methods, which tend to underestimate the BC-DNA quantity with limited reproducibility.

**Fig 2.**
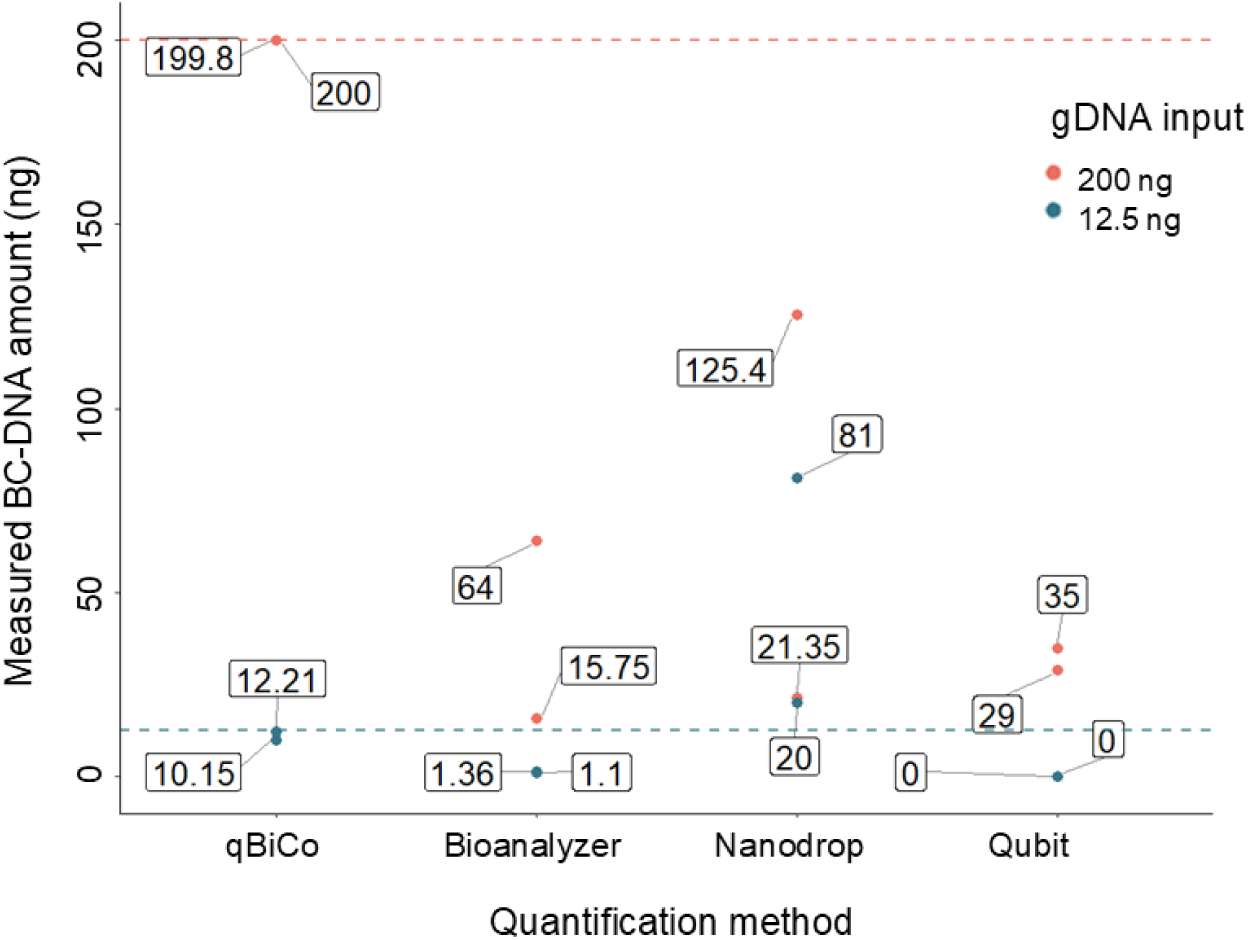
Comparison of BC-DNA measurement capabilities between qBiCo-v1 and other currently used but non-specific methods. Measurements were performed in duplicate. Dotted lines show the expected DNA amounts. Information on protocols used are found in the Methods. BC: Bisulfite conversion; gDNA: Genomic DNA.

However, measuring BC-DNA fragmentation levels turned out more challenging (**S3 FigA-E**). Not only was there large variation (> 25 %) in the detected BC-DNA fragmentation ratio between technical replicates of the same sample within the same experiment, but also across qPCR runs, making this parameter less stable and reproducible. Given that the BC-DNA fragmentation levels we detected in our samples even under optimal conditions were very high (60 – 90 %), the effect of PCR inhibitors and DNA degradation was more severe. Despite these difficulties, we found the performance of qBiCo-v1 during this initial validation process promising.

### Performance evaluation of commercial BC kits

Next, we aimed to test its applicability by assessing popular BC protocols used by epigenetic researchers. To achieve this aim, we employed our method to assess the performance of ten different commercially available BC kits using five decreasing gDNA inputs, (200 ng to 1 ng) using three independent commercial DNA samples. Details on employed BC kits can be found in **Table 1**, while associated data are shared in **S3 Table**. Overall, there was striking evidence that the BC process can differ substantially between commercial kits and their protocols, despite the manufacturers’ promises of > 99 % BC effiency even at minute amounts (< 1 ng). Regarding BC efficiency (**Fig 3A**), all kits except kit 10 (Active Motif) resulted in levels > 90 % using optimal gDNA inputs (200 ng). Kit 10 performed really poorly for all samples and DNA amounts, with BC rate ranging from 2 to 80 % indicating a systemic issue in the BC process. As expected, the efficiency of BC is more variable and reduces with decreasing amounts of input DNA. However, there are certain kits that based on our work performed extremenly well (> 95 %) even at the lowest amounts of input DNA (1-10 ng), particularly kit 3 (Diagenode), kit 5 (Sigma Aldrich) and kit 6 (Qiagen). Moreover, it was common to observe that often only one of the three DNA samples in each input resulted in decreased BC performance, which impacted the observed variance for a particular kit; for example, for kit 4 (Promega).

**Fig 3.**
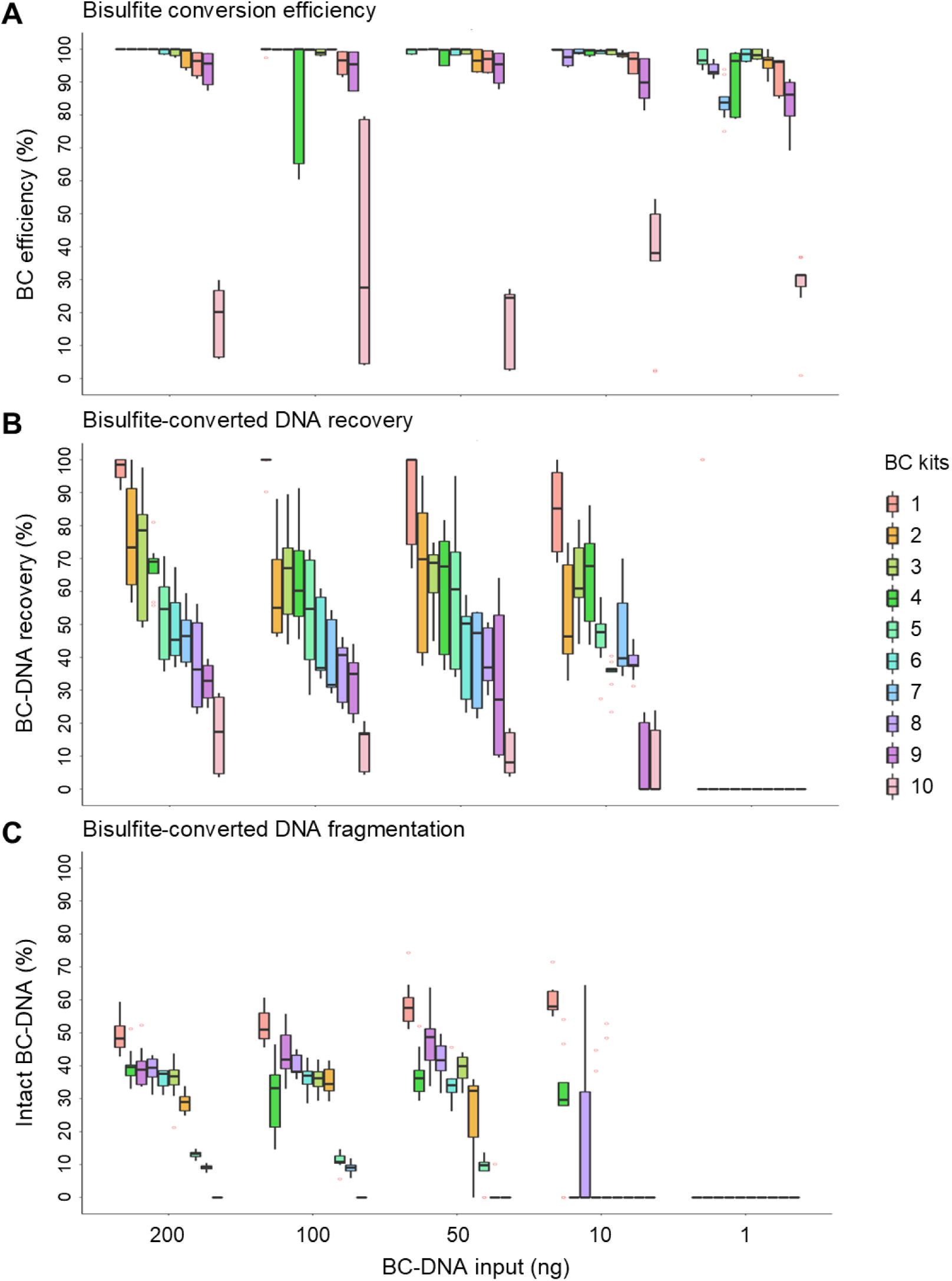
Testing of ten commercial BC kits based on qBiCo-v1 indices. (A) BC efficiency, (B) BC-DNA recovery and (C) BC-DNA fragmentation levels (≥ 235bp) depending on the input DNA amount used for the BC of three commercial DNA standards. Red circles indicate outliers > ± 2SD. Information on kits used are found in Table 1. BC: Bisulfite conversion.

**Table 1.** Information on BC kits used during the testing of qBiCo-v1 method.

| **Kit No** | **Manufacturer** | **Kit name** | **Incubation protocol** |
| --- | --- | --- | --- |
| 1 | Zymo Research | EZ DNA Methylation | 14 - 16 hours at 50 °C |
| 2 | Thermo Fisher Scientific | EpiJET BC | 10 min at 98 °C /<br>150 min at 60 °C |
| 3 | Diagenode | Premium Bisulfite | 8 min at 98 °C /<br>60 min at 54 °C |
| 4 | Promega | MethylEdge® BC System | 8 min at 98 °C /<br>60 min at 54 °C |
| 5 | Sigma Aldrich | Imprint® DNA Modification | 90 min at 65 °C |
| 6 | Qiagen | EpiTect Bisulfite | 5 min at 95 °C /<br>25 min at 60 °C /<br>5 min at 95 °C /<br>85 min at 60 °C /<br>5 min at 95 °C /<br>175 min at 60 °C |
| 7 | Abcam | Fast BC | 20 min at 95 °C |
| 8 | Analytik Jena | innuCONVERT | 45 min at 85 °C |
| 9 | Epigentek | Methylamp DNA modification | 90 min at 65 °C |
| 10 | Active Motif | BC | 30 sec at 95 °C /<br>20 min at 58 °C |

Regarding BC-DNA recovery, the results were even more negatively surprising, particularly because most kit manufacturers promise at least 80 % recovery. **Fig 3B** showcase a remarkable variability among kits in terms of the amount of eluted BC-DNA, regardless of the initial gDNA input. While there are certain kits (1 - 3: Zymo Research, TFS, Diagenode) that on average suffered from 0 – 30 % DNA loss at optimal amounts, there are others (5 - 7: Sigma Aldrich, Qiagen, Abcam) that constantly cause ∼50 % DNA loss. Similarly, certain kits (9 - 10: Epigentek, Active Motif) degrade or lose > 70 % of the available DNA. We also noted that measured BC-DNA recovery is highly variable among samples within a kit, but the trend per kit is similar across DNA amounts, indicating a systematic performance. It is worth noting that no data were obtained when only 1 ng of gDNA was converted.

Finally, it is well-known that the BC process causes DNA fragmentation, but it has been challenging so far to measure it in a quantitiative way. Here, we measured detection differences between the *hTERT* Short (85 bp) and *hTERT* Long (235 bp) assays. Even at this short length, we noticed > 50 % less fluorescent signal for the longer fragments, even under optimal conditions (200 ng input) (**Fig 3C**). Observations and trends among kits were similar down to 50 ng, with kit 1 (Zymo Research) outperforming the other kits. Given the severe fragmentation, we failed to detect any signal for the longer fragment even at the 10 ng input. In combination with our previous observations during initial validation, this qBiCo-v1 parameter is by far the least sensitive.

### Scaling-up towards a small-scale technology prototype

Based on our experience so far and aiming to advance the accuracy, sensitivity and applicability of our method, we followed several optimization steps to improve its functionalities and to make it more user-friendly and cost-effective. Our overall aim was to create an improved version (qBiCo-v2) towards a small-scale technology prototype that can form the basis of a future commercial product.

First, we aimed to improve the individual qPCR assay design. Due to its poor performance, we decided to redesign the *hTERT* Long assay used to obtain the BC-DNA fragmentation index. However, despite redesigning to amplify a slightly shorter fragment at another region of the *hTERT* gene and further adjusting the primer/probe ration in the multiplex qPCR reaction, the performance of the redesigned assay was not satisfactory (efficiency of 61 %) (**S4 FigA**). Therefore, we decided to completely replace this assay with a similar one (222 bp) using another single-copy gene (*TPT1*) that resulted in a more stable and efficient amplification (efficiency of 91.8 %) (**S4 FigB**). Additionally, to boost the amplification of both our single-copy assays, we tested the effect of reducing the copies of the IPC assay (originally 6,000 copies per reaction). As a result, we obtained robust IPC detection down to 3,000 copies without compromising the measurement of PCR inhibition (**S4 FigC**).

Moreover, to optimize the amplification efficiency of all assays, we examined multiple parameters. For example, we further adjusted the primer/probe mix concentrations, the amplification program by changing the denaturation time and temperature, and we examined the effect of adding betaine or DMSO in the reaction, besides MgCl_2_ and BSA that are already contained in the assay. However, the extra additives led to instability of the *TPT1* Long fragment (data not shown) and therefore, we did not proceed further. Finally, to increase cost-effectiveness, we also reduced the qPCR reaction volume by 50 % (10 µl) without compromising performance. The optimized qPCR conditions and updated oligo sequences (qBiCo-v2) are presented in the **S4 Table**. Importantly, we aimed to adjust the composition and preparation strategy of our synthetic DNA standard to make it more representative, considering our observations during the BC kit testing and the usual gDNA input range researchers use. Our goal was to create a synthetic standard that resembled as much as possible the detection and assay detection ratio of samples from the ‘field’. To capture the entire spectrum of expected eluted BC-DNA variability, we created a testing ‘master’ sample by converting a commercial methylated DNA standard using five different amounts and a much larger range of BC kits (n = 16). Following a thorough tuning strategy and multiple optimization qPCR runs, we settled with a new synthetic DNA standard mix (**Table 2**) and dilution strategy (**S5 Table**) with higher robustness and less variability among technical replicates (**S5 Fig**). More specifically, we increased the number of dilutions and pipetted volumes during preparation using TE buffer (pH = 8.0) instead of water. These adjustments increased the stability of the fragments upon storage and subsequently improved the metrics of the standard curves.

**Table 2.** Composition of the updated synthetic DNA standard in qBiCo-v2 method.

| qPCR assay | Number of copies (DNA molecules) |  |  |  |  |
| --- | --- | --- | --- | --- | --- |
|  | StdA | StdB | StdC | StdD | StdE |
| *hTERT* Short | 6000 | 2000 | 667 | 222 | 74 |
| *TPT1* Long | 6000 | 2000 | 667 | 222 | 74 |
| *LINE1* Converted | 384000 | 128000 | 42667 | 14222 | 4741 |
| *LINE1* Genomic | 11400 | 3800 | 1267 | 422 | 141 |
| Synthetic IPC | 3000 | 3000 | 3000 | 3000 | 3000 |

Beyond the wet lab, we also aimed to improve and streamline the data analysis part. First, instead of originally estimating the nanograms of DNA involved, we switched our approach to calculating the actual copy numbers for each detected gene fragment. Second, to enable large-scale analysis in shorter time, we developed a user-friendly spreadsheet file based on formulas and macros for a semi-automatic way of analysing the qPCR data (**S1 File**). The data analysis pipeline also includes the automatic calculation of both qPCR performance metrics and qBiCo parameters per sample. Third, we developed a strategy to systematically detect data outliers, applicable to both duplicate and triplicate analysis.

### Applicability to other real-time qPCR platforms

During the next phase of development, to enable uniform implementation in other laboratories we aimed to transfer and test the qBiCo-v2 method across real-time qPCR platforms of several manufacturers. To achieve our goal, beyond using our own BioRad CFX96 instrument, we collaborated with other researchers to employ five additional commonly encountered qPCR instruments: BioRad CFX384, TFS 7500 Fast, TFS QuantStudio 5 (QS5), TFS QuantStudio 7 Flex (QS7 Flex), and Qiagen RotorGene Q.

Overall, we successfully adopted our qPCR protocol in all five platforms, by simply adjusting the dyes to fit each instrument’s spectra and the qPCR reaction volume (**S6 Table**) and our analysis pipeline to fit the different software used for the experiment set-up and data extraction (**S2 - S3 Files**). Additionally, we evaluated its performance based on the obtained Cq values and relative fluorescent units (RFU) when running the same qPCR run on all instruments (**S7 Table, S6 - S7 Fig**). As expected, there were substantial differences in terms of fluorescence intensity, which can be explained by instrument-to-instrument specifications as each instrument works with different (log) scales. Nevertheless, Cq values were comparable across instruments for each qPCR assay (**S7 Table**). Importantly, we compared the PCR efficiency and goodness-of-fit of the obtained standard curves for each of the four qBiCo-v2 qPCR assays (without IPC) (**Fig 4**). We detected subtle differences in PCR efficiency, particularly evident for the *TPT1* Long assay for most instruments, and for the *LINE1* assay for the TFS instruments, the latter of which is detected with a proprietary dye (ABY™) (**Fig 4A, S6 Table**).

**Fig 4.**
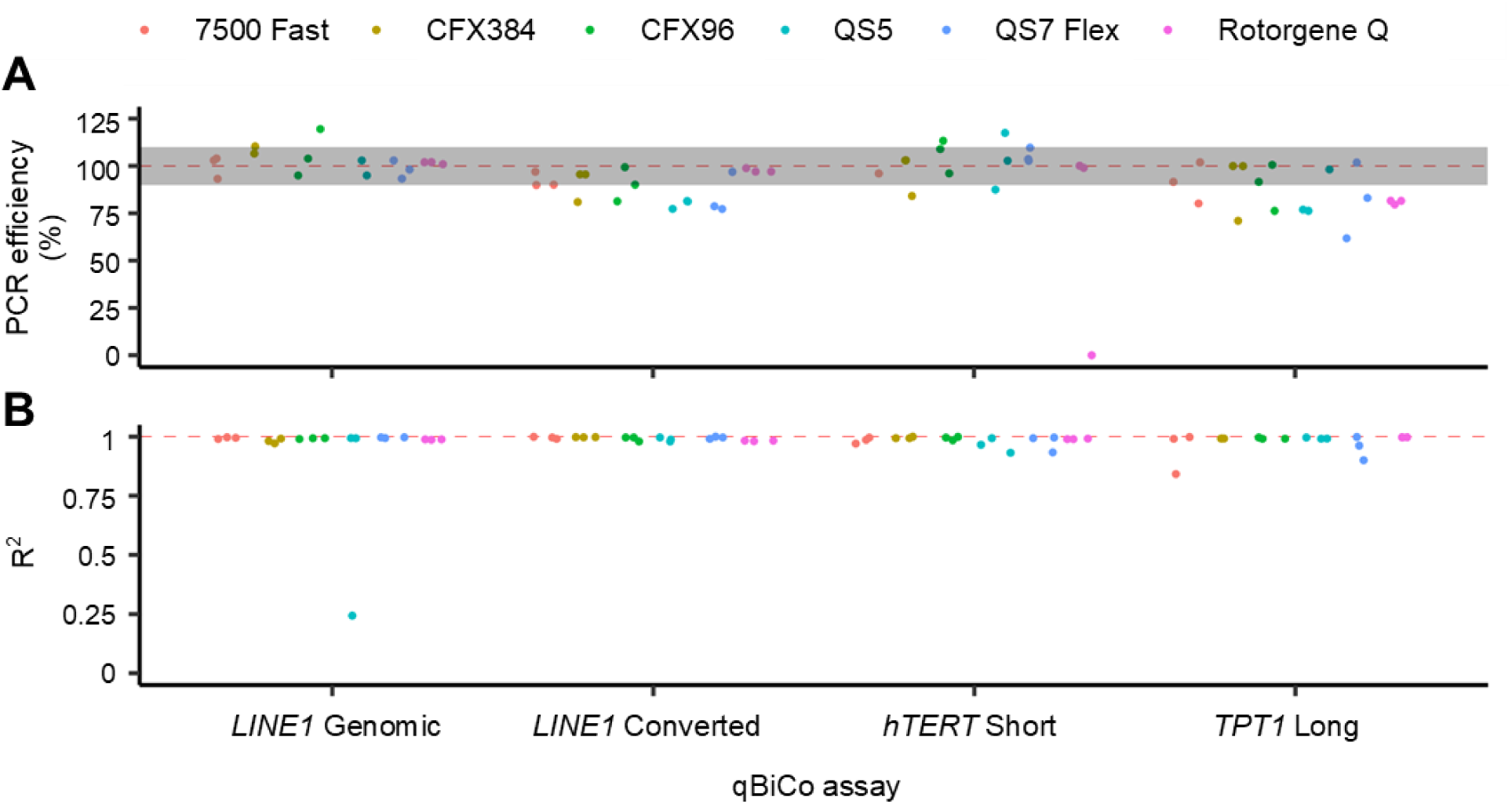
Comparative qBiCo-v2 assay performance on six qPCR instruments. (A) PCR efficiency (%) and (B) goodness-of-fit values (R^2^), of each qPCR assay of qBiCo-v2. Data are included from the initial method transfer runs on six qPCR systems from three biotechnology manufacturers (BioRad, Thermo Fisher Scientific, Qiagen). Dashed red lines indicate ideal performance measurements, while the grey box indicates acceptable range by standard qPCR guidelines (PCR efficiency between 90 – 110 %).

### Instrument-specific developmental validation

To further establish the capacity, usability, and possible instrument-specific limitations of our qBiCo-v2 method, we decided to conduct a small-scale validation study per qPCR platform. Our methodology to assess different performance evaluation parameters is described in Materials and Methods, while all data per instrument and statistical evaluation can be found in **S8 and S9 Tables**, respectively.

First, we evaluated the ability of accurately assessing BC-DNA at two different gDNA inputs: 50 and 5 ng, with an expected maximum of 5 and 0.5 ng/µl input into qBiCo, respectively. We chose these amounts based on our previous experience optimizing this method, but also aiming to go lower with the gDNA input and evaluating the applicability of qBiCo-v2. For BC efficiency, we observed concordant results between instruments: average BC between replicates ranging 0.987 – 0.999 at 50 ng and 0.940 – 0.994 at 5 ng (**Fig 5A**). Nevertheless, for BC-DNA concentration, we observed a larger variation between instruments for both input DNA amounts: average BC-DNA recovery between replicates ranging 0.493 - 2.252 at 50 ng and 0.443 - 1.409 at 5 ng (**Fig 5B**). For three instruments (7500 Fast, CFX384 and CFX96), there were statistically significant differences between the two amounts (p < 0.01, **S9 Table**). It was also clear that the 7500 Fast instrument overestimated the concentration of BC-DNA compared to the other instruments used. Similarly, for BC-DNA fragmentation, we observed a similar trend with BC-DNA recovery; average BC-DNA fragmentation index between replicates ranging 0.861 - 1.116 at 50 ng and 1.026 - 6.412 at 5 ng (**Fig 5C**). In this case, the CFX96 instrument seems to have overestimated the fragmentation of BC-DNA compared to the other instruments applied.

**Fig 5.**
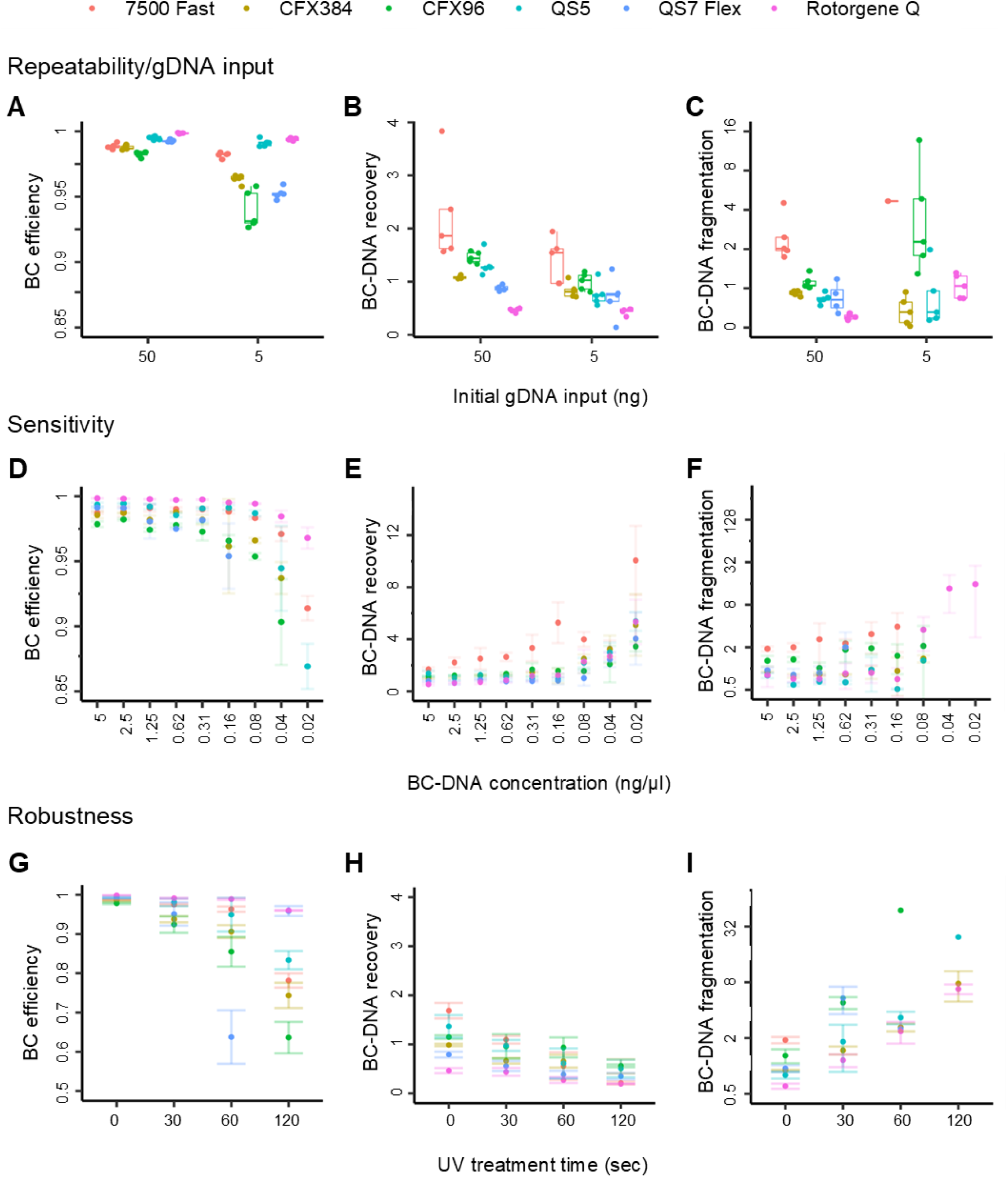
qBiCo-v2 assay validation using synthetic DNA standards on six qPCR instruments. Testing (A-C) repeatability at two DNA inputs, (D-F) sensitivity down to 0.02 ng/µl, and (G-I) robustness at different UV exposure times, per qBiCo index and qPCR instrument (colour-coded). All qBiCo indices are presented in ratios. BC: Bisulfite conversion.

Next, we evaluated the ability of accurately assessing BC-DNA at decreasing qPCR DNA inputs ranging from 5 down to 0.0195 ng. To this end, we employed the BC-DNA samples based on 50 ng of gDNA input, to minimize the effects of the BC process itself. Overall, results and trends among instruments were similar; however, it was evident that adding less BC-DNA into the qBiCo reaction can significantly alter the obtained indices. This was particularly clear for measuring BC efficiency (p < 0.01) and BC-DNA concentration (p < 0.0001), for most instruments (**S9 Table**). Specifically for BC efficiency, measurements were significantly affected below 0.1563 ng BC-DNA (**Fig 5D**). Similarly, BC-DNA concentration measurements were significantly impacted at amounts lower than 0.08 ng BC-DNA, for most instruments (**Fig 5E, S9 Table**). In line with our observations described above, the 7500 Fast instrument seems to overestimate the BC-DNA concentration. Finally, for BC-DNA fragmentation, and as expected from our previous experience, we failed to obtain sufficient data below 0.1563 ng BC-DNA, except for RotorGene Q which seems to be the most sensitive for this qBiCo index (**Fig 5F**). Overall, considering all parameters, we can conclude that 150 pg input DNA is the limit of detection of qBiCo-v2.

Importantly, using this sensitivity threshold, we tested the reproducibility of the qBiCo-v2 method across qPCR platforms. The results were very promising for all decreasing amounts (**S8 Fig**, **S9 Table**). Strong statistically significant differences were only detected for the CFX96 instrument for all three qBiCo indices (p < 0.001) and for the 7500 Fast instrument in terms of BC efficiency (p < 0.0001). Moreover, we evaluated the ability of accurately assessing BC-DNA in compromised, artificially degraded samples using UV treatment. As expected, the measurements using all instruments were significantly affected (**Fig 5G-I**, S9 Table), particularly for longer UV exposure times (60 - 120 seconds). Finally, we evaluated the specificity of BC-DNA evaluation using qBiCo-v2 using a range of non-human DNA samples (**S8 Table**). Unsurprisingly, we obtained false positive results for *Rhesus macaque* (monkey), particularly when measuring global BC efficiency, which can be explained since we target the evolutionally conserved region of the *LINE1* repeat.

## Discussion

BC has been the golden standard for decades, allowing researchers to easily translate methylation differences into sequence variation that can be detected using standard genetic technologies [14], each with their own strengths and weaknesses [28]. So far, these techniques have been used for methylation biomarker discovery and validation, uncovering method-specific methylation biases among targeted techniques [29] and between genome-wide and PCR-based approaches [30], introduced either experimentally or during data analysis [31]. Currently, as we move towards larger global research efforts, more difficult and scarce templates, and implementation in the clinic, the need for methylation assay standardization becomes prominent. Currently, the lack of proper standardization stands out as the most important contributor to DNA methylation variability [32]. Yet, while laboratories validate methylation assays post-BC [33], the performance of BC remains a black box. In genome-wide analysis, BC control probes are only assessed post-analysis after sample processing [34], not as QC for downstream decision making to exclude inappropriately converted samples. In targeted analysis, BC-DNA is enriched by using converted-specific primers or probes, making some think a QC is not necessary. However, different BC-DNA inputs in the PCR and storage conditions that can cause further fragmentation, were among the most significant factors for large methylation variation [32]. Measuring the BC-DNA input is also essential in statistically interpretating methylation data, to map stochastic events that are particularly relevant in low amounts (< 1 ng) of BC-DNA [35].

While usually scientists assume its successful performance, there are a few studies evaluating BC kits [18–22], already highlighting great variance using targeted, non-specific approaches. Motivated by these small-scale efforts, we aimed to offer the community an innovative and complete technological solution for global BC performance assessment, which can also potentially form the basis of a future commercial product. Our goal was to establish a method and prototype tool that can simultaneously assess both the quality and quantity of BC-DNA prior to downstream analysis using a bench-top instrument commonly accessible in molecular biology laboratories. It is important that the BC-QC solution is simple, cost-effective, reliable and robust, hence we opted for a method based on real-time qPCR, which is also often used for the QC of gDNA. Additionally, qPCR enables the simultaneous amplification of multiple small DNA regions in one reaction, hence the assessment of several QC parameters at the same reaction.

Compared to previous studies that have assessed BC-DNA recovery and fragmentation, to the best of our knowledge we are the first to build a QC solution that measures a global BC efficiency index. We did this by analysing the genomic and converted versions of an evolutionally conserved, (non-CpG) C-rich region of a repetitive element (*LINE1*). Recently, Hong and Shin (2021) developed another multiplex qPCR-based system (BisQuE) with a similar goal in mind, but their BC efficiency measurement is based only on one cytosine of a single gene intron [36], which can highly bias the assessment. In contrast, via the *LINE1* Converted probe we target five cytosines across ∼180-200 different *LINE1* regions in the genome per sample, making our BC efficiency measurement much more representative compared to targeting a single-copy locus. While we expect that each human genome might display unique genetic variants in some of these *LINE1* copies, we do not envision this as problematic since the BC efficiency measure is based on a ratio of hundreds of targets, hence is quite robust to individual copy effects. On the other hand, the choice of targets for our single-copy assays was not as straightforward. We opted for the *hTERT* gene, since we also employ this target for gDNA QC assessment in our laboratory (Quantifier^™^ Duo DNA quantification kit, TFS). This allows us for avoiding locus-specific biases when comparing quantities obtained before and after BC. Of course, *LINE1* and hTERT are just possible options we could have used, and in principle the use of any common repetitive element and single-copy gene respectively, would work.

Altogether, qBiCo-v1 showed promising performance, particularly in comparison to other non-specific methods commonly used as QC. While these methods are not BC-DNA-specific, they take advantage of its nature being mostly single-stranded for their quantification. Despite having only preliminary data, we successfully showcased qBiCo’s superior performance. However, there were indications of low sensitivity and robustness issues with the *hTERT* Long assay, that drove the overall limit of detection of qBiCo-v1 at 780 pg BC-DNA per µl reaction. At first glance this looks very little, but if we consider that BC-DNA is usually eluted in 10 µl and that BC-DNA recovery is rather low (often < 50 %), this would translate into at least 15 ng gDNA into BC. On the other hand, likely due to the multi-copy nature of *LINE1* assays, the measurement of BC efficiency was very robust even at extreme inhibition and fragmentation levels. However, caution is required as we observed low specificity with the *LINE1* Genomic assay, which was often amplified earlier than anticipated. Finally, from our initial validation efforts it was clear that the fragmentation index was the weakest assay in qBiCo, where we often obtained no data, even if we only target a 235 bp long fragment.

Eventually, we were able to highlight qBiCo’s need and potential in the field, when using a wide range of BC kits as part of research and diagnostics assays. The employed kits were selected from ten different manufacturers. They display similar protocols concerning the basic structure and steps in the BC process – sodium bisulfite incubation, DNA binding, desulfonation, washing and elution of BC-DNA, which allowed for a fair comparison. Nevertheless, there is variance within some steps i.e., incubation length and temperature, spin column width and size. Overall, our findings were striking. First, it is clear that a promised BC efficiency of > 99 % is not always achieved. While we uncovered systematic issues with some kits, others achieved consistent successful performance even down to 10 ng. Yet, qBiCo could pick out individual sample failures and differences. One question that remains to be determined by the community is the threshold one should choose for successful BC efficiency assessment. Should it be 90, 95 or 99%? Lastly, it was clear that results were worsened when treating 1 ng of BC-DNA. Detected BC efficiencies were much lower, which can be driven not only by the BC process itself, but also by the capabilities of qBiCo (in the best-case scenario amounts translate to 100 pg input into each reaction).

Additionally, our assessment in terms of BC-DNA recovery and fragmentation is in line with recent work by others. Using Qubit and testing 12 BC kits, average BC-DNA recovery was measured between 26.6 and 88.3 % using kit-specific optimal gDNA inputs (135 ng to 2 µg) [19]. On the other hand, using qPCR-based BisQuE Hong and Shin previously showed that average BC-DNA recovery at 50 ng input could range between 18.2% and 50.6% [36]. Results are similar also in terms of BC-DNA fragmentation, even though we acknowledge that it is difficult to compare quantitatively as previous methods use different approaches, indices and fragment lengths. Here, we show that even with the best performing kit, the intact BC-DNA portion (at least at 235 bp level) is no more than 50-60 % of the sample. Previous observations using qPCR have reported several Cq differences between their short and long fragments [19]. Also, in another previous study, BC-DNA degradation indices were measured between 1.5-2.5 [37], which agree with our findings. We cannot exclude locus-specific biases in these assessments, driven for example by DNA accessibility, 3D DNA structures or binding proteins, but overall BC damages DNA substantially. This might be extra challenging when applying qBiCo to low-quality/quantity BC-DNA samples like cell-free or forensic-type DNA. For such applications, the right choice of BC kit is critical. In a recent study on cell-free tumour

DNA, the EpiTect kit (QIAGEN) performed best in terms of BC-DNA recovery and fragmentation based on digital PCR and Bioanalyzer, respectively [38]. Based on these and our findings, every kit performs differently having their own strengths and weaknesses. Therefore, there is a need for every lab to thoroughly validate their BC kit of choice for their own application, for which qBiCo offers a complete solution. This concerns not only well-established commercial BC kits, but also newly developed, not-yet-commercial BC methods like rapid ones combining DNA extraction and conversion [39] or based on microfluidics [40] as well as ones using enzymatic approaches [41].

To enable widespread adoption of qBiCo, we continued our technology development towards building a small-scale prototype. Following a set of functionality improvements, we built an improved our method (qBiCo-v2). Despite replacing our long assay and targeting a new gene (*TPT1*), we still found its performance weak and not robust enough. Future work could explore alternative solutions in a larger screening approach. For example, an idea is to offer a more sensitive and robust approach for BC-DNA recovery and fragmentation also via repetitive elements [42]. Still, caution should be taken in the chosen length of the long assay to make it meaningful enough for downstream applications. Moreover, we updated our approach of creating the synthetic standard from mixing to preparation and improved it more in terms of quantitative power and applicability. We ensured that the synthetic standards were amplified and detected at similar time points with BC-DNA samples considering 16 different BC kits and 5 different amounts. Nevertheless, we still observed substantial batch effects, so it is important to find ways to standardize the production of the synthetic standard in the future. Importantly, we semi-automated the analysis from the raw qPCR data to the qBiCo indices to enable for a faster and normalized approach.

In addition, to further enable implementation, we tested the qBiCo-v2 protocol in several commonly used instruments from three manufacturers. We were pleased by how smooth this transfer was as the only necessary adjustments were driven by instrument-specific limitations in terms of compatible dyes (TFS), number of channels (BioRad) and minimum reaction volume (Qiagen). In the future, using our approach we are confident that scientists can easily extend the transfer to additional instruments, for example the LightCycler^®^ platforms (Roche). Generally, our instrument-specific validation results aligned with qBiCo-v1, with detectable variance within and between BC-DNAs, qPCR runs, qBiCo indices and qPCR instruments still being observed. We can explain these by simple errors during sample preparation and storage as well as micro-pipetting, but systematic, instrument-specific limitations were also uncovered. Nevertheless, our statistical evaluation should be considered cautiously due to the small sample size and missing data particularly in the case of the *TPT1* Long fragment and robustness experiments. All in all, based on our empirical observations, we provide useful guidance to researchers that possess these instruments and wish to perform and/or extend these validation procedures in their own laboratory. Future implementation of qBiCo in the (high throughput) end-user environment will further highlight its true potential and usefulness.

## Conclusion

In 2017, Lind and van Engeland highlighted: “Only when including quality and standardization at every level of DNA methylation analyses, will we be able to achieve the robustness to independently validate DNA methylation analyses and to compare multiple methylation studies in systematic reviews. This is the only way to more efficiently develop future DNA methylation-based biomarkers.” [43]. This is particularly true in specialized applications like forensics, where data standardization and scrutiny are of paramount importance, currently hindered by the lack of BC assessment [44]. In this study, we built the first-of-its-kind method and prototype tool for global BC performance assessment based on qPCR. qBiCo can detect several parameters of a BC-DNA sample: efficiency, recovery, fragmentation and inhibition. Here, we demonstrated our technology development: from basic method formulation (TRL-1) to technology concept (TRL-2) to critically improved, established proof-of-concept (TRL-3) to testing and validation of a small-scale prototype (TRL-4). We conclude that we achieved our aim of offering a BC-QC solution to the epigenetic community, providing evidence of its performance across different samples, BC-DNA inputs, BC kits, qPCR instruments and other conditions. We also present a thorough critical method assessment and suggestions for further improvement and development. Motivated by our findings, we call epigenetic researchers to shift their perspective from assumption to empirical assessment with regards to BC kit performance and integrate qBiCo as a QC step in their methylation assays regardless of downstream analysis. We also hope to inspire others to develop similar methods for other types of data, i.e. BC-RNA data in epitranscriptomics [45].

## Materials and Methods

### DNA sample preparation

During the different phases of technology development, one or more commercially available DNA samples were used: the human high methylated gDNA standard (100ng/µl) (EpigenDX, USA) (qBiCo-v1 validation and BC kit testing), the human Methylated DNA (250 ng/μl) (Zymo Research, USA) (qBiCo-v1 BC kit testing and qBiCo-v2 functionality improvement) and the Quantifiler™ THP DNA Standard (100 ng/μl) (TFS, USA) (qBiCo-v1 BC kit testing). For the final phase of instrument-specific validation (qBiCo-v2), two commercially available biobank human blood samples (Sanquin, The Netherlands) were employed, together with a range of commercially available and already extracted non-human DNA samples from different species (Novagen, USA): *Felis catus* (cat), *Canis lupus familiaris* (dog), *Mus musculus* (mouse), *Rattus* (rat), *Gallus gallus domesticus* (chicken), *Sus scrofa domesticus* (pig), *Bos Taurus* (cow), and *Rhesus macaque* (monkey). gDNA isolation was performed using the QIAamp^®^ DNA Investigator kit (Qiagen, Germany), following the whole blood protocol. All human DNA samples included in this study were quantified prior to BC to assess for their DNA quantity and quality, using the Quantifiler^™^ DUO DNA quantification kit (TFS).

### BC-DNA sample preparation

For qBiCo-v1 validation, 200 ng of the human high methylated gDNA (100 ng/µl) (EpigenDX, USA) were bisulfite-converted using the EZ DNA methylation kit (Zymo Research) using the manufacturer’s instructions. During qBiCo-v1 testing, all three DNA standards mentioned above underwent BC in five different DNA amount - 200, 100, 50, 10, 1 ng using a total of ten commercially available BC kits to compare their performance (**Table 1**). The selected BC kits are popular and belong to ten different companies. All kits are optimized to convert DNA within the range of 100 ng - 2 µg, with a suggested optimal DNA amount set at 200 ng. BC protocols and incubation times were followed according to manufacturer’s instructions and BC-DNA samples were eluted in 10 µl.

During functionality improvement towards qBiCo-v2, we prepared and employed a set of ‘master’ samples that realistically resemble BC-DNA quantity/quality, often seen in real-world analysis. For this, and to capture all possible variation across commercially available products, we used 16 different kits according to manufacturer’s instructions: EpiTect Bisulfite kit (Qiagen), EpiTect Fast bisulfite kit (Qiagen), EZ DNA Methylation kit (Zymo Research), EZ DNA Methylation Gold kit (Zymo Research), BC kit (Active motif, Belgium), Methylamp DNA modification kit (Epigentek, USA), EpiJET BC kit (TFS), MethylCode™ BC kit (TFS), Premium Bisulfite kit (Diagenode, USA), Fast BC kit (Abcam, England), EpiMark^®^ BC kit (New England Biolabs, USA), DNA BC kit I (Biovision, England), Imprint^®^ DNA Modification kit (Sigma-Aldrich), CpGenome Direct Prep Bisulfite Modification kit (Merck Millipore, USA), innuCONVERT (Analytik Jena, Germany), MethylEdge^®^ BC System (Promega, USA). To prepare the master samples, the Human Methylated DNA (250 ng/μl) (Zymo Research) was converted using different DNA inputs (200, 50, 20, 5 and 2 ng) per kit and eluted in 50 μl. Then, for each amount, 30 μl of each BC-DNA sample was combined, to form a 480 μl master solution.

For the final phase of instrument-specific validation (qBiCo-v2), the whole blood DNA samples were bisulfite-converted using EZ DNA Methylation kit (Zymo Research). Each sample was converted 20 times using 50 ng and three times using 5 ng of DNA input. For all conversions, BC-DNA was eluted in 10 μl. To ensure the same quantity and quality across instruments, eluates of the same DNA input (e.g., 50 ng conversions) were pooled together in one tube and subsequently aliquoted to separate tubes. These samples were used to assess repeatability, sensitivity, and reproducibility. For sensitivity, the measured BC-DNA samples of 5 ng/µl were serially diluted (2X) up to nine times. For robustness, 5 ng/μl of BC-DNA was UV treated for different time periods, while for specificity, 200 ng of the non-human samples described above were bisulfite-converted.

### BC-DNA evaluation using existing methods

Two BC-DNA samples (initial gDNA input of 200 and 12.5 ng) were evaluated using other existing spectrophotometric and fluorometric methods often employed for BC-DNA quantity/quality assessment. For all reactions 1 µl was used, while for all methods reactions were performed in duplicate. First, we employed a NanoDrop Microvolume Spectrophotometer (TFS) to measure absorbance at 260 nm using the single-stranded DNA mode option. Next, we employed a 2100 Bioanalyzer instrument (Agilent Technologies, USA) using the RNA 6000 Pico kit (Agilent Technologies). Finally, we used the Qubit™ 4 fluorometer (TFS) using the Qubit™ single-stranded DNA Assay kit (TFS). Kit choices were driven by the fact that strands have lost their complementarity making BC-DNA mostly single-stranded and resembling the secondary structure of RNA. For all methods we used the manufacturer’s instructions to calculate the BC-DNA concentration.

### Synthetic DNA standard

Normally, for the quantification of human gDNA a commercially available human DNA standard is used. However, for the quantification of BC-DNA, this is not possible given that we cannot control neither the quantity nor the quality following BC. Hence, we opted for creating a synthetic DNA standard that resembles the composition of human BC-DNA. For this, we designed five artificial double-stranded gene fragments (gBlocks™, Integrated DNA Technologies, Coralville, IA, United States) with DNA sequences identical to the expected generated PCR product sequences when using the proposed assays in each qBiCo version. In qBiCo-v2, additional sequences were added on both the 5’ and 3’ ends of the gBlocks™ to improve production and stability (**S4 Table**).

During qBiCo method formulation (qBiCo-v1), the four different gBlocks™ (except for IPC) were mixed in a ratio that resembles the one naturally found on human DNA based on our design. In other words, both the *hTERT* Short / *hTERT* Long assays as well as the *LINE1* Genomic / *LINE1* Converted assays were each mixed in a 1:1 ratio, while the single- / multi-copy elements were mixed in ∼ 1 : 200 (GRCh37). Practically, the individual gBlocks™ were received in concentration of 10 ng/μl, and then mixed to resemble the expected copy numbers as in 50 ng/μl of human BC-DNA. Subsequently, serial dilutions by factor two were performed to generate the eight standards used in qPCR, with concentration range from 50 to 0.39 ng/μl.

During qBiCo prototype development (qBiCo-v2), we decided to simplify and improve the quantification process by directly working with copy numbers instead of concentration. To translate the given gBlock™ concentration in DNA copy numbers, we used the following equation:

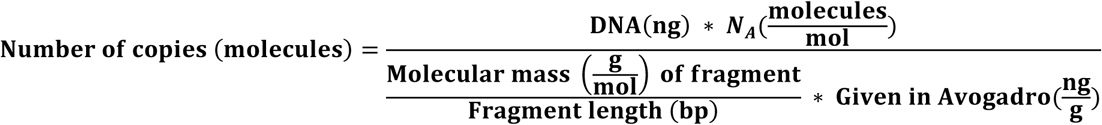

where N_A_ (Avogadro’s number) = 6.02214076×10^23^, and Given in Avogadro = 10^9^ng/g. Additionally, this time we aimed to create a more representative synthetic DNA standard able to quantify what is usually observed in human BC-DNA samples. Hence, the four different gBlocks™ (except for IPC) were mixed in specific empirical ratios following systematic optimization, with a final volume of 100 μl. Practically, the individual gBlocks™ were first diluted four to six times by factor ten and combined accordingly. Subsequently, the synthetic DNA standard mix was diluted by factor two to create the first standard used in qPCR. Four serial dilutions by factor three were performed to generate the rest of the standards. Information on the sequences of the synthetic DNA fragments, copy numbers per qBiCo version and dilution strategy are provided in **S1, S4, S5 Tables** and **Table 2**.

### qPCR assay design

qBiCo is based on the well-established real-time TaqMan^®^-based quantitative PCR technology. It comprises five assays in a multiplex reaction, each of which amplifies either a single- or multi-copy locus to calculate different BC-DNA quality/quantity parameters: global conversion efficiency, concentration, fragmentation and inhibition. Initially, the Ensembl genome browser (https://grch37.ensembl.org/index.html) was used to locate and extract the DNA sequences of the regions of interest in the human genome (GRCh37). Next, the expected BC-DNA sequences were obtained using MethPrimer (https://www.urogene.org/cgi-bin/methprimer/methprimer.cgi) [46]. BC-DNA-specific primers were designed using BiSearch following standard parameters (including no CpG in the primer sequence when possible) [47]. Fluorescently labelled TaqMan^®^ probes were manually designed to target converted C-rich areas and considering instrument-specific requirements. Lastly, Autodimer was employed to check for hairpin and primer-dimer formation [48]. Details on primer/probe sequences, fluorophore labelling for each probe, and length of each fragment per qBiCo version are provided in **S1** and **S4 Tables**.

### qPCR protocol

The five assays were amplified simultaneously in a qPCR reaction using the oligo sequences and conditions described in **S1 Table** (v1) and **S4 Table** (v2). Unless mentioned differently, all reactions were performed in Hard-Shell^®^, thin-wall PCR 96-well plates (BioRad, USA) in a CFX96 Touch™ Real-Time PCR Detection system (BioRad).

For qBiCo-v1 the PCR reaction was optimized in a final volume of 20 µl containing: 10 µl of 2x EpiTect^®^ MethyLight qPCR reagent (Qiagen), 2 μl of 25 mM MgCl_2_ (Applied Biosystems, USA), 0.8 μl of 20 mg/ml BSA (BioLabs, USA), 3.2 μl of nuclease-free water, 2 μl of primer/probe mix, 1 μl of diluted IPC-gBlock™ (1 ng), and 1 μl of BC-DNA template. The cycling conditions for this qPCR consisted of polymerase activation and denaturation (95 °C, 5 min), followed by 33 cycles of denaturation (95 °C, 15 s), annealing (56 °C, 30 s) and extension (60 °C, 70 s). Nuclease-free water was used as negative control. In most cases, qPCR reactions were performed in triplicate.

For qBiCo-v2 the PCR reaction volume was decreased to 10 μl, containing 5 µl of 2x EpiTect® MethyLight qPCR reagent (Qiagen), 1 μl of 25 mM MgCl_2_ (Applied Biosystems), 0.2 μl of 20mg/ml BSA (BioLabs), 1.3 μl of nuclease-free water, 1 μl of primer/probe mix, 0.5 μl of diluted IPC-gBlock™ (**Table 2**), and 1 μl of BC-DNA template. For experiments performed using QS 7 Flex (TFS) and BioRad CFX384 (BioRad), Hard-Shell^®^, thin-wall PCR 384-well plates (BioRad) were used. For experiments performed using QS 5 (TFS) and 7500 Fast (TFS), MicroAmp^®^ Fast 96-well reaction plates (0.1 ml) were used. For qPCR runs performed using the RotorGene Q instrument (Qiagen), the reaction volume was doubled and contained: 10 µl of 2X EpiTect^®^ MethyLight qPCR reagent (Qiagen), 2 μl of 25mM MgCl_2_ (Applied Biosystems), 0.4 μl of 20 mg/ml BSA (BioLabs), 4.1 μl of nuclease-free water, 2 μl of primer/probe mix, 0.5 μl of diluted IPC-gBlock™ (3,000 copies, STD5, **S5 Table**), and 1 μl of BC-DNA template. The reactions were performed in 0.1 ml Strip Tubes (Qiagen), which were placed in a Rotor-Disc 72 Rotor (Qiagen). All qPCR reactions were performed in triplicate, except for assessing repeatability (five replicates) and specificity (duplicates).

### Data analysis

For qBiCo-v1, firstly the PCR efficiency and R^2^ of the standard curve of each assay was calculated to verify the success of the qPCR run. The baseline threshold of each assay was determined in a qualitative manner, by setting it at the beginning of the exponential (geometric) phase. The equation of each standard curve is obtained by the BioRad CFX Maestro software using the following formula:

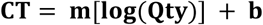

where m is the slope, b the y-intercept, and Qty the starting DNA quantity of each individual standard. According to this equation, the software provides information about the R^2^, slope and PCR efficiency. When R^2^ ≥ 0,985, -3,3 ≤ slope ≤ 3,6, and 90 % ≤ PCR efficiency ≤ 110 %, the qPCR run was indicated as successful. Regarding the qBiCo indices, we followed the following formulas:

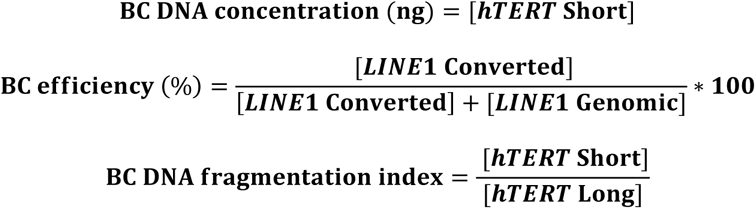

Naturally, the intact BC-DNA ratio is calculated as 1 minus the fragmented BC-DNA amount. The presence of PCR inhibition was indicated qualitatively if the average IPC-Cq of the sample was higher than the average plus standard deviation (SD) of the average IPC-Cq of the synthetic standards.

For qBiCo-v2, a semi-automated pipeline using a spreadsheet macro file (Microsoft Excel) was developed to provide with a quicker and more standardized approach. Because of the different set-up among instruments, we created three files in total: **S1 File** for 96-well, **S2 File** for 72-well and **S3 File** for 384-well set-up. Here is a short step-by-step description of the analysis: First, results including the well position, targeted assay, fluorophore, sample name, and Cq values are manually extracted from the instrument. On the “Sample Data” tab of the data analysis file, the plate layout is imported, while the extracted results are imported on the “Raw Results” tab of the file. Only in the case of 72-well setup, results are first imported on the “Rotorgene72_FormatCorrection” tab to be converted to a suitable form before importing them on the “Raw Results” tab. Unless mentioned differently, all samples and standards were tested in triplicate. To exclude any individual qPCR reaction outliers, we examine whether the Cq value of a technical replicate of a sample lies within two SDs of the mean Cq of the triplicates. Hence, when a replicate lies outside this range, it is excluded from analysis and automatically flagged in the pipeline. In case of duplicates, a data point is considered outlier, when: SD / average Cq > 0.02. Possible PCR inhibition is now indicated when the IPC-Cq value has more than one qPCR cycle difference than the IPC-Cq values of the synthetic standards. Following these steps, the equation and metrics of each standard curve are calculated and evaluated as described previously. Finally, the Cq values are then translated to copy numbers to calculate the qBiCo indices using the formulas mentioned above.

QC-ed qPCR and qBiCo parameter data were analysed using R (version 3.6.3, 29^th^ February 2020) with R-packages: beanplot, cowplot, data.table, dplyr, ggbreak, ggplot2, ggpubr, ggrepel, grid, gridExtra, gtable, plyr, reshape2, rio, rstatix and writexl. During instrument-specific qBiCo-v2 validation, we employed various methods to assess differences with statistical significance depending on the type of data. Particularly, we used the non-parametric Wilcoxon signed-rank test to assess differences between two gDNA inputs (repeatability). On the other hand, we employed the non-parametric Kruskal-Wallis test (one-way ANOVA) to assess differences between decreasing BC-DNA inputs (reproducibility/sensitivity) and among differently treated BC-DNA samples (robustness). Finally, we used student’s t-test to assess pairwise differences between the optimal and changed conditions for all experiments. We assessed significance based on difference levels: *: p < 0.05, **: p < 0.01, ***: p < 0.001, ****: p < 0.0001.

## Acknowledgments

We would like to thank previous members of the Department of Genetic Identification (Erasmus MC) for their technical assistance and contributions to this project, in particular Rochelle Chotkan during qPCR primer design and development of single-plex reactions, Diego Montiel González during *LINE1* primer design and Benjamin Planterose Jiménez during implementation of R for data analysis. Floor Claessens contributed to this study as part of her bachelor’s education (Forensic Research) at Hogeschool van Amsterdam, with work carried out during her research internship at the Department of Genetic Identification at Erasmus MC. Furthermore, we would also like to thank the multiple collaborators for enabling us access to different qPCR instruments: Isabel Chu (Department of Haematology, Erasmus MC, TFS QuantStudio 5), Michael Verbiest (Department of Internal Medicine, Erasmus MC, TFS QuantStudio 7 Flex), Claudia Erpelinck (Department of Haematology, Erasmus MC, TFS 7500 Fast), and Cathleen van der Lee (QIAGEN, RotorGene Q).

## Author contributions

**Conceptualization:** Athina Vidaki; **Methodology:** Faidra Karkala, Athina Vidaki; **Investigation:** Faidra Karkala, Floor Claessens, Vivian Kalamara; **Data curation:** Faidra Karkala, Roy Simons, Athina Vidaki; **Formal analysis:** Faidra Karkala, Roy Simons, Athina Vidaki; **Visualization:** Roy Simons, Athina Vidaki; **Funding acquisition:** Athina Vidaki; **Resources:** Manfred Kayser; **Supervision:** Athina Vidaki; **Writing – original draft:** Athina Vidaki; **Writing – review & editing:** Faidra Karkala, Roy Simons, Floor Claessens, Vivian Kalamara, Manfred Kayser.

## Abbreviations

BC: bisulfite conversion
BC-DNA: bisulfite-converted DNA
CpG: cytosine-guanine dinucleotides
Cq: quantification cycle
DNA: deoxyribonucleic acid
gDNA: genomic DNA
*hTERT*: human telomerase reverse transcriptase
IPC: internal positive control
*LINE1*: long interspersed nuclear element 1
PCR: polymerase chain reaction
RFU: relative fluorescent units
qBiCo: qualification/quantitation of bisulfite-converted DNA
QC: quality control
qPCR: quantitative PCR
SD: standard deviation
STD: standard
TFS: Thermo Fisher Scientific
*TPT1*: tumour protein, translationally-controlled 1
TRL: technology readiness level.

## Data availability

The authors confirm that all data underlying the findings of the study are fully available without restriction. All relevant data can be found within the paper and its Supporting Information files, including qBiCo index data and semi-automated analysis files.

