## Supplementary figures for "qBiCo: A method to assess global DNA conversion performance in epigenetics via single-copy genes and repetitive elements"

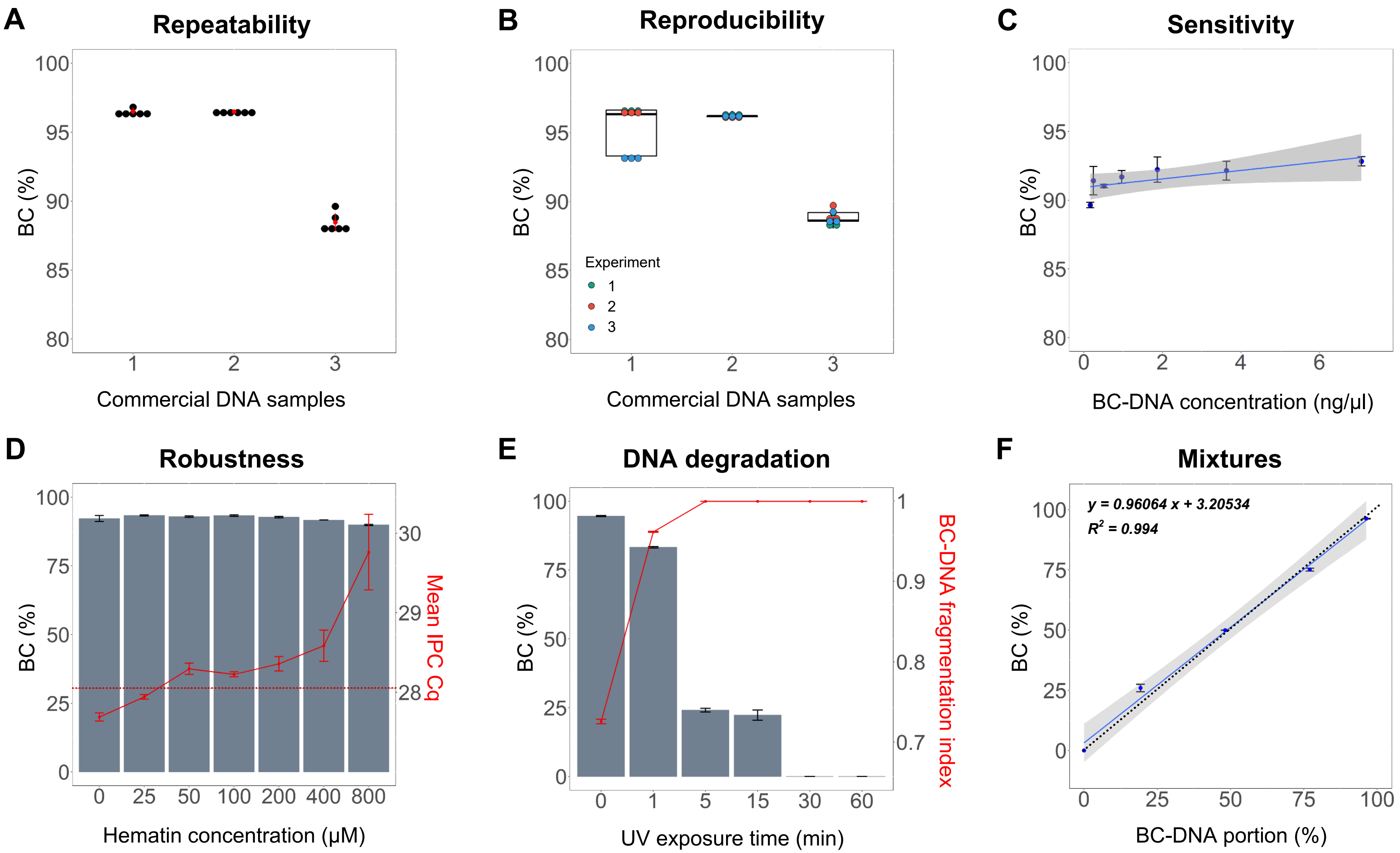


**S1 Fig. Initial qBiCo-v1 validation testing in terms of BC efficiency.** (A) Three different BC-DNA samples in six replicates within the same qPCR run, (B) Three different BC-DNA samples in three replicates within three different qPCR runs, (C) Eight BC-DNA samples with concentrations ranging from 6,25 - 0,048 ng/µl, (D) Seven BC-DNA samples exposed to different concentrations of PCR inhibitor hematin (0 - 800 µmol/L), (E) Six BC-DNA samples after initial exposure to UV for different times ranging from 0 to 60 minutes prior to conversion, (F) Five BC-DNA samples containing different ratios of converted/non-converted DNA ranging from 0 - 100 %. Dotted line corresponds to y = x. BC: Bisulfite conversion.

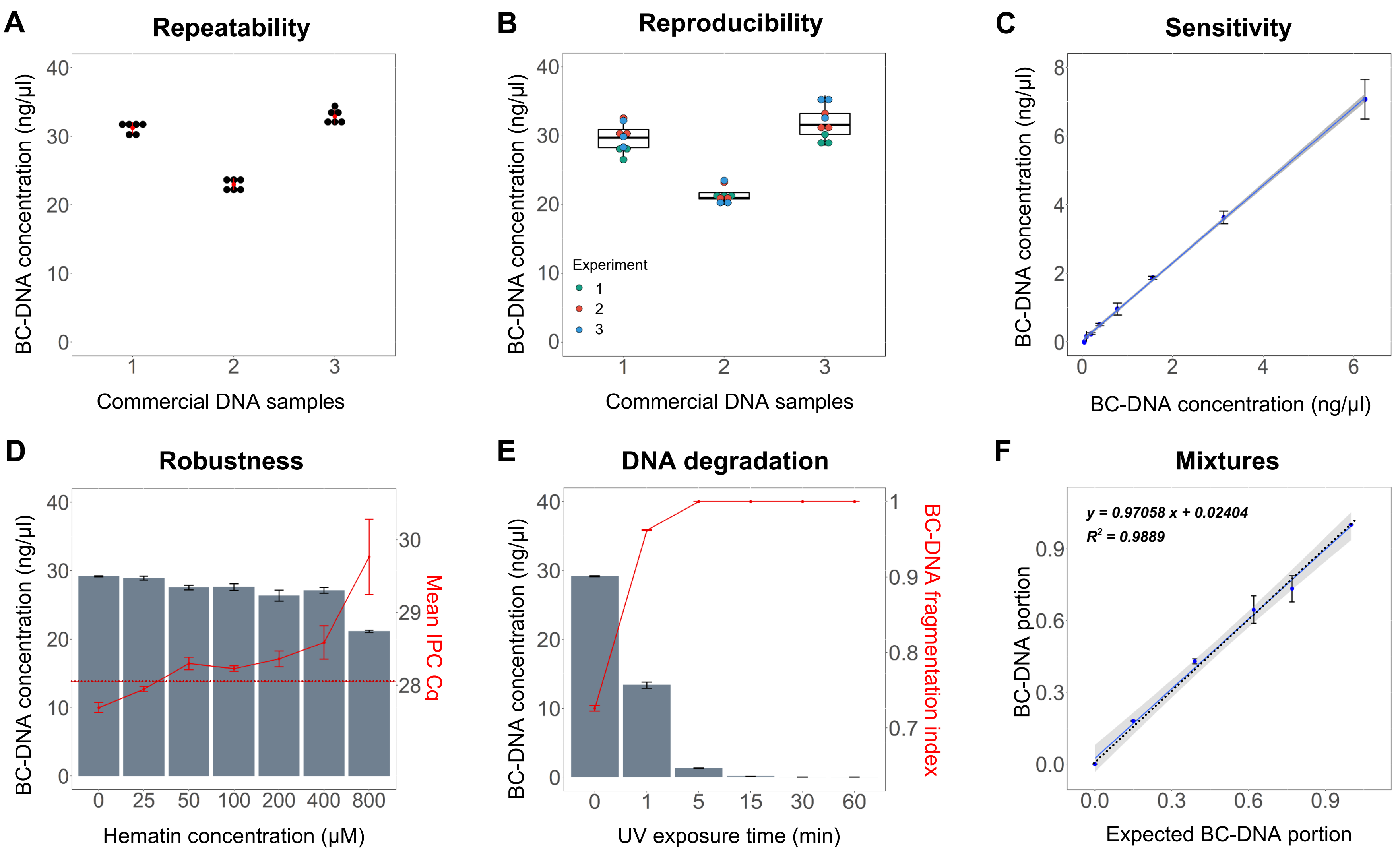


**S2 Fig. Initial qBiCo-v1 validation testing in terms of BC-DNA concentration.** (A) Three different BC-DNA samples in six replicates within the same qPCR run, (B) Three different BC-DNA samples in three replicates within three different qPCR runs, (C) Eight BC-DNA samples with concentrations ranging from 6,25 - 0,048 ng/µl, (D) Seven BC-DNA samples exposed to different concentrations of PCR inhibitor hematin (0 - 800 µmol/L), (E) Six BC-DNA samples after initial exposure to UV for different times ranging from 0 to 60 minutes prior to conversion, (F) Six BC-DNA samples containing a different portion of BC-DNA measured using 2 ng. Dotted line corresponds to y = x. BC: Bisulfite conversion.

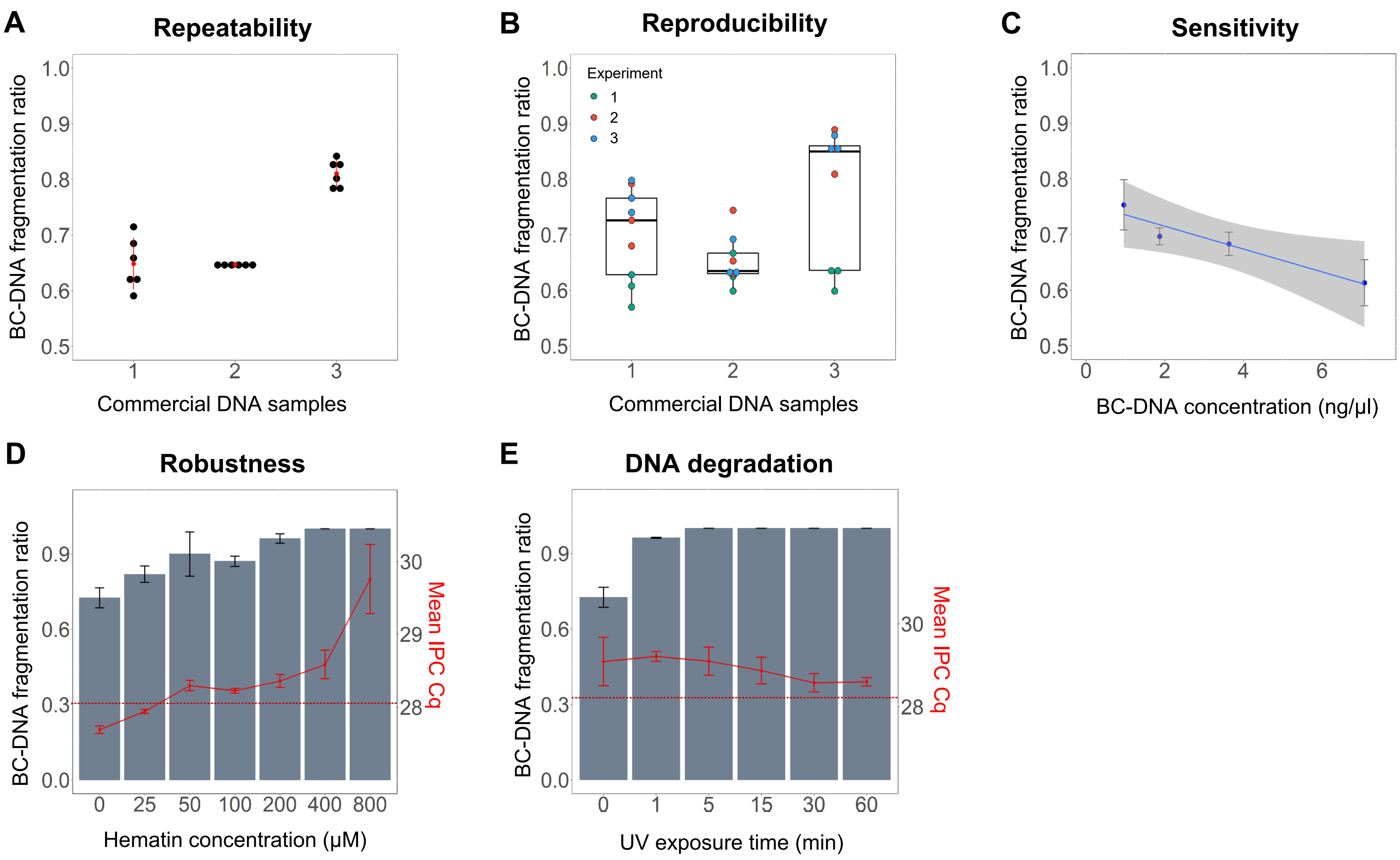


**S3 Fig. Initial qBiCo-v1 validation testing in terms of BC-DNA fragmentation.** (A) Three different BC-DNA samples in six replicates within the same qPCR run, (B) Three different BC-DNA samples in three replicates within three different qPCR runs, (C) Eight BC-DNA samples with concentrations ranging from 6,25 - 0,048 ng/µl (with missing data), (D) Seven BC-DNA samples exposed to different concentrations of PCR inhibitor hematin (0 - 800 µmol/L), (E) Six BC-DNA samples after initial exposure to UV for different times ranging from 0 to 60 minutes prior to conversion. BC: Bisulfite conversion.

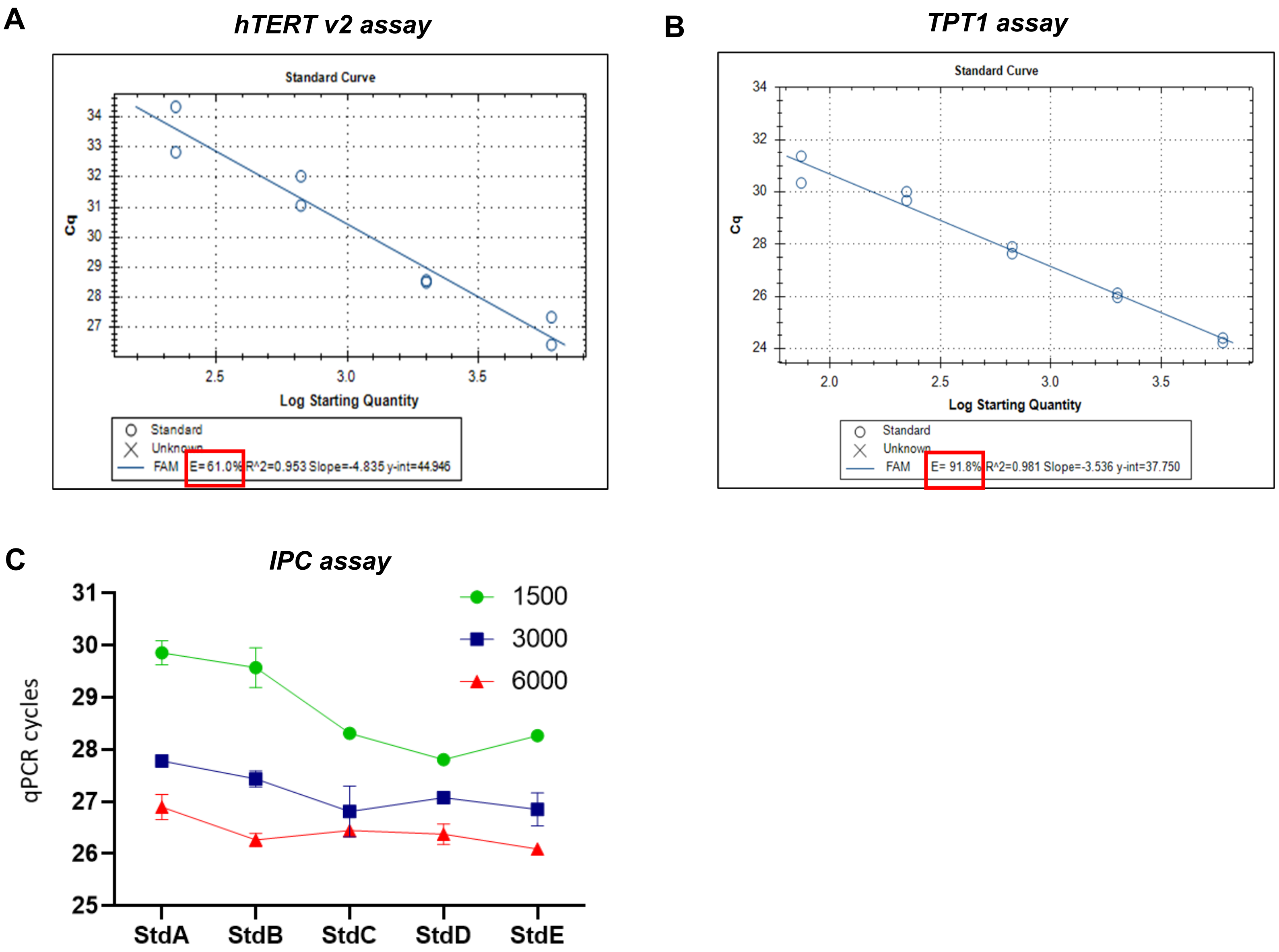


**S4 Fig. Optimisation and fine-tuning of individual qBiCo-v2 assays.** Replacement of the original long assay using (A) a redesigned (*hTERT*) and (B) a new, improved version (*TPT1*) based on a five-standard curve, (C) Determination of the ideal artificial fragment copy number (IPC) to be pre-added in the qPCR reaction based on the detection in the synthetic standards. Error bars correspond to standard deviation.

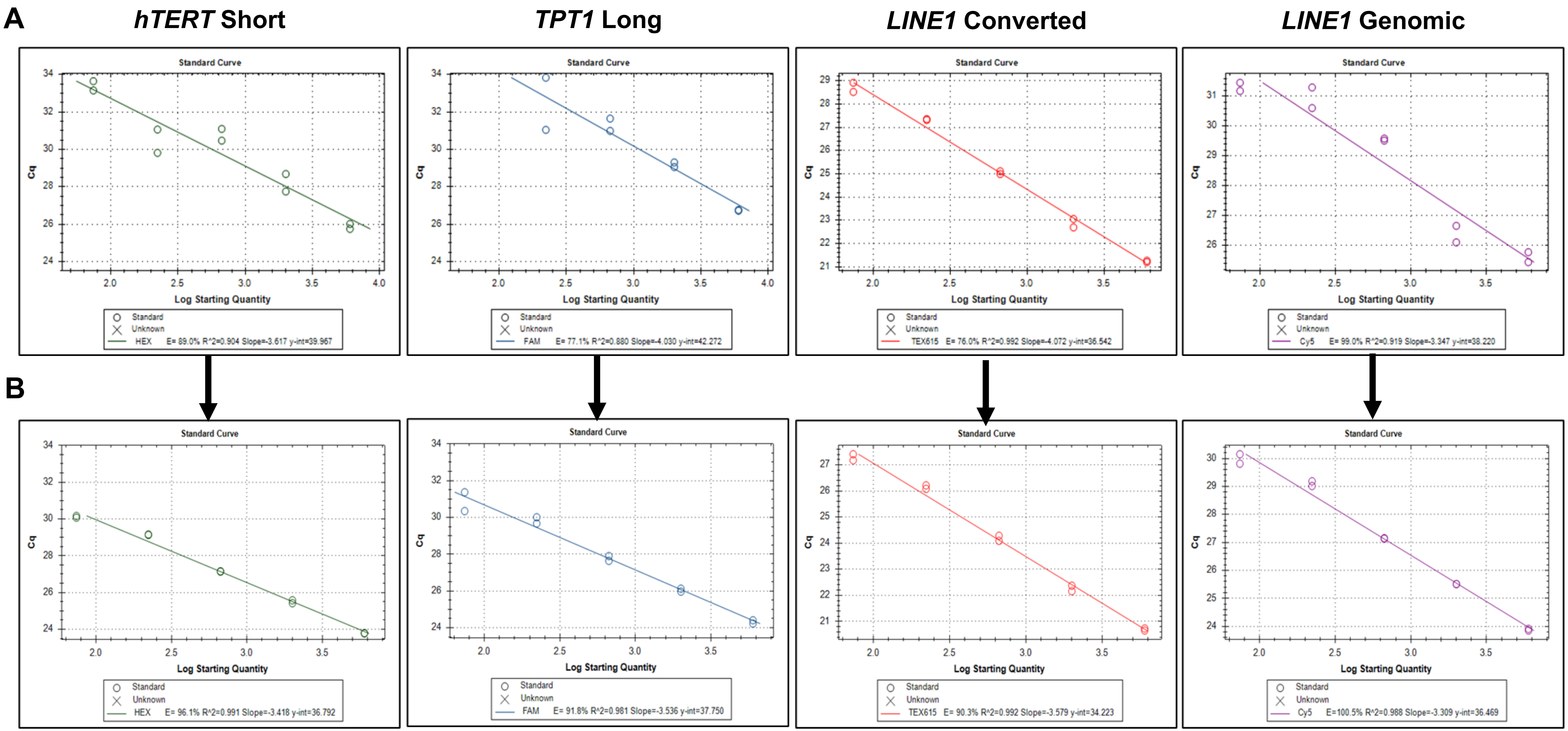


**S5 Fig. Improvement of standard curve (n = 5 STDs) performance for all qBiCo-v2 assays by adjusting the dilution approach of the synthetic standards.** (A) Old dilution approach in $H_2O$, (B) New dilution approach in TE. E: PCR efficiency. STD: Synthetic standard

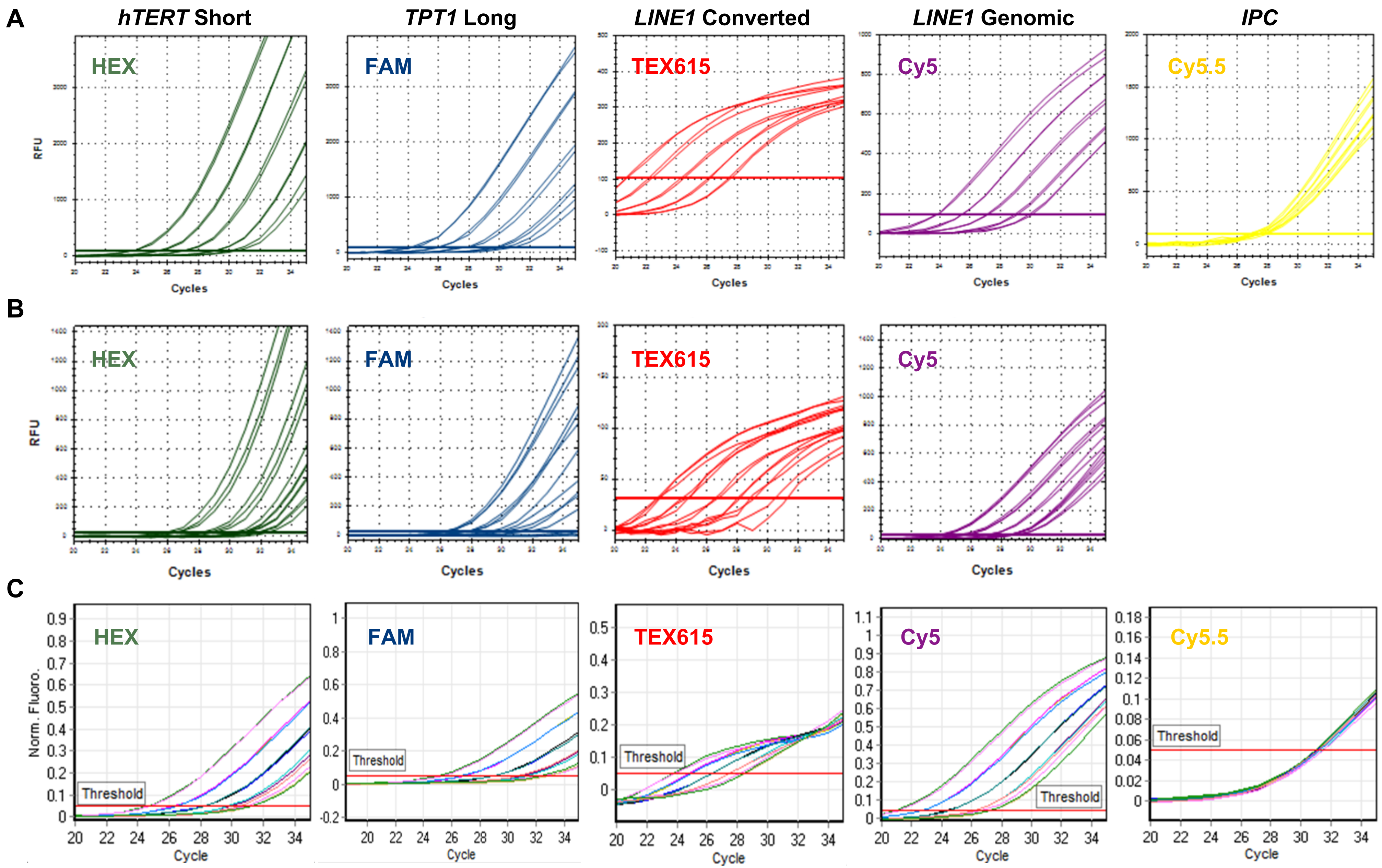


**S6 Fig. Amplification curves for each individual assay of qBiCo-v2 for the BioRad and Qiagen systems.** (A) BioRad CFX96, (B) BioRad CFX384 (4plex without IPC) and (C) Qiagen RotorGene Q. Labels correspond to fluorophores used.

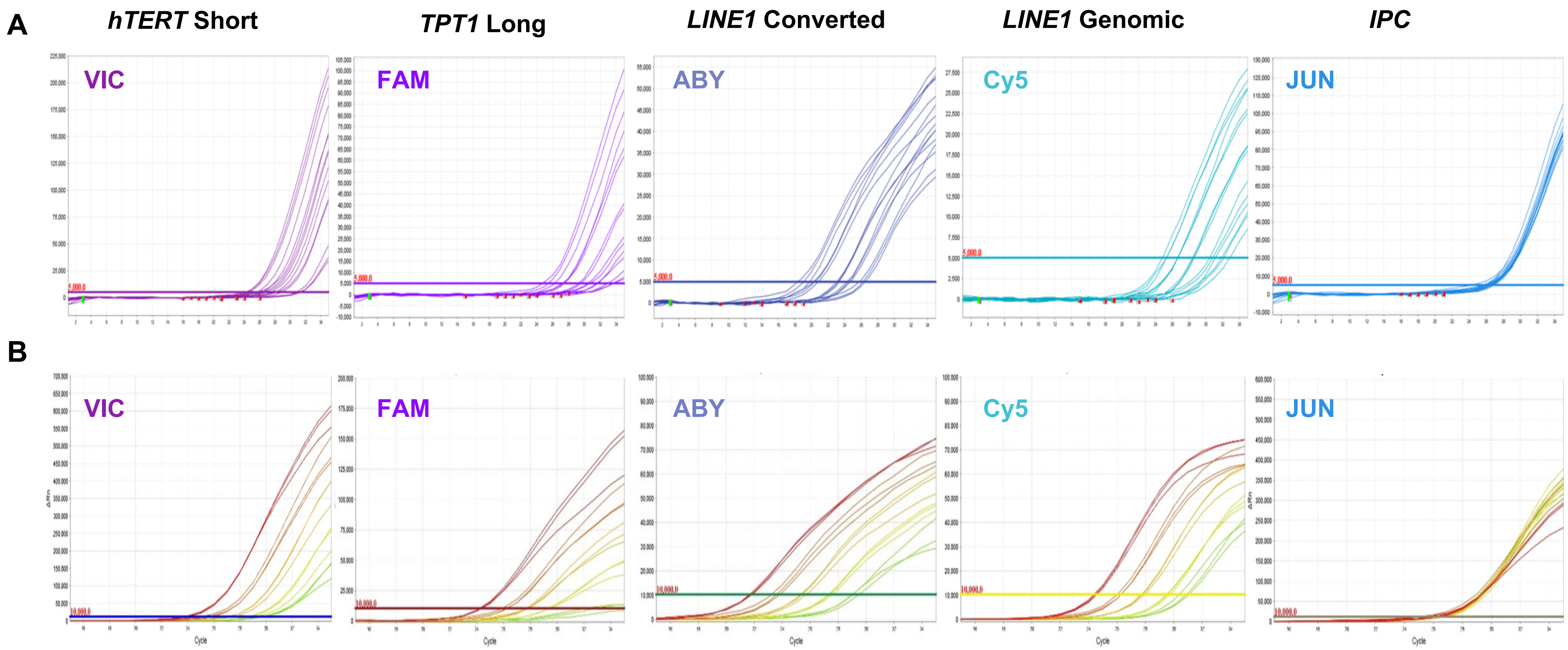


**S7 Fig. Amplification curves for each individual assay of qBiCo-v2 for the Thermo Fisher Scientific systems.** (A) QuantStudio 5 and (B) QuantStudio 7 Flex. Labels correspond to fluorophores used.

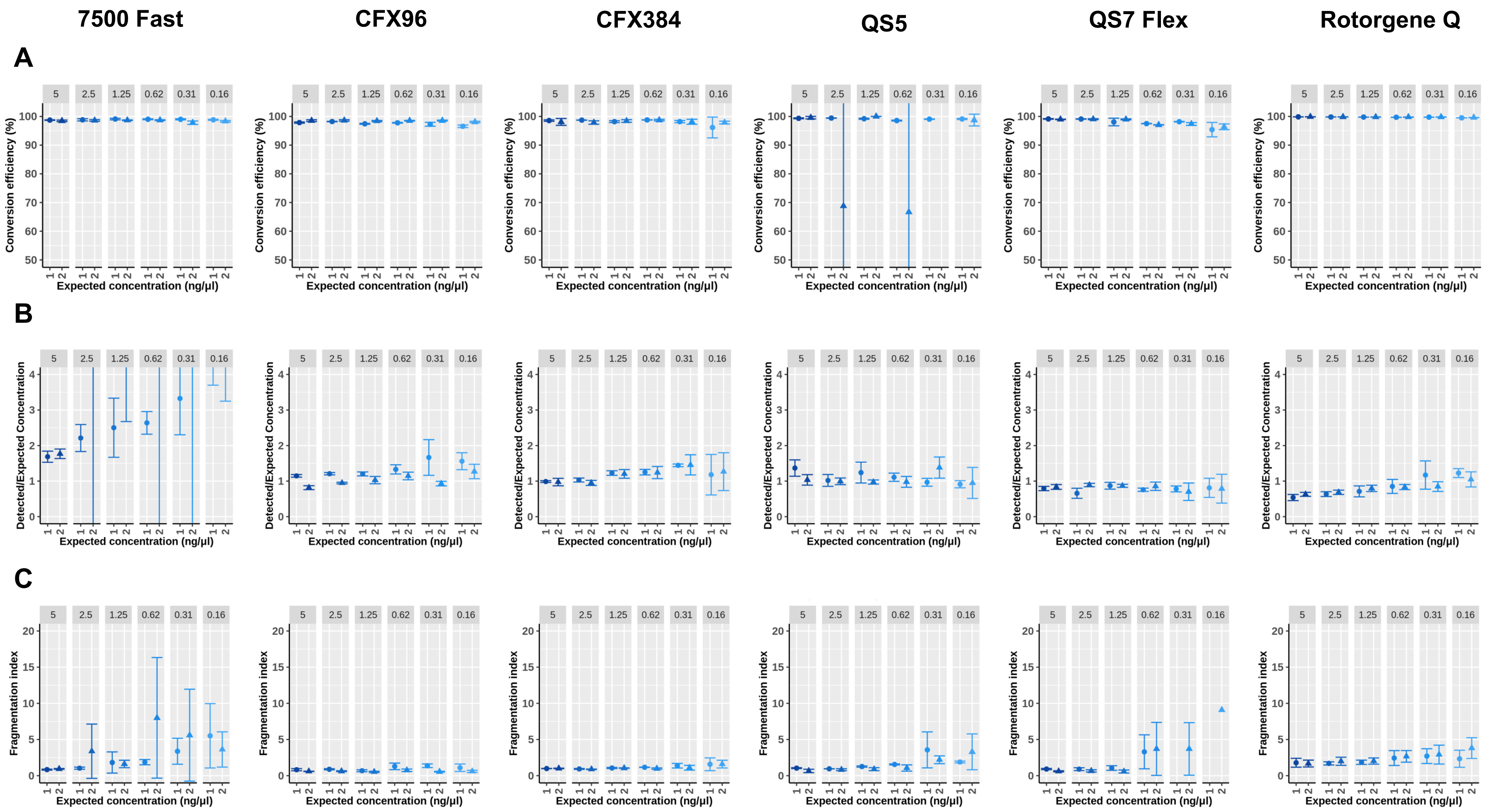


**S8 Fig. Reproducibility testing at six different BC-DNA amounts via two separate qPCR experiments during qBiCo-v2 assay validation on various qPCR systems.** (A) BC efficiency (%), (B) Detected/Expected BC-DNA concentration ratio and (C) BC-DNA fragmentation index. Labels correspond to fluorophores used. BC: Bisulfite conversion.
