## Supplementary tables for "qBiCo: A method to assess global DNA conversion performance in epigenetics via single-copy genes and repetitive elements"

**S1 Table. Information on qBiCo-v1 method.** Sequences of target DNA regions, primer, probes and synthetic DNA standards per qPCR assay. Expected converted Cs on the *LINE1* Converted probe sequence are indicated in green.

| **Assay** | **Oligo** | **Sequence (5'→3')** | **Concentration (µM)** | **Fluorophore** | **Amplicon length (bp)** |
|---|---|---|---|---|---|
| ***hTERT* Short** | Forward primer | TGGGTTTTAGAGTTGATTTTT | 0.6 | NA | 85 |
| | Reverse primer | ACCACATTAAACAATCCCCT | 0.6 | | |
| | Probe | AATTAGGTATAGGGGATTTGGTTTTAGT | 0.3 | HEX™ | |
| | Synthetic standard | TGGGTTTTAGAGTTGATTTTTGTGAATTAAATTAAAATTAGGTATAGGGGATTTGGTTTTAGTATAGGGGATTGTTTAATGTGG<br>T | NA | NA | |
| ***hTERT* Long** | Forward primer | AGGGTTTTAGTTAGAAGATT | 0.2 | NA | 235 |
| | Reverse primer | CATTATATATTAACTTATTCCCAC | 0.2 | | |
| | Probe | GTGTTGGGGTTATTTTTTTGTATTTGG | 0.1 | FAM™ | |
| | Synthetic standard | AGGGTTTTAGTTAGAAGATTAGGGTTTTTTTAGTTTTTTTGTATATTCGAGTTTTTGGGGGGTTTTGTGATATTTTATGTTTTAA<br>ATTAGGATGTTTGTAGAGGGAGTTGGTAGTAGATTTCGTTAGAGGTAATATAGTTTTTGGGTTGGGGATTTCGACGTGGTGT<br>TGGGGTTATTTTTTTGTATTTGGGGGAGGGTTAGGGTTTTTTTGTGGGAATAAGTTAATATATAATG | NA | NA | |
| ***LINE1* Converted** | Forward primer | TTTTGAGTTAGGTGTGGGATATA | 0.1 | NA | 148 |
| | Reverse primer | AAAATCAAAAAATTCCCTTTC | 0.1 | | |
| | Probe | TGGGAGTGATTTGATTTTTTAGGTGY | 0.1 | TEX615™ | |
| | Synthetic standard | TTTTGAGTTAGGTGTGGGATATAATTTTTTGGTGTGTTGTTTTTTAAGTTTTTTGGAAAAGTGTAGTATTTAGGGTGGGAGTG<br>ATTTGATTTTTTAGGTGTTGTTTGTTATTTTTTTTTTTGATTAGGAAAGGGAATTTTTTGATTTT | NA | NA | |
| ***LINE1* Genomic** | Forward primer | GAGCCAGGTGTGGGATATAA | 0.1 | NA | 119 |
| | Reverse primer | TCAAAGAAAGGGGTGACAGA | 0.1 | | |
| | Probe | TGGGAGTGACCCAATTTTCCAGGTG | 0.1 | Cy5™ | |
| | Synthetic standard | GAGCCAGGTGTGGGATATAATCTCCTGGTGTGCTGTTTTTTAAGCCCTTTGGAAAAGTGCAGTATTTAGGGTGGGAGTGAC<br>CCGATTTTCCAGGTGCCGTCTGTCACCCCTTTCTTTGA | NA | NA | |
| **Synthetic IPC** | Forward primer | GGTGATTTTTAATAATTTTTGAGG | 0.2 | NA | 99 |
| | Reverse primer | ACTTTCAACCCATTACCTCAT | 0.2 | | |
| | Probe | AGTGTTCGTATAGGTTTTTTTTTTTAGT | 0.1 | Cy5.5™ | |
| | Synthetic standard | GGTGATTTTTAATAATTTTTGAGGGAGTATTGGTATAGGCGATAGAGTGTTCGTATAGGTTTTTTTTTTTAGTGGATGATGAG<br>GTAATGGGTTGAAAGT | NA | NA | |

**S2 Table. Data obtained by evaluating qBiCo-v1 assay using standard performance parameters.** Information is provided per sample, condition and unit, including additional data for a subset. These data were used to produce S1 - S3 figures. BC: Bisulfite conversion.

| Performance parameter | BC performance parameter | Additional information | | Replicates | | | | | | | | |
|---|---|---|---|---|---|---|---|---|---|---|---|---|
| | | | | Samples | | | IPC Cq | | | Fragmentation index | | |
| | | | | 1 | 2 | 3 | 1 | 2 | 3 | 1 | 2 | 3 |
| Repeatability | BC efficiency (%) | Run1 | | 96.60 | 96.18 | 88.27 | NA | NA | NA | NA | NA | NA |
| | | | | 96.45 | 96.57 | 88.21 | NA | NA | NA | NA | NA | NA |
| | | | | 96.04 | 96.58 | 87.73 | NA | NA | NA | NA | NA | NA |
| | | | | 96.82 | 96.55 | 88.15 | NA | NA | NA | NA | NA | NA |
| | | | | 96.62 | 96.35 | 88.79 | NA | NA | NA | NA | NA | NA |
| | | | | 96.56 | 96.65 | 89.62 | NA | NA | NA | NA | NA | NA |
| | BC-DNA concentration (ng/µl) | | | 29.74 | 24.12 | 34.09 | NA | NA | NA | NA | NA | NA |
| | | | | 31.92 | 23.48 | 34.40 | NA | NA | NA | NA | NA | NA |
| | | | | 31.44 | 21.58 | 31.81 | NA | NA | NA | NA | NA | NA |
| | | | | 31.43 | 22.64 | 32.73 | NA | NA | NA | NA | NA | NA |
| | | | | 30.74 | 23.17 | 31.44 | NA | NA | NA | NA | NA | NA |
| | | | | 32.02 | 22.86 | 32.80 | NA | NA | NA | NA | NA | NA |
| | BC-DNA fragmentation ratio | | | 0.62 | 0.65 | 0.83 | NA | NA | NA | NA | NA | NA |
| | | | | 0.62 | 0.64 | 0.80 | NA | NA | NA | NA | NA | NA |
| | | | | 0.66 | 0.65 | 0.82 | NA | NA | NA | NA | NA | NA |
| | | | | 0.59 | 0.64 | 0.79 | NA | NA | NA | NA | NA | NA |
| | | | | 0.69 | 0.65 | 0.84 | NA | NA | NA | NA | NA | NA |
| | | | | 0.72 | 0.65 | 0.78 | NA | NA | NA | NA | NA | NA |
| Reproducibility | BC efficiency (%) | Run1 | | 96.33 | 96.07 | 88.02 | NA | NA | NA | NA | NA | NA |
| | | | | 96.79 | 96.49 | 89.20 | NA | NA | NA | NA | NA | NA |
| | | | | 96.52 | 96.20 | 88.57 | NA | NA | NA | NA | NA | NA |
| | | Run2 | | 96.62 | 96.01 | 88.91 | NA | NA | NA | NA | NA | NA |
| | | | | 96.61 | 96.20 | 89.70 | NA | NA | NA | NA | NA | NA |
| | | | | 96.26 | 96.14 | 88.62 | NA | NA | NA | NA | NA | NA |
| | | Run3 | | 92.98 | 96.16 | 88.62 | NA | NA | NA | NA | NA | NA |
| | | | | 93.29 | 96.19 | 89.25 | NA | NA | NA | NA | NA | NA |
| | | | | 93.07 | 96.20 | 88.50 | NA | NA | NA | NA | NA | NA |
| | BC-DNA concentration (ng/µl) | Run1 | | 27.87 | 21.25 | 30.19 | NA | NA | NA | NA | NA | NA |
| | | | | 28.26 | 20.83 | 29.37 | NA | NA | NA | NA | NA | NA |
| | | | | 26.53 | 21.71 | 28.54 | NA | NA | NA | NA | NA | NA |
| | | Run2 | | 29.71 | 23.24 | 31.59 | NA | NA | NA | NA | NA | NA |
| | | | | 32.55 | 20.92 | 33.22 | NA | NA | NA | NA | NA | NA |
| | | | | 30.91 | 20.98 | 30.77 | NA | NA | NA | NA | NA | NA |
| | | Run3 | | 28.32 | 20.60 | 32.57 | NA | NA | NA | NA | NA | NA |
| | | | | 32.19 | 19.97 | 34.64 | NA | NA | NA | NA | NA | NA |
| | | | | 29.89 | 23.49 | 35.84 | NA | NA | NA | NA | NA | NA |
| | BC-DNA fragmentation ratio | Run1 | | 0.63 | 0.63 | 0.64 | NA | NA | NA | NA | NA | NA |
| | | | | 0.57 | 0.67 | 0.64 | NA | NA | NA | NA | NA | NA |
| | | | | 0.61 | 0.60 | 0.60 | NA | NA | NA | NA | NA | NA |
| | | Run2 | | 0.68 | 0.74 | 0.89 | NA | NA | NA | NA | NA | NA |
| | | | | 0.73 | 0.63 | 0.81 | NA | NA | NA | NA | NA | NA |
| | | | | 0.79 | 0.65 | 0.86 | NA | NA | NA | NA | NA | NA |
| | | Run3 | | 0.74 | 0.64 | 0.86 | NA | NA | NA | NA | NA | NA |
| | | | | 0.80 | 0.63 | 0.85 | NA | NA | NA | NA | NA | NA |
| | | | | 0.77 | 0.69 | 0.88 | NA | NA | NA | NA | NA | NA |
| Sensitivity | BC efficiency (%) | Measured BC-DNA concentration | 7.07 | 93.09 | 92.96 | 92.45 | NA | NA | NA | NA | NA | NA |
| | | | 3.63 | 92.93 | 91.89 | 91.64 | NA | NA | NA | NA | NA | NA |
| | | | 1.87 | 93.29 | 91.64 | 91.75 | NA | NA | NA | NA | NA | NA |
| | | | 0.96 | 91.18 | 92.06 | 91.85 | NA | NA | NA | NA | NA | NA |
| | | | 0.51 | 90.96 | 91.18 | 90.96 | NA | NA | NA | NA | NA | NA |
| | | | 0.24 | 92.02 | 90.23 | 92.02 | NA | NA | NA | NA | NA | NA |
| | | | 0.16 | 89.76 | 89.42 | 89.76 | NA | NA | NA | NA | NA | NA |
| | | | 0 | 0 | 0 | 0 | NA | NA | NA | NA | NA | NA |
| | BC-DNA concentration (ng/µl) | Expected BC-DNA concentration | 6.25 | 6.45 | 7.17 | 7.59 | NA | NA | NA | NA | NA | NA |
| | | | 3.13 | 3.44 | 3.62 | 3.81 | NA | NA | NA | NA | NA | NA |
| | | | 1.56 | 1.91 | 1.82 | 1.89 | NA | NA | NA | NA | NA | NA |
| | | | 0.78 | 0.98 | 0.78 | 1.12 | NA | NA | NA | NA | NA | NA |
| | | | 0.39 | 0.53 | 0.53 | 0.46 | NA | NA | NA | NA | NA | NA |
| | | | 0.195 | 0.22 | 0.27 | 0.22 | NA | NA | NA | NA | NA | NA |
| | | | 0.097 | 0.31 | 0.00 | 0.18 | NA | NA | NA | NA | NA | NA |
| | | | 0.048 | 0 | 0 | 0 | NA | NA | NA | NA | NA | NA |
| | BC-DNA fragmentation ratio | Measured BC-DNA concentration | 7.07 | 0.66 | 0.58 | 0.60 | NA | NA | NA | NA | NA | NA |
| | | | 3.63 | 0.69 | 0.66 | 0.70 | NA | NA | NA | NA | NA | NA |
| | | | 1.87 | 0.71 | 0.68 | 0.70 | NA | NA | NA | NA | NA | NA |
| | | | 0.96 | 0.80 | 0.71 | 0.75 | NA | NA | NA | NA | NA | NA |
| | | | 0.51 | 0 | 0 | 0 | NA | NA | NA | NA | NA | NA |
| | | | 0.24 | 0 | 0 | 0 | NA | NA | NA | NA | NA | NA |
| | | | 0.16 | 0 | 0 | 0 | NA | NA | NA | NA | NA | NA |

| | | | | | | | | | | | | |
|---|---|---|---|---|---|---|---|---|---|---|---|---|
| | | | 0 | 0 | 0 | 0 | NA | NA | NA | NA | NA | NA |
| **Robustness** | BC efficiency (%) | Hematin concentration (μM) | 0 | 92.41 | 93.20 | 91.07 | 27.97 | 27.66 | 27.44 | NA | NA | NA |
| | | | 25 | 93.19 | 93.25 | 93.57 | 27.82 | 27.98 | 28.03 | NA | NA | NA |
| | | | 50 | 92.84 | 93.19 | 92.71 | 28.20 | 28.17 | 28.52 | NA | NA | NA |
| | | | 100 | 93.01 | 93.57 | 93.22 | 28.24 | 28.31 | 28.13 | NA | NA | NA |
| | | | 200 | 92.48 | 92.96 | 92.74 | 28.51 | 28.48 | 28.09 | NA | NA | NA |
| | | | 400 | 91.65 | 91.66 | 91.72 | 28.90 | 28.09 | 28.77 | NA | NA | NA |
| | | | 800 | 90.01 | 89.99 | 89.61 | 29.39 | 29.47 | 30.43 | NA | NA | NA |
| | BC-DNA concentration (ng/μl) | | 0 | 29.12 | 29.24 | 29.11 | 27.97 | 27.66 | 27.44 | NA | NA | NA |
| | | | 25 | 28.57 | 29.14 | 29.00 | 27.82 | 27.98 | 28.03 | NA | NA | NA |
| | | | 50 | 27.80 | 27.16 | 27.60 | 28.20 | 28.17 | 28.52 | NA | NA | NA |
| | | | 100 | 28.12 | 27.41 | 27.21 | 28.24 | 28.31 | 28.13 | NA | NA | NA |
| | | | 200 | 26.00 | 27.23 | 25.77 | 28.51 | 28.48 | 28.09 | NA | NA | NA |
| | | | 400 | 27.17 | 26.64 | 27.48 | 28.90 | 28.09 | 28.77 | NA | NA | NA |
| | | | 800 | 21.29 | 20.97 | 21.14 | 29.39 | 29.47 | 30.43 | NA | NA | NA |
| | BC-DNA fragmentation ratio | | 0 | 0.77 | 0.71 | 0.70 | 27.97 | 27.66 | 27.44 | NA | NA | NA |
| | | | 25 | 0.79 | 0.81 | 0.85 | 27.82 | 27.98 | 28.03 | NA | NA | NA |
| | | | 50 | 0.84 | 0.86 | 1.00 | 28.20 | 28.17 | 28.52 | NA | NA | NA |
| | | | 100 | 0.88 | 0.88 | 0.85 | 28.24 | 28.31 | 28.13 | NA | NA | NA |
| | | | 200 | 0.98 | 0.94 | 0.96 | 28.51 | 28.48 | 28.09 | NA | NA | NA |
| | | | 400 | 1 | 1 | 1 | 28.90 | 28.09 | 28.77 | NA | NA | NA |
| | | | 800 | 1 | 1 | 1 | 29.39 | 29.47 | 30.43 | NA | NA | NA |
| **DNA degradation** | BC efficiency (%) | UV exposure (sec) | 0 | 94.49 | 94.51 | 94.83 | NA | NA | NA | 0.77 | 0.71 | 0.70 |
| | | | 1 | 83.41 | 83.50 | 83.02 | NA | NA | NA | 0.96 | 0.96 | 0.96 |
| | | | 5 | 24.85 | 23.53 | 24.01 | NA | NA | NA | 1 | 1 | 1 |
| | | | 15 | 20.86 | 21.66 | 24.44 | NA | NA | NA | 1 | 1 | 1 |
| | | | 30 | 0 | 0 | 0 | NA | NA | NA | 1 | 1 | 1 |
| | | | 60 | 0 | 0 | 0 | NA | NA | NA | 1 | 1 | 1 |
| | BC-DNA concentration (ng/μl) | | 0 | 29.12 | 29.24 | 29.11 | NA | NA | NA | 0.77 | 0.71 | 0.70 |
| | | | 1 | 12.98 | 13.87 | 13.20 | NA | NA | NA | 0.96 | 0.96 | 0.96 |
| | | | 5 | 1.33 | 1.38 | 1.28 | NA | NA | NA | 1 | 1 | 1 |
| | | | 15 | 0.15 | 0.10 | 0.10 | NA | NA | NA | 1 | 1 | 1 |
| | | | 30 | 0 | 0 | 0 | NA | NA | NA | 1 | 1 | 1 |
| | | | 60 | 0 | 0 | 0 | NA | NA | NA | 1 | 1 | 1 |
| | BC-DNA fragmentation ratio | | 0 | 0.77 | 0.71 | 0.70 | 28.92 | 29.08 | 29.31 | NA | NA | NA |
| | | | 1 | 0.96 | 0.96 | 0.96 | 29.19 | 29.22 | 29.27 | NA | NA | NA |
| | | | 5 | 1 | 1 | 1 | 29.03 | 29.06 | 29.24 | NA | NA | NA |
| | | | 15 | 1 | 1 | 1 | 28.76 | 28.93 | 28.96 | NA | NA | NA |
| | | | 30 | 1 | 1 | 1 | 28.51 | 28.59 | 28.66 | NA | NA | NA |
| | | | 60 | 1 | 1 | 1 | 28.57 | 28.63 | 28.63 | NA | NA | NA |
| **Mixtures** | BC efficiency (%) | Measured using 2ng as BC-DNA input | 96.30 | 96.30 | 96.42 | 96.17 | NA | NA | NA | NA | NA | NA |
| | | | 77.04 | 74.47 | 75.54 | 75.40 | NA | NA | NA | NA | NA | NA |
| | | | 48.15 | 49.95 | 49.86 | 49.96 | NA | NA | NA | NA | NA | NA |
| | | | 19.26 | 27.49 | 25.97 | 24.37 | NA | NA | NA | NA | NA | NA |
| | | | 0 | 0 | 0 | 0 | NA | NA | NA | NA | NA | NA |
| | BC-DNA portion | | 1 | 1 | 1 | 1 | NA | NA | NA | NA | NA | NA |
| | | | 0.77 | 0.90 | 0.71 | 0.64 | NA | NA | NA | NA | NA | NA |
| | | | 0.62 | 0.60 | 0.71 | 0.62 | NA | NA | NA | NA | NA | NA |
| | | | 0.39 | 0.42 | 0.43 | 0.44 | NA | NA | NA | NA | NA | NA |
| | | | 0.15 | 0.18 | 0.18 | 0.18 | NA | NA | NA | NA | NA | NA |
| | | | 0 | 0 | 0 | 0 | NA | NA | NA | NA | NA | NA |

**S3 Table. Data obtained by testing ten commercially available BC kits based the qBiCo-v1 assay using different initial gDNA amounts.** Information on kits used are found in the Methods. Data are presented in ratios, with 1 representing the ideal condition per index. These data were used to produce figure 3. BC: Bisulfite conversion; gDNA: Genomic DNA

| Manufacturer | Kit No | Methylated standard / Manufacturer | BC performance parameter | qBiCo replicates | | | | | | | | | | | | | | |
|---|---|---|---|---|---|---|---|---|---|---|---|---|---|---|---|---|---|---|
| | | | | 200ng | | | 100ng | | | 50ng | | | 10ng | | | 1ng | | |
| | | | | 1 | 2 | 3 | 1 | 2 | 3 | 1 | 2 | 3 | 1 | 2 | 3 | 1 | 2 | 3 |
| **Abcam** | 7 | EpigenDX | BC-DNA recovery | 0.513 | 0.545 | 0.595 | 0.544 | 0.534 | 0.514 | 0.474 | 0.538 | 0.465 | 0.700 | 0.650 | 0.565 | NA | NA | NA |
| | | TFS | | 0.465 | 0.400 | 0.375 | 0.293 | 0.317 | 0.291 | 0.535 | 0.526 | 0.538 | 0.375 | 0.397 | 0.344 | NA | NA | NA |
| | | ZymoResearch | | 0.482 | 0.371 | 0.386 | 0.314 | 0.309 | 0.328 | 0.215 | 0.246 | 0.235 | 0.429 | 0.373 | 0.344 | NA | NA | NA |
| | | EpigenDX | Intact BC-DNA | 0.101 | 0.088 | 0.075 | 0.081 | 0.112 | 0.119 | NA | NA | NA | NA | NA | NA | NA | NA | NA |
| | | TFS | | 0.094 | 0.093 | 0.105 | 0.081 | 0.091 | 0.098 | NA | NA | 0.101 | NA | NA | NA | NA | NA | NA |
| | | ZymoResearch | | 0.087 | 0.080 | 0.096 | 0.074 | 0.093 | 0.059 | NA | NA | NA | NA | NA | NA | NA | NA | NA |
| | | EpigenDX | BC efficiency | 1.000 | 1.000 | 1.000 | 1.000 | 1.000 | 1.000 | 0.997 | 0.998 | 0.997 | 0.983 | 0.986 | 0.985 | 0.750 | 0.816 | 0.792 |
| | | TFS | | 0.998 | 0.999 | 0.999 | 0.999 | 0.999 | 0.999 | 1.000 | 1.000 | 1.000 | 0.988 | 0.988 | 0.989 | 0.855 | 0.838 | 0.847 |
| | | ZymoResearch | | 1.000 | 1.000 | 1.000 | 1.000 | 0.999 | 0.999 | 0.998 | 1.000 | 1.000 | 1.000 | 1.000 | 1.000 | 0.923 | 0.837 | 0.938 |
| **Active Motif** | 10 | EpigenDX | BC-DNA recovery | 0.280 | 0.279 | 0.292 | 0.167 | 0.206 | 0.175 | 0.185 | 0.171 | 0.181 | NA | NA | 0.154 | NA | NA | NA |
| | | TFS | | 0.037 | 0.047 | 0.036 | 0.043 | 0.049 | 0.053 | 0.049 | 0.045 | 0.038 | NA | NA | NA | NA | NA | NA |
| | | ZymoResearch | | 0.178 | 0.174 | 0.160 | 0.150 | 0.167 | 0.171 | 0.075 | 0.082 | 0.081 | 0.222 | 0.179 | 0.239 | NA | NA | NA |
| | | EpigenDX | Intact BC-DNA | NA | NA | NA | NA | NA | NA | NA | NA | NA | NA | NA | NA | NA | NA | NA |
| | | TFS | | NA | NA | NA | NA | NA | NA | NA | NA | NA | NA | NA | NA | NA | NA | NA |
| | | ZymoResearch | | NA | NA | NA | NA | NA | NA | NA | NA | NA | NA | NA | NA | NA | NA | NA |
| | | EpigenDX | BC efficiency | 0.175 | 0.202 | 0.202 | 0.786 | 0.796 | 0.789 | 0.245 | 0.272 | 0.266 | 0.499 | 0.541 | 0.545 | 0.282 | 0.314 | 0.314 |
| | | TFS | | 0.059 | 0.065 | 0.061 | 0.045 | 0.041 | 0.040 | 0.028 | 0.028 | 0.023 | 0.025 | 0.021 | 0.420 | 0.279 | 0.009 | 0.314 |
| | | ZymoResearch | | 0.267 | 0.273 | 0.299 | 0.268 | 0.279 | 0.276 | 0.240 | 0.254 | 0.255 | 0.357 | 0.381 | 0.380 | 0.369 | 0.245 | 0.367 |
| **Analytik Jena** | 8 | EpigenDX | BC-DNA recovery | 0.505 | 0.562 | 0.525 | 0.243 | 0.264 | 0.253 | 0.329 | 0.306 | 0.344 | 0.394 | 0.376 | 0.441 | NA | NA | NA |
| | | TFS | | 0.379 | 0.329 | 0.363 | 0.407 | 0.461 | 0.437 | 0.506 | 0.494 | 0.489 | 0.375 | 0.406 | 0.373 | NA | NA | NA |
| | | ZymoResearch | | 0.249 | 0.231 | 0.228 | 0.405 | 0.415 | 0.429 | 0.369 | 0.381 | 0.285 | 0.332 | 0.312 | 0.456 | NA | NA | NA |
| | | EpigenDX | Intact BC-DNA | 0.366 | 0.312 | 0.394 | 0.363 | 0.428 | 0.359 | 0.380 | 0.390 | 0.385 | NA | NA | NA | NA | NA | NA |
| | | TFS | | 0.432 | 0.425 | 0.417 | 0.432 | 0.382 | 0.440 | 0.497 | 0.460 | 0.417 | NA | 0.645 | NA | NA | NA | NA |
| | | ZymoResearch | | 0.356 | 0.420 | 0.393 | 0.450 | 0.380 | 0.381 | 0.420 | 0.317 | 0.474 | 0.408 | NA | 0.321 | NA | NA | NA |
| | | EpigenDX | BC efficiency | 1.000 | 1.000 | 1.000 | 1.000 | 1.000 | 1.000 | 0.999 | 0.999 | 0.999 | 0.999 | 1.000 | 1.000 | 0.954 | 0.966 | 0.970 |
| | | TFS | | 1.000 | 1.000 | 1.000 | 0.999 | 0.999 | 0.999 | 0.999 | 0.999 | 0.999 | 0.974 | 0.976 | 0.978 | 0.916 | 0.924 | 0.910 |
| | | ZymoResearch | | 1.000 | 1.000 | 1.000 | 0.999 | 0.999 | 1.000 | 0.999 | 0.999 | 0.999 | 0.947 | 0.950 | 0.944 | 0.930 | 0.929 | 0.938 |
| **Diagenode** | 3 | EpigenDX | BC-DNA recovery | 0.976 | 0.924 | 0.834 | 0.837 | 0.895 | 0.531 | 0.749 | 0.741 | 0.695 | 0.733 | 0.673 | 0.585 | NA | NA | NA |
| | | TFS | | 0.511 | 0.503 | 0.490 | 0.531 | 0.594 | 0.440 | 0.711 | 0.687 | 0.670 | 0.441 | 0.582 | 0.540 | NA | NA | NA |
| | | ZymoResearch | | 0.757 | 0.786 | 0.794 | 0.711 | 0.671 | 0.732 | 0.596 | 0.506 | 0.450 | 0.733 | 0.818 | 0.609 | NA | NA | NA |
| | | EpigenDX | Intact BC-DNA | 0.342 | 0.212 | 0.368 | 0.319 | 0.337 | 0.382 | 0.377 | 0.336 | 0.415 | 0.484 | NA | 0.528 | NA | NA | NA |
| | | TFS | | 0.384 | 0.437 | 0.393 | 0.382 | 0.419 | 0.407 | 0.426 | 0.441 | 0.399 | NA | NA | NA | NA | NA | NA |
| | | ZymoResearch | | 0.348 | 0.387 | 0.308 | 0.294 | 0.362 | 0.354 | 0.317 | 0.432 | 0.362 | NA | NA | NA | NA | NA | NA |
| | | EpigenDX | BC efficiency | 0.974 | 0.977 | 0.980 | 0.978 | 0.980 | 0.982 | 0.999 | 0.999 | 0.999 | 0.998 | 1.000 | 0.997 | 0.977 | 0.982 | 0.987 |
| | | TFS | | 1.000 | 1.000 | 1.000 | 1.000 | 1.000 | 1.000 | 1.000 | 1.000 | 1.000 | 1.000 | 1.000 | 1.000 | 1.000 | 1.000 | 1.000 |
| | | ZymoResearch | | 1.000 | 1.000 | 1.000 | 0.989 | 0.988 | 0.990 | 0.984 | 0.985 | 0.985 | 0.981 | 0.981 | 0.983 | 0.967 | 0.970 | 0.967 |
| **Epigentek** | 9 | EpigenDX | BC-DNA recovery | 0.314 | 0.329 | 0.375 | 0.341 | 0.384 | 0.396 | 0.269 | 0.323 | 0.272 | 0.233 | 0.202 | 0.229 | NA | NA | NA |
| | | TFS | | 0.396 | 0.350 | 0.384 | 0.441 | 0.358 | 0.350 | 0.641 | 0.528 | 0.552 | NA | NA | NA | NA | NA | NA |
| | | ZymoResearch | | 0.277 | 0.266 | 0.247 | 0.229 | 0.200 | 0.203 | 0.103 | 0.095 | 0.104 | NA | NA | NA | NA | NA | NA |
| | | EpigenDX | Intact BC-DNA | 0.388 | 0.337 | 0.343 | 0.427 | 0.359 | 0.419 | 0.417 | 0.360 | 0.487 | NA | NA | NA | NA | NA | NA |
| | | TFS | | 0.343 | 0.407 | 0.364 | 0.330 | 0.391 | 0.403 | 0.338 | 0.433 | 0.489 | NA | NA | NA | NA | NA | NA |
| | | ZymoResearch | | 0.414 | 0.454 | 0.523 | 0.508 | 0.493 | 0.558 | 0.556 | 0.638 | 0.512 | NA | NA | NA | NA | NA | NA |
| | | EpigenDX | BC efficiency | 0.957 | 0.956 | 0.955 | 0.948 | 0.954 | 0.956 | 0.949 | 0.954 | 0.954 | 0.902 | 0.899 | 0.897 | 0.899 | 0.900 | 0.909 |
| | | TFS | | 0.874 | 0.892 | 0.884 | 0.872 | 0.872 | 0.873 | 0.897 | 0.878 | 0.887 | 0.814 | 0.822 | 0.851 | 0.695 | 0.821 | 0.692 |
| | | ZymoResearch | | 0.986 | 0.988 | 0.987 | 0.993 | 0.991 | 0.993 | 0.989 | 0.987 | 0.988 | 0.972 | 0.971 | 0.972 | 0.797 | 0.862 | 0.882 |
| **Promega** | 4 | EpigenDX | BC-DNA recovery | 0.810 | 0.699 | 0.691 | 0.724 | 0.913 | 0.724 | 0.817 | 0.676 | 0.706 | 0.862 | 0.740 | 0.785 | NA | NA | NA |
| | | TFS | | 0.558 | 0.655 | 0.567 | 0.511 | 0.525 | 0.455 | 0.752 | 0.757 | 0.665 | 0.468 | 0.438 | 0.535 | NA | NA | NA |
| | | ZymoResearch | | 0.716 | 0.700 | 0.664 | 0.549 | 0.602 | 0.634 | 0.391 | 0.408 | 0.361 | 0.678 | 0.746 | 0.509 | NA | NA | NA |
| | | EpigenDX | Intact BC-DNA | 0.402 | 0.330 | 0.370 | 0.431 | 0.296 | 0.465 | 0.294 | 0.386 | 0.374 | 0.279 | 0.297 | 0.292 | NA | NA | NA |
| | | TFS | | 0.445 | 0.512 | 0.396 | 0.146 | 0.214 | 0.203 | 0.299 | 0.362 | 0.351 | 0.466 | NA | NA | NA | NA | NA |
| | | ZymoResearch | | 0.397 | 0.366 | 0.395 | 0.332 | 0.353 | 0.372 | 0.458 | 0.322 | 0.520 | 0.349 | 0.333 | 0.540 | NA | NA | NA |
| | | EpigenDX | BC efficiency | 1.000 | 1.000 | 1.000 | 1.000 | 1.000 | 1.000 | 0.998 | 0.998 | 0.998 | 0.976 | 0.980 | 0.981 | 0.961 | 0.964 | 0.969 |
| | | TFS | | 1.000 | 1.000 | 1.000 | 0.604 | 0.652 | 0.626 | 0.950 | 0.950 | 0.950 | 0.996 | 0.996 | 0.997 | 0.788 | 0.791 | 0.793 |
| | | ZymoResearch | | 1.000 | 1.000 | 1.000 | 1.000 | 1.000 | 1.000 | 0.998 | 0.998 | 0.998 | 0.999 | 0.999 | 0.999 | 0.987 | 0.991 | 0.989 |
| **Qiagen** | 6 | EpigenDX | BC-DNA recovery | 0.566 | 0.674 | 0.610 | 0.348 | 0.361 | 0.336 | 0.524 | 0.590 | 0.529 | 0.386 | 0.404 | 0.364 | NA | NA | NA |
| | | TFS | | 0.490 | 0.454 | 0.448 | 0.609 | 0.582 | 0.597 | 0.508 | 0.503 | 0.463 | 0.318 | 0.356 | 0.234 | NA | NA | NA |
| | | ZymoResearch | | 0.405 | 0.370 | 0.406 | 0.361 | 0.368 | 0.389 | 0.235 | 0.273 | 0.232 | 0.367 | 0.357 | 0.362 | NA | NA | NA |
| | | EpigenDX | Intact BC-DNA | 0.386 | 0.331 | 0.385 | 0.375 | 0.324 | 0.425 | 0.456 | 0.360 | 0.362 | 0.447 | NA | 0.384 | NA | NA | NA |
| | | TFS | | 0.311 | 0.383 | 0.339 | 0.286 | 0.383 | 0.344 | 0.262 | 0.319 | 0.341 | NA | NA | NA | NA | NA | NA |
| | | ZymoResearch | | 0.386 | 0.371 | 0.376 | 0.370 | 0.425 | 0.363 | 0.344 | 0.269 | 0.330 | NA | NA | NA | NA | NA | NA |
| | | EpigenDX | BC efficiency | 0.982 | 0.983 | 0.985 | 0.999 | 0.999 | 0.998 | 1.000 | 1.000 | 1.000 | 0.996 | 0.999 | 1.000 | 0.981 | 0.985 | 0.986 |
| | | TFS | | 1.000 | 1.000 | 1.000 | 1.000 | 1.000 | 1.000 | 1.000 | 1.000 | 1.000 | 0.995 | 0.995 | 0.996 | 1.000 | 1.000 | 1.000 |
| | | ZymoResearch | | 1.000 | 1.000 | 1.000 | 0.996 | 0.995 | 0.996 | 0.982 | 0.981 | 0.982 | 0.986 | 0.985 | 0.986 | 0.959 | 0.963 | 0.963 |
| **Sigma Aldrich** | 5 | EpigenDX | BC-DNA recovery | 0.614 | 0.661 | 0.707 | 0.707 | 0.727 | 0.695 | 0.564 | 0.616 | 0.607 | 0.583 | 0.476 | 0.501 | NA | NA | NA |
| | | TFS | | 0.394 | 0.561 | 0.547 | 0.561 | 0.547 | 0.547 | 0.871 | 0.951 | 0.719 | 0.484 | 0.449 | 0.566 | NA | NA | NA |
| | | ZymoResearch | | 0.440 | 0.357 | 0.369 | 0.286 | 0.351 | 0.393 | 0.356 | 0.364 | 0.340 | 0.274 | 0.399 | 0.430 | NA | NA | NA |
| | | EpigenDX | Intact BC-DNA | 0.120 | 0.132 | 0.148 | 0.126 | 0.109 | 0.135 | 0.082 | 0.107 | 0.107 | NA | NA | NA | NA | NA | NA |
| | | TFS | | 0.139 | 0.133 | 0.133 | 0.108 | 0.108 | 0.147 | 0.137 | 0.098 | 0.119 | NA | NA | NA | NA | NA | NA |
| | | ZymoResearch | | 0.111 | 0.137 | 0.124 | 0.104 | 0.099 | 0.056 | NA | 0.081 | NA | NA | NA | NA | NA | NA | NA |
| | | EpigenDX | BC efficiency | 1.000 | 1.000 | 1.000 | 1.000 | 1.000 | 1.000 | 0.999 | 0.999 | 0.999 | 0.999 | 1.000 | 1.000 | 0.954 | 0.966 | 0.970 |
| | | TFS | | 1.000 | 1.000 | 1.000 | 1.000 | 1.000 | 1.000 | 1.000 | 1.000 | 1.000 | 0.998 | 0.998 | 0.998 | 0.955 | 0.936 | 0.945 |
| | | ZymoResearch | | 1.000 | 1.000 | 1.000 | 0.998 | 1.000 | 0.974 | 0.982 | 0.985 | 0.984 | 0.998 | 0.998 | 1.000 | 1.000 | 1.000 | 1.000 |
| **Thermo Fisher Scientific** | 2 | EpigenDX | BC-DNA recovery | 0.912 | 1.000 | 0.996 | 0.697 | 0.794 | 0.881 | 0.577 | 0.708 | 0.698 | 0.681 | 0.734 | 0.748 | NA | NA | NA |
| | | TFS | | 0.611 | 0.621 | 0.566 | 0.551 | 0.568 | 0.466 | 0.936 | 0.952 | 0.838 | 0.467 | 0.410 | 0.463 | NA | NA | NA |
| | | ZymoResearch | | 0.734 | 0.733 | 0.742 | 0.475 | 0.537 | 0.462 | 0.384 | 0.414 | 0.375 | 0.411 | 0.329 | 0.371 | NA | NA | NA |
| | | EpigenDX | Intact BC-DNA | 0.338 | 0.290 | 0.260 | 0.410 | 0.345 | 0.341 | 0.359 | 0.324 | 0.339 | NA | NA | NA | NA | NA | NA |
| | | TFS | | 0.249 | 0.266 | 0.263 | 0.389 | 0.385 | 0.416 | 0.333 | 0.260 | 0.357 | NA | NA | NA | NA | NA | NA |
| | | ZymoResearch | | 0.295 | 0.322 | 0.306 | 0.292 | 0.324 | 0.320 | NA | 0.130 | 0.183 | NA | NA | NA | NA | NA | NA |
| | | EpigenDX | BC efficiency | 0.996 | 0.983 | 0.996 | 0.999 | 0.999 | 0.999 | 0.999 | 0.999 | 1.000 | 0.997 | 0.988 | 0.997 | 0.968 | 0.969 | 0.975 |
| | | TFS | | 0.935 | 0.944 | 0.938 | 1.000 | 1.000 | 1.000 | 0.966 | 0.965 | 0.965 | 0.977 | 0.983 | 0.984 | 0.919 | 1.000 | 1.000 |
| | | ZymoResearch | | 1.000 | 1.000 | 1.000 | 0.999 | 0.999 | 0.999 | 0.928 | 0.930 | 0.931 | 0.973 | 0.975 | 0.978 | 0.942 | 0.901 | 0.952 |
| **Zymo Research** | 1 | EpigenDX | BC-DNA recovery | 1.000 | 1.000 | 1.000 | 0.999 | 1.000 | 1.000 | 1.000 | 1.000 | 1.000 | 1.000 | 0.957 | 0.992 | 1.000 | NA | 1.000 |
| | | TFS | | 0.908 | 0.985 | 0.946 | 1.000 | 1.000 | 1.000 | 1.000 | 1.000 | 1.000 | 0.687 | 0.708 | 0.721 | NA | NA | NA |
| | | ZymoResearch | | 0.973 | 0.942 | 1.000 | 1.000 | 0.902 | 1.000 | 0.670 | 0.743 | 0.676 | 0.852 | 0.849 | 0.961 | NA | NA | NA |
| | | EpigenDX | Intact BC-DNA | 0.508 | 0.449 | 0.428 | 0.556 | 0.510 | 0.489 | 0.530 | 0.646 | 0.511 | 0.631 | 0.580 | 0.578 | NA | NA | NA |
| | | TFS | | 0.456 | 0.527 | 0.480 | 0.560 | 0.607 | 0.566 | 0.580 | 0.535 | 0.554 | 0.570 | 0.550 | 0.556 | NA | NA | NA |
| | | ZymoResearch | | 0.521 | 0.483 | 0.594 | 0.482 | 0.471 | 0.456 | 0.576 | 0.607 | 0.743 | 0.595 | 0.715 | 0.626 | NA | NA | NA |
| | | EpigenDX | BC efficiency | 0.963 | 0.964 | 0.966 | 0.967 | 0.961 | 0.966 | 0.970 | 0.969 | 0.971 | 0.963 | 0.972 | 0.971 | 0.956 | 0.961 | 0.965 |
| | | TFS | | 0.910 | 0.912 | 0.919 | 0.992 | 0.993 | 0.994 | 0.926 | 0.926 | 0.929 | 0.924 | 0.924 | 0.925 | 0.854 | 0.858 | 0.850 |
| | | ZymoResearch | | 0.989 | 0.990 | 0.990 | 0.923 | 0.923 | 0.915 | 0.995 | 0.995 | 0.996 | 0.991 | 0.990 | 0.992 | 0.965 | 0.962 | 0.967 |

**S4 Table. Information on qBiCo-v2 method.** Sequences of target DNA regions, primer, probes and synthetic DNA standards per qPCR assay. Additional sequences on synthetic standards indicated in red were added for improved production and stability. Oligo changes compared to qBiCo-v1 are highlighted in blue.

| Assay | Oligo | Sequence (5'→3') | Concentration (µM) | Fluorophore | (Amplicon) length (bp) |
|---|---|---|---|---|---|
| ***hTERT* Short** | Forward primer | TGGGTTTTAGAGTTGATTTTT | 0.6 | NA | 85 |
| | Reverse primer | ACCACATTAAACAATCCCCT | 0.6 | | |
| | Probe | ATTTAGGTATAGGGGATTTGGTTTTAGT | 0.4 | HEX™ | |
| | Synthetic standard | GAATTACGGAGATGGTTAGGAGTGGGTTTTAGAGTTGATTTTTGTGAATTA<br>AATTAAAATTAGGTATAGGGGATTTGGTTTTAGTATAGGGGATTGTTTAAT<br>GTGGTTTTTTTTAAGGGCGTTTT | NA | NA | |
| ***TPT1* Long** | Forward primer | AAGGTGTTTTAATTTAGTGA | 0.6 | NA | 222 |
| | Reverse primer | CAATAAAACAACCATACCATCTA | 0.6 | | |
| | Probe | TGAAGTTTTGATTTTTGTGGTGGT | 0.4 | FAM™ | |
| | Synthetic standard | CGCCAAGGTGTTTTAATTTAGTGAATAGATGTTTATGATAAGTGAGTATTA<br>GAGTTTTTGGGTATTGAAGTTTTGATTTTTGTGGTGGTTTAAATTTTTTTTT<br>GTATTGTAGTTTGTTTTGAATGGAATGTATTTGTATGTAATAGTTTAATTTTA<br>GGTATTTTGTTTGTTTTTTAAGTTTTTTATTGGTGAAAATATGAATTTAGATG<br>GTATGGTTGTTTTATTGCCGC | NA | NA | 230 |
| ***LINE1* Converted** | Forward primer | TTTTGAGTTAGGTGTGGGATATA | 0.05 | NA | 148 |
| | Reverse primer | AAAATCAAAAAATTCCCTTTC | 0.05 | | |
| | Probe | TGGGAGTGATTTGATTTTTTAGGTGY | 0.02 | TEX615™ | |
| | Synthetic standard | GGGCGTAGGATTTTTTGAGTTAGGTGTGGGATATAATTTTTTGGTGTGTTG<br>TTTTTTAAGTTTTTTGGAAAAGTGTAGTATTTAGGGTGGGAGTGATTTGATT<br>TTTTAGGTGTTGTTTGTTATTTTTTTTTTTGATTAGGAAAGGGAATTTTTTGA<br>TTTTTTGTATTTTTCGAGTGAGGTAATGTTTCG | NA | NA | 189 |
| ***LINE1* Genomic** | Forward primer | GAGCCAGGTGTGGGATATAA | 0.1 | NA | 119 |
| | Reverse primer | TCAAAG---GGGGTCACAGA | 0.1 | | |
| | Probe | TGGGAGTGACCCGATTTTCCAGGTG | 0.05 | Cy5™ | |
| | Synthetic standard | CTCTGAGCCAGGTGTGGGATATAATCTCCTGGTGTGCTGTTTTTTAAGCCC<br>TTTGGAAAAGTGCAGTATTTAGGGTGGGAGTGACCCGATTTTCCAGGTGC<br>CGTCTGTCACCCCTTTCTTTGACTAGG | NA | NA | 128 |
| **Synthetic IPC** | Forward primer | GGTGATTTTTAATAATTTTTGAGG | 0.3 µM | NA | 99 |
| | Reverse primer | ACTTTCAACCCATTACCTCAT | 0.3 µM | | |
| | Probe | AGTGTTCGTATAGGTTTTTTTTTTTAGT | 0.2 µM | Cy5.5™ | |
| | Synthetic standard | CGTAGAGTGTAATTGTATTAGGGTGATTTTTAATAATTTTTGAGGGAGTATT<br>GGTATAGGCGATAGAGTGTTCGTATAGGTTTTTTTTTTTAGTGGATGATGA<br>GGTAATGGGTTGAAAGTTATATCGTGG | NA | NA | 130 |

**S5 Table. Adjustment of synthetic DNA standard preparation between qBiCo-v1 and -v2 methods.**
gBlocks™: Gene fragments ordered from IDT.

| **qBiCo method** | **Dilution factor** | **Synthetic standards** | **Amounts** |
|---|---|---|---|
| v1 | *Original gBlock stock standards* | | |
| | 10X | STD1 | 3ul gBlock stock mix + 27 ul water |
| | 10X | STD2 | 3ul STD1 + 27 ul water |
| | 10X | STD3 | 3ul STD2 + 27 ul water |
| | 10X | STD4 | 3ul STD3 + 27 ul water |
| | 10X | STD5 | 5ul STD4 + 45 ul water |
| | *Working qBiCo standards* | | |
| | 4X | STDA | 4ul gBlock mix+ 12 ul water |
| | 3X | STDB | 4ul STDA+ 8 ul water |
| | 3X | STDC | 4ul STDB+ 8 ul water |
| | 3X | STDD | 4ul STDC+ 8 ul water |
| | 3X | STDE | 4ul STDD+ 8 ul water |
| v2 | *Original gBlock stock standards* | | |
| | 10X | STD1 | 10ul gBlock stock mix + 90ul TE buffer |
| | 10X | STD2 | 10ul STD1 + 90ul TE |
| | 10X | STD3 | 10ul STD2 + 90ul TE |
| | 10X | STD4 | 10ul STD3 + 90ul TE |
| | 10X | STD5 | 10ul STD4 + 90ul TE |
| | 10X | STD6 | 10ul STD5 + 90ul TE |
| | *Working qBiCo standards* | | |
| | 2X | STDA | 10ul gBlock mix+ 10ul TE |
| | 3X | STDB | 5ul STDA+ 10ul TE |
| | 3X | STDC | 5ul STDB+ 10ul TE |
| | 3X | STDD | 5ul STDC+ 10ul TE |
| | 3X | STDE | 5ul STDD+ 10ul TE |

**S6 Table. Overview of qPCR platforms and adjustments made for transferring qBiCo-v2 assay.** More details on the technology transfer can be found in the Methods. IPC: Internal positive control.

| Instrument | Manufacturer | Well set-up | Multiplex capabilities | Fluorescent dyes used/replaced | Reaction volume (µl) | Instrument software used for analysis |
|---|---|---|---|---|---|---|
| CFX96 | Bio-Rad | 96 | 5-plex | FAM, HEX, TEX615, Cy5, Cy5.5 | 10 | CFX Manager Software BioRad 96-well |
| CFX384 | Bio-Rad | 384 | 4-plex (excl. IPC) | FAM, HEX, TEX615, Cy5 | 10 | CFX Manager Software BioRad 384-well |
| QS5 | Applied Biosystems | 96 | 5-plex | FAM, VIC, ABY, Cy5, JUN | 10 | QuantStudio 3 and 5 Real-Time PCR System Software |
| QS7 Flex | Applied Biosystems | 384 | 5-plex | FAM, VIC, ABY, Cy5, JUN | 10 | QuantStudio 7 Flex Real-Time PCR System, 384-well |
| 7500 Fast | Applied Biosystems | 96 | 5-plex | FAM, VIC, ABY, Cy5, JUN | 10 | 7500 Software v2.3 96-well |
| Rotorgene Q | Qiagen | 72 | 5-plex | FAM, HEX, TEX615, Cy5, Cy5.5 | 20 | Rotor-Gene Q Series Software 72-tubes |

**S7 Table. Data obtained by transferring qBiCo-v2 assay to different qPCR instruments from various manufacturers.** Data include all technical replicates of the synthetic standards per qBiCo assay, as measured in the optimized transfer qPCR run. Information on the make-up of the synthetic standards is provided in the Methods. Cq: Quantification cycle; eRFU: End-point relative fluorescent units.

| qPCR instrument | Synthetic standard | Replicate | Cq | | | | | eRFU | | | | |
|---|---|---|---|---|---|---|---|---|---|---|---|---|
| | | | ***hTERT*** **Short** | ***TPT1*** **Long** | ***LINE1*** **Converted** | ***LINE1*** **Genomic** | **IPC** | ***hTERT*** **Short** | ***TPT1*** **Long** | ***LINE1*** **Converted** | ***LINE1*** **Genomic** | **IPC** |
| | | 1 | 25.55 | 24.96 | 17.63 | 23.10 | 29.19 | 4049 | 2661 | 335 | 752 | 687 |
| | StdA | 2 | 25.17 | 24.42 | 17.59 | 22.28 | 28.71 | 4369 | 2895 | 348 | 809 | 739 |
| | | 3 | 25.36 | 24.69 | 17.17 | 22.50 | 28.68 | 4257 | 2850 | 351 | 776 | 779 |
| | | 1 | 27.21 | 26.33 | 19.49 | 24.26 | 28.55 | 2937 | 1904 | 313 | 618 | 776 |
| | StdB | 2 | 27.02 | 26.20 | 19.07 | 24.24 | 28.89 | 3209 | 2138 | 341 | 670 | 757 |
| | | 3 | 27.12 | 26.31 | 19.02 | 24.04 | 28.56 | 3194 | 2085 | 365 | 669 | 840 |
| | | 1 | 28.77 | 28.26 | 20.95 | 25.86 | 28.22 | 1988 | 1183 | 298 | 518 | 871 |
| **CFX96** | StdC | 2 | 28.61 | 28.13 | 20.87 | 25.86 | 28.48 | 2112 | 1300 | 305 | 536 | 864 |
| | | 3 | 28.67 | 27.93 | 20.90 | 25.90 | 28.37 | 2172 | 1415 | 330 | 539 | 858 |
| | | 1 | 30.61 | 30.04 | 22.83 | 27.67 | 28.16 | 1101 | 702 | 293 | 388 | 993 |
| | StdD | 2 | 30.68 | 30.21 | 23.09 | 27.86 | 28.28 | 1075 | 639 | 304 | 377 | 969 |
| | | 3 | 30.49 | 30.34 | 22.79 | 27.58 | 28.26 | 1164 | 595 | 316 | 396 | 967 |
| | | 1 | 31.68 | 30.65 | 24.17 | 28.71 | 28.03 | 697 | 511 | 279 | 309 | 1087 |
| | StdE | 2 | 32.15 | 31.24 | 24.20 | 28.63 | 27.92 | 554 | 406 | 271 | 311 | 1130 |
| | | 3 | 31.52 | 31.66 | 24.11 | 28.38 | 27.55 | 766 | 296 | 311 | 334 | 1209 |
| | | 1 | 26.15 | 24.01 | 19.57 | 23.43 | NA | 1473 | 1383 | 139 | 792 | NA |
| | StdA | 2 | 25.75 | 24.07 | 20.26 | 23.25 | NA | 1562 | 1371 | 127 | 817 | NA |
| | | 3 | 26.07 | 24.20 | 20.04 | 23.41 | NA | 1501 | 1305 | 137 | 777 | NA |
| | | 1 | 27.90 | 25.97 | 20.89 | 25.15 | NA | 954 | 960 | 137 | 654 | NA |
| | StdB | 2 | 27.77 | 25.83 | 21.79 | 25.23 | NA | 967 | 932 | 134 | 625 | NA |
| | | 3 | 27.88 | 26.05 | 21.26 | 25.32 | NA | 973 | 981 | 135 | 644 | NA |
| | | 1 | 29.42 | 27.41 | 22.90 | 26.58 | NA | 611 | 696 | 126 | 595 | NA |
| **CFX384** | StdC | 2 | 29.19 | 27.58 | 23.47 | 26.49 | NA | 640 | 680 | 119 | 586 | NA |
| | | 3 | 29.68 | 27.61 | 23.26 | 26.56 | NA | 519 | 629 | 114 | 557 | NA |
| | | 1 | 31.20 | 29.12 | 24.73 | 27.58 | NA | 287 | 434 | 123 | 517 | NA |
| | StdD | 2 | 30.84 | 29.60 | 25.02 | 27.85 | NA | 323 | 324 | 118 | 475 | NA |
| | | 3 | 31.05 | 29.23 | 24.85 | 28.13 | NA | 302 | 391 | 107 | 428 | NA |
| | | 1 | 31.36 | 30.65 | 26.40 | 28.90 | NA | 256 | 229 | 104 | 393 | NA |
| | StdE | 2 | 32.61 | 30.26 | 26.30 | 28.61 | NA | 127 | 274 | 104 | 422 | NA |
| | | 3 | 32.52 | 30.11 | 26.42 | 28.52 | NA | 134 | 287 | 108 | 411 | NA |
| | | 1 | 24.89 | 25.95 | 18.76 | 23.17 | 27.21 | 214388 | 81731 | 54839 | 26593 | 85561 |
| | StdA | 2 | 24.81 | 24.37 | 18.01 | 22.33 | 26.89 | 207083 | 100842 | 52258 | 27937 | 82778 |
| | | 3 | 25.15 | 25.12 | 16.99 | 22.65 | 27.22 | 196220 | 91675 | 52532 | 25622 | 80989 |
| | | 1 | 26.57 | 26.93 | 18.87 | 24.27 | 25.89 | 153939 | 73314 | 53099 | 25509 | 97816 |
| | StdB | 2 | 26.13 | 27.11 | 19.31 | 24.37 | 27.14 | 179621 | 61908 | 48209 | 23088 | 87937 |
| | | 3 | 26.84 | 26.99 | 19.82 | 24.07 | 26.91 | 151074 | 65412 | 43711 | 22567 | 85202 |
| | | 1 | 27.61 | 29.21 | 22.12 | 26.38 | 27.31 | 135471 | 38516 | 46215 | 17969 | 87895 |
| **QS5** | StdC | 2 | 28.66 | 30.03 | 22.47 | 26.35 | 27.02 | 109689 | 25726 | 40252 | 18484 | 92927 |
| | | 3 | 29.44 | 28.97 | 22.27 | 26.09 | 27.49 | 90110 | 40865 | 38084 | 18622 | 89221 |
| | | 1 | 28.02 | 30.31 | 23.33 | 28.84 | 26.79 | 121756 | 23164 | 41768 | 11531 | 89413 |
| | StdD | 2 | 27.19 | 31.18 | 23.67 | 27.77 | 27.40 | 137790 | 16516 | 40287 | 12548 | 82574 |
| | | 3 | 31.28 | 31.26 | 23.59 | 27.15 | 27.63 | 48976 | 19347 | 35341 | 14327 | 90517 |
| | | 1 | 28.97 | 33.06 | 24.48 | 29.02 | 26.61 | 92246 | 6890 | 37079 | 10164 | 89330 |
| | StdE | 2 | 31.83 | 33.10 | 25.48 | 28.55 | 27.26 | 34696 | 7826 | 29411 | 10566 | 88630 |
| | | 3 | 31.80 | 32.50 | 25.22 | 29.73 | 26.63 | 37817 | 10879 | 31206 | 8546 | 105335 |
| | | 1 | 25.20 | 24.99 | 22.38 | 23.50 | 27.39 | 595688 | 69090 | 62547 | 74917 | 281510 |
| | StdA | 2 | 25.16 | 25.02 | 22.58 | 23.54 | 27.43 | 594212 | 64992 | 57758 | 73024 | 252357 |
| | | 3 | 25.34 | 25.40 | 22.42 | 23.39 | 27.63 | 591778 | 58937 | 57129 | 74654 | 269070 |
| | | 1 | 27.31 | 27.21 | 24.57 | 25.56 | 27.77 | 416272 | 41340 | 45854 | 59582 | 294102 |
| | StdB | 2 | 27.29 | 26.95 | 24.62 | 25.52 | 27.79 | 453032 | 48298 | 52440 | 69469 | 314369 |
| | | 3 | 27.32 | 26.87 | 24.61 | 25.44 | 27.60 | 439180 | 42012 | 48633 | 66563 | 324435 |
| | | 1 | 28.68 | 28.58 | 26.16 | 26.80 | 27.81 | 329547 | 33185 | 38571 | 57471 | 340930 |
| **QS7 Flex** | StdC | 2 | 28.46 | 28.29 | 25.93 | 26.48 | 27.57 | 351364 | 36788 | 44595 | 63820 | 365017 |
| | | 3 | 28.92 | 28.19 | 26.13 | 26.89 | 27.69 | 333612 | 39388 | 43271 | 64776 | 373739 |
| | | 1 | 30.49 | 31.20 | 28.21 | 28.55 | 28.06 | 197544 | 18681 | 29790 | 47640 | 346505 |
| | StdD | 2 | 30.32 | 31.22 | 28.31 | 28.45 | 27.76 | 197796 | 16761 | 28177 | 45304 | 363521 |
| | | 3 | 29.71 | 30.24 | 27.85 | 28.37 | 27.91 | 262983 | 22588 | 31442 | 51843 | 371357 |
| | | 1 | 31.28 | NA | 29.85 | 29.25 | 28.21 | 161326 | 6771 | 21660 | 43559 | 344891 |
| | StdE | 2 | 31.92 | 34.62 | 30.77 | 30.26 | 28.09 | 112762 | 9338 | 17693 | 34250 | 349310 |
| | | 3 | 31.52 | NA | 30.27 | 29.78 | 27.72 | 128125 | 5544 | 17578 | 39145 | 386688 |

| | | | | | | | | | | | | |
|---|---|---|---|---|---|---|---|---|---|---|---|---|
| **7500 Fast** | StdA | 1 | 26.29 | 26.09 | 19.30 | 24.26 | 28.17 | 967415 | 183145 | 172913 | 123841 | 432542 |
| | | 2 | 26.20 | 28.56 | 19.58 | 24.99 | 28.19 | 942715 | 77858 | 156046 | 109478 | 433949 |
| | | 3 | 26.94 | 28.24 | 19.61 | 25.44 | 27.85 | 796631 | 85417 | 140694 | 99079 | 470216 |
| | StdB | 1 | 29.15 | 29.02 | 20.56 | 27.18 | 28.37 | 600488 | 88968 | 143755 | 85168 | 454741 |
| | | 2 | 27.74 | 29.52 | 20.80 | 26.16 | 27.57 | 773175 | 56478 | 155221 | 96512 | 513948 |
| | | 3 | 29.05 | 29.30 | 20.96 | 26.93 | 28.29 | 580956 | 75634 | 134032 | 86703 | 476964 |
| | StdC | 1 | 30.29 | 33.97 | 22.50 | 28.14 | 28.56 | 323130 | 15902 | 141125 | 56653 | 384021 |
| | | 2 | 29.17 | 32.05 | 22.44 | 28.33 | 27.68 | 578312 | 26396 | 139918 | 72967 | 532338 |
| | | 3 | 28.87 | 30.40 | 22.92 | 27.96 | 27.68 | 575107 | 46295 | 132649 | 75831 | 527170 |
| | StdD | 1 | 31.10 | NA | 24.92 | 30.44 | 26.40 | 321967 | 8229 | 108811 | 39971 | 643553 |
| | | 2 | 30.76 | 34.44 | 24.23 | 30.79 | 28.08 | 370700 | 15853 | 115858 | 37675 | 513912 |
| | | 3 | 30.77 | 27.25 | 24.75 | 29.19 | 27.80 | 304556 | 181784 | 108831 | 61069 | 549001 |
| | StdE | 1 | 32.26 | NA | 26.08 | 31.00 | 28.73 | 196255 | 10790 | 97693 | 41558 | 453676 |
| | | 2 | 31.97 | NA | 25.42 | 30.68 | 27.91 | 175562 | 13525 | 105386 | 43205 | 546251 |
| | | 3 | 30.98 | 34.67 | 25.51 | 32.95 | 27.05 | 274172 | 14564 | 106874 | 20940 | 591536 |
| **Rotorgene Q** | StdA | 1 | 23.79 | 24.90 | 18.96 | 20.79 | 29.70 | 0.618 | 0.529 | 0.260 | 0.846 | 0.093 |
| | | 2 | 23.52 | 24.42 | 17.97 | 20.61 | 29.16 | 0.643 | 0.551 | 0.295 | 0.880 | 0.102 |
| | | 3 | 23.77 | 24.70 | 18.60 | 20.90 | 29.75 | 0.640 | 0.539 | 0.276 | 0.871 | 0.096 |
| | StdB | 1 | 25.26 | 26.34 | 19.97 | 22.46 | 29.49 | 0.532 | 0.439 | 0.275 | 0.822 | 0.101 |
| | | 2 | 25.29 | 26.29 | 19.66 | 22.37 | 29.43 | 0.532 | 0.438 | 0.281 | 0.820 | 0.101 |
| | | 3 | 25.47 | 26.11 | 20.23 | 22.48 | 29.51 | 0.522 | 0.434 | 0.263 | 0.800 | 0.102 |
| | StdC | 1 | 27.02 | 28.40 | 22.21 | 24.23 | 29.32 | 0.395 | 0.300 | 0.270 | 0.727 | 0.102 |
| | | 2 | 27.18 | 28.32 | 21.95 | 24.23 | 29.46 | 0.406 | 0.313 | 0.272 | 0.723 | 0.105 |
| | | 3 | 27.09 | 28.09 | 22.17 | 24.28 | 29.25 | 0.410 | 0.297 | 0.267 | 0.724 | 0.107 |
| | StdD | 1 | 28.92 | 30.34 | 23.85 | 25.98 | 29.51 | 0.284 | 0.197 | 0.262 | 0.654 | 0.107 |
| | | 2 | 28.70 | 30.64 | 23.49 | 25.93 | 29.75 | 0.308 | 0.184 | 0.264 | 0.641 | 0.103 |
| | | 3 | 29.30 | 29.97 | 24.16 | 25.90 | 29.42 | 0.266 | 0.203 | 0.250 | 0.616 | 0.104 |
| | StdE | 1 | 29.53 | 31.64 | 23.73 | 26.45 | 29.57 | 0.229 | 0.104 | 0.280 | 0.612 | 0.104 |
| | | 2 | 30.09 | 32.27 | 25.33 | 27.15 | 29.47 | 0.214 | 0.112 | 0.259 | 0.570 | 0.108 |
| | | 3 | 29.96 | 31.64 | 24.95 | 26.89 | 29.23 | 0.204 | 0.125 | 0.265 | 0.573 | 0.110 |

**S8 Table. Data obtained by validating qBiCo-v2 assay across qPCR platforms.** Data include all technical replicates of all samples per qBiCo assay among different qPCR experiments, in terms of Cq values, measured fragment copies and qBiCo indices. BC: Bisulfite conversion; Cq: Quantification cycle; IPC: Internal positive control.

| qPCR instrument | Sample source | Sample ID | Genome DNA input (ng) | qPCR run | Replicate | Cq | | | | | No copies | | | | qBiCo indices | | |
|---|---|---|---|---|---|---|---|---|---|---|---|---|---|---|---|---|---|
| | | | | | | *TPT1* Long | *hTERT* Short | *LINE1* Converted | *LINE1* Genomic | IPC | *TPT1* Long | *hTERT* Short | *LINE1* Converted | *LINE1* Genomic | BC (%) | BC-DNA concentration (ng/µl) | BC-DNA fragmentation |
| CFX96 | Human whole blood | BB_1A | 50 | 1 | 1 | 26.83 | 27.33 | 19.93 | 24.43 | 29.76 | 1329 | 1685 | 80863 | 3342 | 97.98 | 5.53 | 0.81 |
| | | | | | 2 | 27.02 | 27.24 | 19.82 | 24.34 | 29.54 | 1175 | 1790 | 86784 | 3563 | 97.99 | 5.88 | 0.98 |
| | | | | | 3 | 26.51 | 27.26 | 20.16 | 24.4 | 29.88 | 1637 | 1766 | 69758 | 3414 | 97.61 | 5.80 | 0.69 |
| | | | | | 4 | 26.03 | 26.84 | 19.87 | 24.53 | 29.9 | 2236 | 2343 | 84041 | 3112 | 98.18 | 7.69 | 0.66 |
| | | | | | 5 | 26.39 | 26.8 | 19.52 | 24.38 | 29.13 | 1769 | 2407 | 105227 | 3463 | 98.38 | 7.90 | 0.86 |
| | | | | | 6 | 26.08 | 26.94 | 19.46 | 24.35 | 29.88 | 2165 | 2190 | 109361 | 3538 | 98.41 | 7.19 | 0.64 |
| | | | | | 7 | 26.19 | 27 | 20.05 | 24.5 | 29.73 | 2015 | 2104 | 74865 | 3179 | 97.92 | 6.91 | 0.66 |
| | | | | | 8 | 26.37 | 27.05 | 19.85 | 24.6 | 29.71 | 1793 | 2034 | 85127 | 2961 | 98.29 | 6.68 | 0.72 |
| | | BB_1A_UV120 | | | 1 | NA | 28.26 | 22.99 | 22.41 | 30.05 | NA | 901 | 11327 | 14090 | 61.65 | 2.96 | NA |
| | | | | | 2 | NA | 28.37 | 22.89 | 22.28 | 29.68 | NA | 836 | 12079 | 15458 | 60.98 | 2.75 | NA |
| | | | | | 3 | NA | 28.38 | 22.95 | 22.78 | 30.36 | NA | 831 | 11622 | 10826 | 68.23 | 2.73 | 0.00 |
| | | BB_1A_UV30 | | | 1 | 28.95 | 27.16 | 20.46 | 22.61 | 30.18 | 335 | 1889 | 57531 | 12219 | 90.40 | 6.20 | 3.80 |
| | | | | | 2 | 29.36 | 27.88 | 20.97 | 23.93 | 29.54 | 257 | 1163 | 41460 | 4772 | 94.56 | 3.82 | 3.08 |
| | | | | | 3 | 29.08 | 27.69 | 21.11 | 23.54 | 29.67 | 308 | 1322 | 37895 | 6300 | 92.33 | 4.34 | 2.90 |
| | | BB_1A_UV60 | | | 1 | NA | 27.25 | 21.47 | 22.44 | 30.24 | NA | 1778 | 30071 | 13792 | 81.35 | 5.84 | NA |
| | | | | | 2 | NA | 27.85 | 21.3 | 23.12 | 30.04 | NA | 1187 | 33541 | 8497 | 88.76 | 3.90 | 0.00 |
| | | | | | 3 | 32.8 | 27.7 | 21.94 | 23.39 | 29.74 | 27 | 1313 | 22235 | 7010 | 86.38 | 4.31 | 35.27 |
| | | BB_1B | | | 1 | 27.66 | 28.19 | 20.63 | 25.3 | 29.59 | 775 | 944 | 51580 | 1798 | 98.29 | 3.10 | 0.80 |
| | | | | | 2 | 27.92 | 28.23 | 20.62 | 25.25 | 29.83 | 654 | 919 | 51912 | 1863 | 98.24 | 3.02 | 0.92 |
| | | | | | 3 | 27.97 | 28.27 | 20.65 | 25.2 | 29.74 | 633 | 895 | 50921 | 1931 | 98.14 | 2.94 | 0.93 |
| | | BB_1C | | | 1 | 28.41 | 29.29 | 21.98 | 25.93 | 29.52 | 476 | 450 | 21671 | 1148 | 97.42 | 1.48 | 0.63 |
| | | | | | 2 | 28.3 | 29.32 | 21.97 | 25.84 | 29.4 | 511 | 441 | 21811 | 1224 | 97.27 | 1.45 | 0.57 |
| | | | | | 3 | 28.72 | 29.19 | 21.75 | 25.82 | 29.64 | 389 | 482 | 25121 | 1241 | 97.59 | 1.58 | 0.83 |
| | | BB_1D | | | 1 | 30.88 | 30.26 | 22.68 | 26.75 | 29.5 | 96 | 234 | 13823 | 640 | 97.74 | 0.77 | 1.73 |
| | | | | | 2 | 30.41 | 30.2 | 22.68 | 26.73 | 30 | 130 | 244 | 13823 | 649 | 97.71 | 0.80 | 1.31 |
| | | | | | 3 | 29.51 | 29.99 | 22.79 | 26.97 | 29.87 | 233 | 281 | 12880 | 547 | 97.92 | 0.92 | 0.82 |
| | | BB_1E | | | 1 | 31.58 | 31.46 | 24.04 | 27.37 | 29.06 | 61 | 104 | 5771 | 412 | 96.56 | 0.34 | 1.23 |
| | | | | | 2 | 31.09 | 30.53 | 23.74 | 27.81 | 29.73 | 83 | 195 | 6997 | 301 | 97.90 | 0.64 | 1.66 |
| | | | | | 3 | 30.79 | 30.69 | 24.18 | 27.88 | 30.1 | 101 | 175 | 5274 | 286 | 97.36 | 0.58 | 1.22 |
| | | BB_1F | | | 1 | 32.72 | 32.23 | 25.11 | 28.57 | 29.99 | 29 | 62 | 2902 | 175 | 97.07 | 0.20 | 1.58 |
| | | | | | 2 | 30.88 | 31.94 | 25.3 | 28.44 | 29.62 | 96 | 76 | 2569 | 192 | 96.40 | 0.25 | 0.56 |
| | | | | | 3 | 31.79 | 31.77 | 25.12 | 28.22 | 29.57 | 53 | 85 | 2884 | 225 | 96.25 | 0.28 | 1.15 |
| | | BB_1G | | | 1 | 34.44 | 32.86 | 26.28 | 28.95 | 29.27 | 9 | 41 | 1369 | 134 | 95.35 | 0.13 | 3.30 |
| | | | | | 2 | 32.56 | 33.13 | 26.24 | 29.01 | 29.06 | 32 | 34 | 1404 | 128 | 95.64 | 0.11 | 0.78 |
| | | | | | 3 | 32.22 | 33.06 | 26.37 | 28.96 | 29.39 | 40 | 36 | 1292 | 133 | 95.12 | 0.12 | 0.65 |
| | | BB_1H | | | 1 | NA | 32.75 | 28.08 | 30.11 | 29.42 | NA | 44 | 431 | 58 | 93.65 | 0.14 | NA |
| | | | | | 2 | NA | 34.11 | 28.05 | 28.98 | 28.79 | NA | 18 | 439 | 131 | 87.05 | 0.06 | NA |
| | | | | | 3 | NA | 34.57 | 28.09 | 29.47 | 29.23 | NA | 13 | 428 | 92 | 90.28 | 0.04 | NA |
| | | BB_1I | | | 1 | NA | 33.77 | 30.05 | 29.87 | 29.64 | NA | 22 | 122 | 69 | 77.81 | 0.07 | NA |
| | | | | | 2 | NA | 34 | 30.34 | 29.61 | 29.87 | NA | 19 | 101 | 83 | 70.74 | 0.06 | NA |
| | | | | | 3 | NA | NA | 30.29 | 29.71 | 28.93 | NA | NA | 104 | 78 | 72.83 | NA | NA |
| | | BB_2A | 5 | | 1 | 30.89 | 31.22 | 24.15 | 26.45 | 29.6 | 95 | 123 | 5377 | 793 | 93.14 | 0.40 | 0.91 |
| | | | | | 2 | 31.25 | 30.73 | 24.12 | 26.32 | 29.77 | 75 | 171 | 5482 | 869 | 92.65 | 0.56 | 1.62 |
| | | | | | 3 | 31.28 | 31.12 | 23.58 | 26.49 | 29.55 | 74 | 131 | 7754 | 770 | 95.27 | 0.43 | 1.27 |
| | | | | | 4 | 32.55 | 30.86 | 23.67 | 26.75 | 29.29 | 32 | 156 | 7319 | 640 | 95.81 | 0.51 | 3.55 |
| | | | | | 5 | 33.92 | 30.64 | 24.01 | 26.29 | 28.89 | 13 | 181 | 5883 | 888 | 92.98 | 0.60 | 10.36 |
| | Negative control | NC | NA | | 1 | NA | 34.41 | NA | 30.07 | 29.34 | NA | 14 | NA | 60 | NA | 0.05 | NA |
| | | | | | 2 | NA | NA | NA | NA | 33.46 | NA | NA | NA | NA | NA | NA | NA |
| | | | | | 3 | NA | 33.9 | NA | 30.58 | 29.91 | NA | 20 | NA | 42 | NA | 0.07 | NA |
| | *Gallus gallus domesticus* | NHS_1 | 200 | | 1 | NA | NA | NA | 30.13 | 32.17 | NA | NA | NA | 58 | NA | NA | NA |
| | | | | | 2 | NA | 33.37 | NA | 30.65 | 32.57 | NA | 29 | NA | 40 | NA | 0.09 | NA |
| | *Sus scrofa domesticus* | NHS_2 | 80 | | 1 | NA | 33.86 | NA | 29.47 | 30.02 | NA | 21 | NA | 92 | NA | 0.07 | NA |
| | | | | | 2 | NA | 32.77 | NA | 29.38 | 29.88 | NA | 43 | NA | 98 | NA | 0.14 | NA |
| | *Felis catus* | NHS_3 | | | 1 | NA | 32.87 | NA | 30.16 | 30 | NA | 40 | NA | 56 | NA | 0.13 | NA |
| | | | | | 2 | NA | 33.95 | NA | 29.73 | 29.52 | NA | 20 | NA | 77 | NA | 0.06 | NA |
| | *Canis lupus familiaris* | NHS_4 | | | 1 | NA | 33.04 | NA | 30.22 | 30.02 | NA | 36 | NA | 54 | NA | 0.12 | NA |
| | | | | | 2 | NA | 33.19 | NA | 30.06 | 29.21 | NA | 33 | NA | 61 | NA | 0.11 | NA |
| | *Rattus* | NHS_5 | | | 1 | NA | 34.68 | NA | 29.26 | 29.26 | NA | 12 | NA | 107 | NA | 0.04 | NA |
| | | | | | 2 | NA | 34.06 | NA | 28.51 | 29.29 | NA | 18 | NA | 183 | NA | 0.06 | NA |
| | *Mus musculus* | NHS_6 | | | 1 | NA | 33.11 | NA | 29.77 | 29.42 | NA | 34 | NA | 74 | NA | 0.11 | NA |
| | | | | | 2 | NA | 33.94 | NA | 30.17 | 29.36 | NA | 20 | NA | 56 | NA | 0.06 | NA |
| | *Bos taurus* | NHS_7 | | | 1 | NA | NA | NA | 29.82 | 29.33 | NA | NA | NA | 72 | NA | NA | NA |
| | | | | | 2 | NA | 33.01 | NA | 30.16 | 29.4 | NA | 37 | NA | 56 | NA | 0.12 | NA |
| | *Rhesus* | NHS_8 | | | 1 | 29.17 | 28.49 | 24.54 | 28.73 | 29.54 | 290 | 771 | 4185 | 156 | 98.17 | 2.53 | 1.80 |
| | | | | | 2 | 28.86 | 28.55 | 25.28 | 29.15 | 29.78 | 355 | 741 | 2602 | 116 | 97.82 | 2.43 | 1.40 |
| | Positive control | PC | NA | | 1 | 27.66 | 31.62 | 18.03 | 30.37 | 29.62 | 775 | 94 | 274018 | 49 | 99.99 | 0.31 | 0.08 |
| | | | | | 2 | 27.22 | 31.2 | 17.7 | 29.63 | 30.09 | 1032 | 124 | 338716 | 82 | 99.99 | 0.41 | 0.08 |
| | | | | | 3 | 27.39 | 31.15 | 18.15 | 28.06 | 25.58 | 924 | 129 | 253690 | 252 | 99.95 | 0.42 | 0.09 |
| | | BB_1A | | | 1 | 26.68 | 27.68 | 19.88 | 24.56 | 30.6 | 1465 | 1331 | 83503 | 3046 | 98.21 | 4.37 | 0.58 |
| | | | | | 2 | 26.9 | 27.88 | 19.75 | 24.78 | 30.88 | 1270 | 1163 | 90775 | 2604 | 98.59 | 3.82 | 0.59 |

| | | | | | | | | | | | | | | | | |
|---|---|---|---|---|---|---|---|---|---|---|---|---|---|---|---|---|
| Human whole blood | | 50 | 2 | 3 | 26.91 | 27.8 | 19.21 | 24.55 | 30.27 | 1262 | 1228 | 128412 | 3068 | 98.82 | 4.03 | 0.63 |
| | BB_1B | | | 1 | 27.74 | 28.56 | 20.43 | 25.53 | 29.81 | 736 | 736 | 58650 | 1526 | 98.72 | 2.42 | 0.66 |
| | | | | 2 | 27.57 | 28.63 | 20.79 | 25.48 | 30.12 | 822 | 702 | 46542 | 1582 | 98.33 | 2.31 | 0.56 |
| | | | | 3 | 27.78 | 28.58 | 20.38 | 25.62 | 30.64 | 717 | 726 | 60565 | 1432 | 98.83 | 2.38 | 0.66 |
| | BB_1C | | | 1 | 28.22 | 29.68 | 21.68 | 26.29 | 28.86 | 538 | 346 | 26277 | 888 | 98.34 | 1.14 | 0.43 |
| | | | | 2 | 28.35 | 29.42 | 21.41 | 26.3 | 29.77 | 495 | 412 | 31253 | 882 | 98.61 | 1.35 | 0.55 |
| | | | | 3 | 28.45 | 29.41 | 21.66 | 26.16 | 30.06 | 464 | 415 | 26616 | 974 | 98.20 | 1.36 | 0.60 |
| | BB_1D | | | 1 | 29.47 | 30.51 | 22.48 | 27.08 | 30.2 | 239 | 198 | 15718 | 506 | 98.42 | 0.65 | 0.57 |
| | | | | 2 | 29.67 | 30.23 | 22.57 | 27.21 | 30.59 | 210 | 239 | 14835 | 461 | 98.47 | 0.79 | 0.78 |
| | | | | 3 | 30.03 | 30.38 | 22.19 | 27.03 | 30.25 | 166 | 216 | 18937 | 524 | 98.63 | 0.71 | 0.90 |
| | BB_1E | | | 1 | 30.56 | 31.8 | 23.42 | 28.05 | 29.95 | 118 | 83 | 8594 | 254 | 98.55 | 0.27 | 0.49 |
| | | | | 2 | 30.71 | 31.73 | 23.52 | 28.33 | 30.29 | 107 | 87 | 8059 | 208 | 98.73 | 0.29 | 0.57 |
| | | | | 3 | 30.63 | 31.6 | 23.57 | 28.05 | 29.9 | 112 | 95 | 7804 | 254 | 98.40 | 0.31 | 0.59 |
| | BB_1F | | | 1 | 31.79 | 32.4 | 24.78 | 28.75 | 30.16 | 53 | 55 | 3587 | 154 | 97.90 | 0.18 | 0.76 |
| | | | | 2 | 31.37 | 32.43 | 24.26 | 28.62 | 29.7 | 69 | 54 | 5010 | 169 | 98.34 | 0.18 | 0.56 |
| | | | | 3 | 30.7 | 32.02 | 24.32 | 28.39 | 30 | 107 | 72 | 4821 | 199 | 97.98 | 0.24 | 0.47 |
| | BB_1G | | | 1 | 33.08 | 33.01 | 26.11 | 29.46 | 29.84 | 23 | 37 | 1527 | 93 | 97.05 | 0.12 | 1.19 |
| | | | | 2 | 32.8 | 33.02 | 25.75 | 25.19 | 29.97 | 27 | 37 | 1924 | 1945 | 66.43 | 0.12 | 0.98 |
| | | | | 3 | NA | 33.26 | 26.12 | 29.16 | 29.24 | NA | 31 | 1517 | 115 | 96.35 | 0.10 | NA |
| | BB_1H | | | 1 | 33.37 | 34.82 | 28.06 | 30.04 | 29.44 | 19 | 11 | 436 | 61 | 93.42 | 0.04 | 0.43 |
| | | | | 2 | 34.25 | 34.98 | 27.86 | 30.53 | 29.81 | 11 | 10 | 496 | 43 | 95.82 | 0.03 | 0.70 |
| | | | | 3 | 34.25 | 33.63 | 27.85 | 29.75 | 29.62 | 11 | 24 | 499 | 76 | 92.97 | 0.08 | 1.73 |
| | BB_1I | | | 1 | NA | 33.91 | 29.97 | 30.09 | 29.15 | NA | 20 | 128 | 59 | 81.19 | 0.07 | NA |
| | | | | 2 | 34.07 | 34.4 | 27.42 | 28.77 | 29.24 | 12 | 14 | 658 | 152 | 89.66 | 0.05 | 0.91 |
| | | | | 3 | NA | 33.87 | 30.15 | 29.52 | 29.36 | NA | 21 | 114 | 89 | 71.93 | 0.07 | NA |
| Negative control | NC | NA | | 1 | NA | 33.4 | 0 | 30.57 | 29.81 | NA | 28 | 0 | 42 | NA | 0.09 | NA |
| Positive control | PC | | | 1 | 27.36 | 31.3 | 17.37 | 30.38 | 30.1 | 942 | 116 | 418691 | 48 | 99.99 | 0.38 | 0.08 |
| | | | | 2 | 27.23 | 31.2 | 17.73 | 30.44 | 29.92 | 1025 | 124 | 332252 | 46 | 99.99 | 0.41 | 0.08 |
| Human whole blood | BB_1A | 50 | 1 | 1 | 27.21 | 27.42 | 20.85 | 23.97 | NA | 1693 | 1540 | 208896 | 4882 | 98.84 | 5.06 | 1.00 |
| | | | | 2 | 27.23 | 27.46 | 21.53 | 24.12 | NA | 1675 | 1503 | 139575 | 4379 | 98.46 | 4.94 | 0.99 |
| | | | | 3 | 27.23 | 27.5 | 21.7 | 24.19 | NA | 1675 | 1467 | 126191 | 4162 | 98.38 | 4.82 | 0.97 |
| | | | | 4 | 27.16 | 27.35 | 21.38 | 24.19 | NA | 1739 | 1608 | 152559 | 4162 | 98.65 | 5.28 | 1.01 |
| | | | | 5 | 27.1 | 27.24 | 21.09 | 24.18 | NA | 1796 | 1719 | 181185 | 4193 | 98.86 | 5.65 | 1.05 |
| | | | | 6 | 27.01 | 27.34 | 21.4 | 24.23 | NA | 1885 | 1618 | 150761 | 4043 | 98.68 | 5.31 | 0.93 |
| | | | | 7 | 27.15 | 27.29 | 21.31 | 24.06 | NA | 1748 | 1668 | 159025 | 4574 | 98.58 | 5.48 | 1.05 |
| | | | | 8 | 27.07 | 27.32 | 21.08 | 24.34 | NA | 1825 | 1637 | 182263 | 3733 | 98.99 | 5.38 | 0.98 |
| | BB_1A_UV120 | | | 1 | 33.3 | 28.71 | 23.99 | 22.04 | NA | 65 | 701 | 32455 | 19792 | 76.63 | 2.30 | 18.75 |
| | | | | 2 | 31.19 | 27.99 | 24.03 | NA | NA | 200 | 1088 | 31695 | NA | NA | 3.57 | 8.03 |
| | | | | 3 | 32.41 | 28.66 | 24.32 | 21.98 | NA | 104 | 723 | 26687 | 20672 | 72.08 | 2.37 | 11.23 |
| | BB_1A_UV30 | | | 1 | 28.51 | 27.96 | 22.43 | 22.69 | NA | 843 | 1108 | 81853 | 12353 | 92.98 | 3.64 | 1.59 |
| | | | | 2 | 29.04 | 28.2 | 22.83 | 23.19 | NA | 634 | 957 | 64568 | 8596 | 93.76 | 3.14 | 1.90 |
| | | | | 3 | 29.14 | 28.19 | 22.72 | 23.3 | NA | 601 | 963 | 68921 | 7937 | 94.56 | 3.16 | 2.03 |
| | BB_1A_UV60 | | | 1 | 29.47 | 27.85 | 22.72 | 22.25 | NA | 504 | 1185 | 68921 | 16996 | 89.02 | 3.89 | 3.06 |
| | | | | 2 | 29.93 | 28.08 | 22.83 | 22.59 | NA | 393 | 1030 | 64568 | 13282 | 90.67 | 3.38 | 3.52 |
| | | | | 3 | 30.6 | 28.52 | 23.48 | 23.4 | NA | 275 | 787 | 43916 | 7382 | 92.25 | 2.58 | 4.05 |
| | BB_1B | | | 1 | 28.1 | 28.43 | NA | 24.72 | NA | 1050 | 832 | NA | 2834 | NA | 2.73 | 0.93 |
| | | | | 2 | 28.16 | 28.6 | 22.53 | 25.22 | NA | 1017 | 750 | 77140 | 1972 | 98.74 | 2.46 | 0.87 |
| | | | | 3 | 28.32 | 28.55 | 22.48 | 25.13 | NA | 933 | 773 | 79461 | 2105 | 98.69 | 2.54 | 0.99 |
| | BB_1C | | | 1 | 29.09 | 29.28 | 23.32 | 25.11 | NA | 617 | 495 | 48287 | 2136 | 97.84 | 1.63 | 1.01 |
| | | | | 2 | 29.22 | 29.41 | 23.49 | 25.57 | NA | 576 | 457 | 43657 | 1530 | 98.28 | 1.50 | 1.01 |
| | | | | 3 | 29.46 | 29.44 | 23.44 | 25.68 | NA | 506 | 449 | 44970 | 1413 | 98.45 | 1.47 | 1.15 |
| | BB_1D | | | 1 | 30.56 | 30.5 | 24.16 | 26.55 | NA | 281 | 235 | 29343 | 752 | 98.74 | 0.77 | 1.18 |
| | | | | 2 | 30.36 | 30.37 | 24.24 | 26.71 | NA | 312 | 254 | 27984 | 669 | 98.82 | 0.84 | 1.13 |
| | | | | 3 | 30.6 | 30.57 | 24.39 | 26.81 | NA | 275 | 225 | 25602 | 622 | 98.80 | 0.74 | 1.16 |
| | BB_1E | | | 1 | 31.9 | 31.35 | 25.27 | 27.32 | NA | 137 | 140 | 15193 | 430 | 98.60 | 0.46 | 1.59 |
| | | | | 2 | 31.84 | 31.36 | 25.49 | 26.89 | NA | 141 | 139 | 13335 | 587 | 97.84 | 0.46 | 1.53 |
| | | | | 3 | 31.24 | 31.43 | 25.56 | 27.16 | NA | 195 | 133 | 12793 | 483 | 98.15 | 0.44 | 1.01 |
| | BB_1F | | | 1 | NA | 33.43 | 31.14 | 29.61 | NA | NA | 39 | 468 | 82 | 91.97 | 0.13 | NA |
| | | | | 2 | 33.2 | 32.12 | 26.66 | 28.32 | NA | 68 | 87 | 6663 | 208 | 98.46 | 0.29 | 2.20 |
| | | | | 3 | 33.03 | 33.33 | 27.2 | 28.41 | NA | 75 | 42 | 4837 | 195 | 98.02 | 0.14 | 0.95 |
| | BB_1G | | | 1 | 33.95 | 32.54 | 27.96 | 28.19 | NA | 46 | 68 | 3082 | 229 | 96.42 | 0.22 | 2.69 |
| | | | | 2 | 33.93 | 32.79 | 27.71 | 28.15 | NA | 46 | 58 | 3575 | 236 | 96.81 | 0.19 | 2.28 |
| | | | | 3 | NA | 32.96 | 28.72 | 28.87 | NA | NA | 52 | 1964 | 140 | 96.56 | 0.17 | NA |
| | BB_1H | | | 1 | NA | 34 | 29.76 | 28.64 | NA | NA | 28 | 1060 | 165 | 92.78 | 0.09 | NA |
| | | | | 2 | NA | 33.63 | 30.18 | 29.08 | NA | NA | 35 | 826 | 120 | 93.23 | 0.11 | NA |
| | | | | 3 | NA | 32.96 | 29.6 | 29.08 | NA | NA | 52 | 1166 | 120 | 95.10 | 0.17 | NA |
| | BB_1I | | | 1 | NA | NA | 32.3 | 28.9 | NA | NA | NA | 235 | 137 | 77.47 | NA | NA |
| | | | | 2 | NA | 33.39 | 32.02 | 28.93 | NA | NA | 40 | 278 | 134 | 80.58 | 0.13 | NA |
| | | | | 3 | NA | 34.51 | 31.93 | 27.42 | NA | NA | 20 | 293 | 400 | 59.41 | 0.07 | NA |
| | BB_2A | 5 | | 1 | 31.16 | 31.73 | 25.58 | 26.3 | NA | 203 | 111 | 12642 | 901 | 96.56 | 0.36 | 0.80 |
| | | | | 2 | 31.08 | 31.76 | 25.54 | 26.18 | NA | 212 | 109 | 12945 | 983 | 96.34 | 0.36 | 0.75 |
| | | | | 3 | 31.53 | 31.46 | 25.47 | 26.23 | NA | 167 | 131 | 13494 | 948 | 96.61 | 0.43 | 1.19 |
| | | | | 4 | 31.44 | 31.09 | 25.81 | 26.2 | NA | 175 | 164 | 11030 | 969 | 95.79 | 0.54 | 1.41 |
| | | | | 5 | 31.3 | 31.55 | 25.62 | 26.27 | NA | 189 | 124 | 12346 | 921 | 96.40 | 0.41 | 0.98 |
| Negative control | NC | NA | | 1 | NA | 33.87 | NA | 29.25 | NA | NA | 30 | NA | 106 | NA | 0.10 | NA |
| | | | | 2 | NA | 32.76 | NA | 29.46 | NA | NA | 59 | NA | 91 | NA | 0.19 | NA |
| | | | | 3 | NA | 34.23 | NA | 29.48 | NA | NA | 24 | NA | 90 | NA | 0.08 | NA |

| | | | | | | | | | | | | | | | | | |
|---|---|---|---|---|---|---|---|---|---|---|---|---|---|---|---|---|---|
| **CFX384** | *Gallus gallus domesticus* | NHS_1 | 200 | | 1 | NA | 33.15 | NA | 28.85 | NA | NA | 47 | NA | 142 | NA | 0.15 | NA |
| | | | | | 2 | NA | 34.14 | NA | 28.35 | NA | NA | 25 | NA | 204 | NA | 0.08 | NA |
| | *Sus scrofa domesticus* | NHS_2 | 80 | | 1 | NA | NA | NA | 28.62 | NA | NA | 0 | NA | 168 | NA | NA | NA |
| | | | | | 2 | NA | 34.06 | NA | 28.34 | NA | NA | 27 | NA | 205 | NA | 0.09 | NA |
| | *Felis catus* | NHS_3 | | | 1 | NA | NA | NA | 29.04 | NA | NA | 0 | NA | 124 | NA | NA | NA |
| | | | | | 2 | NA | 34.31 | NA | 29.29 | NA | NA | 23 | NA | 103 | NA | 0.08 | NA |
| | *Canis lupus familiaris* | NHS_4 | | | 1 | NA | 32.96 | NA | 29.28 | NA | NA | 52 | NA | 104 | NA | 0.17 | NA |
| | | | | | 2 | NA | 33.75 | NA | 28.86 | NA | NA | 32 | NA | 141 | NA | 0.11 | NA |
| | *Rattus* | NHS_5 | | | 1 | NA | 34.99 | NA | 28.62 | NA | NA | 15 | NA | 168 | NA | 0.05 | NA |
| | | | | | 2 | NA | 34.19 | NA | 28.89 | NA | NA | 25 | NA | 138 | NA | 0.08 | NA |
| | *Mus musculus* | NHS_6 | | | 1 | NA | 34.25 | NA | 29.37 | NA | NA | 24 | NA | 97 | NA | 0.08 | NA |
| | | | | | 2 | NA | 33.59 | NA | 28.89 | NA | NA | 36 | NA | 138 | NA | 0.12 | NA |
| | *Bos taurus* | NHS_7 | | | 1 | NA | 34.51 | NA | 28.48 | NA | NA | 20 | NA | 185 | NA | 0.07 | NA |
| | | | | | 2 | NA | 33.7 | NA | 28.73 | NA | NA | 33 | NA | 155 | NA | 0.11 | NA |
| | *Rhesus* | NHS_8 | | | 1 | 34.24 | 29.3 | 28.78 | 28.14 | NA | 39 | 489 | 1895 | 237 | 94.11 | 1.61 | 23.22 |
| | | | | | 2 | NA | 29.31 | NA | NA | NA | NA | 486 | NA | NA | NA | 1.60 | NA |
| | Positive control | PC | NA | | 1 | 28.31 | 31.23 | 19.61 | 29.27 | NA | 938 | 151 | 435779 | 105 | 99.99 | 0.49 | 0.19 |
| | | | | | 2 | 27.76 | 31.59 | 19.19 | 29.01 | NA | 1260 | 121 | 559022 | 126 | 99.99 | 0.40 | 0.11 |
| | | | | | 3 | 28.15 | 31.07 | 19.52 | 29.23 | NA | 1022 | 166 | 459668 | 108 | 99.99 | 0.55 | 0.19 |
| | Human whole blood | BB_1A | 50 | 2 | 1 | 27.16 | 27.3 | 21.23 | 22.83 | NA | 1739 | 1658 | 166751 | 11160 | 96.76 | 5.44 | 1.05 |
| | | | | | 2 | 27.42 | 27.64 | 20.37 | 23.95 | NA | 1513 | 1347 | 277679 | 4954 | 99.12 | 4.42 | 1.00 |
| | | | | | 3 | 27.27 | 27.53 | 21.36 | 23.91 | NA | 1639 | 1440 | 154379 | 5099 | 98.38 | 4.73 | 0.97 |
| | | BB_1B | | | 1 | 28.31 | 28.8 | 22.18 | 24.5 | NA | 938 | 663 | 94932 | 3324 | 98.28 | 2.18 | 0.84 |
| | | | | | 2 | 28.17 | 28.56 | 22.43 | 24.01 | NA | 1012 | 768 | 81853 | 4743 | 97.18 | 2.52 | 0.90 |
| | | | | | 3 | 28.26 | 28.63 | 22.05 | 24.29 | NA | 964 | 736 | 102540 | 3871 | 98.15 | 2.42 | 0.91 |
| | | BB_1C | | | 1 | 29.08 | 29.23 | 23.07 | 24.96 | NA | 621 | 510 | 56003 | 2381 | 97.92 | 1.68 | 1.04 |
| | | | | | 2 | 29.37 | 29.54 | 22.58 | 25.36 | NA | 531 | 422 | 74886 | 1782 | 98.82 | 1.39 | 1.03 |
| | | | | | 3 | 29.4 | 29.47 | 23.21 | 25.54 | NA | 523 | 441 | 51542 | 1564 | 98.51 | 1.45 | 1.09 |
| | | BB_1D | | | 1 | 30.14 | 30.41 | 24 | 26.11 | NA | 352 | 248 | 32263 | 1034 | 98.42 | 0.82 | 0.97 |
| | | | | | 2 | 30.25 | 30.33 | 24.05 | 26.65 | NA | 331 | 261 | 31321 | 699 | 98.90 | 0.86 | 1.08 |
| | | | | | 3 | 30.38 | 30.77 | 24.47 | 26.94 | NA | 309 | 199 | 24416 | 566 | 98.85 | 0.65 | 0.90 |
| | | BB_1E | | | 1 | 31.14 | 31.31 | 25.21 | 26.47 | NA | 206 | 143 | 15743 | 797 | 97.53 | 0.47 | 1.03 |
| | | | | | 2 | 31.5 | 31.1 | 25.06 | 27.31 | NA | 169 | 163 | 17208 | 433 | 98.76 | 0.54 | 1.45 |
| | | | | | 3 | 31.19 | 31.75 | NA | 27.16 | NA | 200 | 110 | NA | 483 | NA | 0.36 | 0.81 |
| | | BB_1F | | | 1 | 33.43 | 32.37 | 26.14 | 27.15 | NA | 60 | 75 | 9070 | 486 | 97.39 | 0.25 | 2.17 |
| | | | | | 2 | 32.83 | 32.38 | 26.24 | 27.5 | NA | 83 | 75 | 8548 | 377 | 97.84 | 0.25 | 1.50 |
| | | | | | 3 | 33.8 | 33.82 | 26.26 | 27.9 | NA | 49 | 31 | 8447 | 282 | 98.36 | 0.10 | 1.12 |
| | | BB_1G | | | 1 | 34.2 | 34.1 | 27.19 | 27.84 | NA | 40 | 26 | 4866 | 295 | 97.06 | 0.09 | 1.21 |
| | | | | | 2 | 34.75 | 33.18 | 27.32 | 28.44 | NA | 30 | 46 | 4505 | 191 | 97.93 | 0.15 | 2.97 |
| | | | | | 3 | 33.83 | 33 | 27.75 | 28.13 | NA | 49 | 51 | 3491 | 239 | 96.69 | 0.17 | 1.89 |
| | | BB_1H | | | 1 | NA | 33.6 | 28.57 | 28.69 | NA | NA | 35 | 2147 | 159 | 96.42 | 0.12 | NA |
| | | | | | 2 | 34.05 | NA | 29.19 | 28.82 | NA | 43 | NA | 1486 | 145 | 95.35 | NA | NA |
| | | | | | 3 | 34.1 | 34.21 | 28.57 | 28.56 | NA | 42 | 24 | 2147 | 175 | 96.08 | 0.08 | 1.06 |
| | | BB_1I | | | 1 | NA | 34.47 | 31.49 | 28.96 | NA | NA | 21 | 380 | 131 | 85.31 | 0.07 | NA |
| | | | | | 2 | NA | 33.04 | 31.05 | 28.54 | NA | NA | 50 | 493 | 178 | 84.75 | 0.16 | NA |
| | | | | | 3 | NA | 34.08 | 31.27 | 28.59 | NA | NA | 26 | 433 | 171 | 83.49 | 0.09 | NA |
| | Negative control | NC | NA | | 1 | NA | NA | NA | 29.41 | NA | NA | NA | NA | 94 | NA | NA | NA |
| | | | | | 2 | NA | 33.42 | NA | 29.16 | NA | NA | 40 | NA | 113 | NA | 0.13 | NA |
| | | | | | 3 | NA | 33.94 | NA | 29.16 | NA | NA | 29 | NA | 113 | NA | 0.09 | NA |
| | Positive control | PC | | | 1 | 28.02 | 31.31 | 19.3 | 28.96 | NA | 1096 | 143 | 523722 | 131 | 99.99 | 0.47 | 0.15 |
| | | | | | 2 | 28.08 | 32 | 19.33 | 29.39 | NA | 1062 | 94 | 514487 | 96 | 99.99 | 0.31 | 0.10 |
| | | | | | 3 | 27.79 | 31.09 | 19 | 29.18 | NA | 1240 | 164 | 625690 | 112 | 99.99 | 0.54 | 0.15 |
| | Human whole blood | BB_1A | 50 | | 1 | 26.32 | 26.39 | 19.11 | 24.64 | 26.63 | 2787 | 2347 | 190455 | 3247 | 99.15 | 7.71 | 1.08 |
| | | | | | 2 | 26.4 | 26.47 | 19.01 | 25.17 | 26.61 | 2663 | 2212 | 202122 | 2279 | 99.44 | 7.27 | 1.08 |
| | | | | | 3 | 26.63 | 26.84 | 19.16 | 25.34 | 27.58 | 2338 | 1684 | 184876 | 2035 | 99.45 | 5.53 | 0.98 |
| | | | | | 4 | 26.86 | 26.64 | 18.46 | 24.38 | 27.47 | 2052 | 1952 | 280310 | 3862 | 99.32 | 6.41 | 1.34 |
| | | | | | 5 | 26.8 | 26.81 | 18.09 | 24.64 | 27.6 | 2123 | 1722 | 349287 | 3247 | 99.54 | 5.65 | 1.13 |
| | | | | | 6 | 26.45 | 26.67 | 18.41 | 24.45 | 27.24 | 2589 | 1909 | 288769 | 3686 | 99.37 | 6.27 | 0.97 |
| | | | | | 7 | 26.62 | 26.65 | 18.13 | 24.94 | 27.56 | 2351 | 1938 | 341078 | 2657 | 99.61 | 6.36 | 1.11 |
| | | | | | 8 | 26.15 | 26.25 | 18.01 | 24.95 | 27.09 | 3069 | 2602 | 366303 | 2640 | 99.64 | 8.54 | 1.06 |
| | | BB_1A_UV120 | | | 1 | NA | 27.74 | 20.92 | 21.8 | 27.11 | NA | 868 | 64923 | 21622 | 85.72 | 2.85 | NA |
| | | | | | 2 | 33.81 | 27.57 | 21.58 | 22.1 | 26.56 | 40 | 983 | 43850 | 17698 | 83.21 | 3.23 | 113.19 |
| | | | | | 3 | 12.49 | 28.6 | 22.27 | 22.5 | 27.08 | 7076403 | 460 | 29093 | 13550 | 81.11 | 1.51 | 0.00 |
| | | BB_1A_UV30 | | | 1 | 28.26 | 27.01 | 19.49 | 23.23 | 26.23 | 928 | 1486 | 151938 | 8323 | 97.33 | 4.88 | 2.86 |
| | | | | | 2 | 27.63 | 27.17 | 19.53 | 23.54 | 27.07 | 1326 | 1321 | 148367 | 6767 | 97.77 | 4.34 | 1.60 |
| | | | | | 3 | 29.1 | 26.86 | 19.42 | 24.95 | 27.02 | 576 | 1660 | 158395 | 2640 | 99.17 | 5.45 | 5.93 |
| | | BB_1A_UV60 | | | 1 | 29.97 | 27.19 | 19.89 | 23.41 | 27.13 | 352 | 1301 | 119777 | 7380 | 97.01 | 4.27 | 8.83 |
| | | | | | 2 | 29.91 | 27.44 | 20.03 | 23.99 | 26.47 | 364 | 1082 | 110210 | 5011 | 97.78 | 3.55 | 7.03 |
| | | | | | 3 | NA | 28.76 | 24.46 | 25.56 | 26.49 | NA | 409 | 7912 | 1757 | 90.01 | 1.34 | NA |
| | | BB_1B | | | 1 | 27.43 | 27.66 | 19.09 | 25.42 | 27.48 | 1486 | 920 | 192733 | 1929 | 99.50 | 3.02 | 0.96 |
| | | | | | 2 | 27.72 | 27.98 | 19.08 | 25.2 | 27.6 | 1260 | 727 | 193882 | 2234 | 99.43 | 2.39 | 0.94 |
| | | | | | 3 | 27.81 | 28.07 | 19.75 | 25.69 | 27.83 | 1198 | 680 | 130175 | 1611 | 99.39 | 2.23 | 0.94 |
| | | BB_1C | | | 1 | 28.4 | 28.24 | 20.22 | 26.25 | 27.09 | 857 | 600 | 98437 | 1108 | 99.44 | 1.97 | 1.28 |
| | | | | | 2 | 29.02 | 28.8 | 20.76 | 25.97 | 27.97 | 603 | 397 | 71403 | 1336 | 99.07 | 1.30 | 1.34 |
| | | | | | 3 | 28.76 | 28.73 | 20.56 | 25.86 | 27.75 | 699 | 418 | 80420 | 1438 | 99.11 | 1.37 | 1.16 |
| | | BB_1D | | | 1 | 30.2 | 29.83 | 21.14 | 25.46 | 27.62 | 309 | 186 | 56963 | 1878 | 98.38 | 0.61 | 1.50 |
| | | | | | 2 | 30.01 | 29.6 | 21.56 | 26.09 | 27.85 | 344 | 220 | 44375 | 1233 | 98.63 | 0.72 | 1.54 |

| | | | | | | | | | | | | | | | | | |
|---|---|---|---|---|---|---|---|---|---|---|---|---|---|---|---|---|---|
| QS5 | | | | 1 | 3 | 30.05 | 29.56 | 21.41 | 25.98 | 27.97 | 336 | 227 | 48514 | 1327 | 98.65 | 0.74 | 1.63 |
| | | BB_1E | | | 1 | 31.57 | 30.8 | 22.69 | 27.72 | 27.64 | 142 | 91 | 22664 | 415 | 99.09 | 0.30 | 2.01 |
| | | | | | 2 | 31.56 | 30.63 | 23.06 | 27.87 | 27.98 | 143 | 103 | 18188 | 376 | 98.98 | 0.34 | 2.26 |
| | | | | | 3 | 33.29 | 30.94 | 23.17 | 28.18 | 27.85 | 54 | 82 | 17037 | 305 | 99.11 | 0.27 | 6.44 |
| | | BB_1F | | | 1 | NA | 31.79 | 24.06 | 29.27 | 27.85 | NA | 44 | 10036 | 148 | 99.27 | 0.14 | NA |
| | | | | | 2 | 32.29 | 31.67 | 24.06 | 29.08 | 27.48 | 95 | 48 | 10036 | 168 | 99.17 | 0.16 | 1.80 |
| | | | | | 3 | 32.72 | 31.97 | 24.4 | 28.95 | 27.86 | 74 | 38 | 8199 | 183 | 98.90 | 0.13 | 1.98 |
| | | BB_1G | | | 1 | 33.73 | 31.5 | 25.02 | 29.57 | 27.75 | 42 | 54 | 5671 | 121 | 98.95 | 0.18 | 5.89 |
| | | | | | 2 | NA | 31.37 | 26.08 | 30.13 | 28.08 | NA | 60 | 3020 | 83 | 98.64 | 0.20 | NA |
| | | | | | 3 | NA | 31.82 | 25.75 | 29.68 | 27.57 | NA | 43 | 3674 | 112 | 98.50 | 0.14 | NA |
| | | BB_1H | | | 1 | NA | 31.85 | 27.71 | 30.31 | 26.94 | NA | 42 | 1146 | 74 | 96.88 | 0.14 | NA |
| | | | | | 2 | NA | 32.53 | 28.18 | 29 | 27.42 | NA | 25 | 866 | 177 | 90.75 | 0.08 | NA |
| | | | | | 3 | NA | 31.94 | 28.35 | 30.4 | 27.36 | NA | 39 | 783 | 69 | 95.76 | 0.13 | NA |
| | | BB_1I | | | 1 | NA | 32.16 | 29.72 | 29.66 | 27.82 | NA | 33 | 347 | 114 | 85.91 | 0.11 | NA |
| | | | | | 2 | NA | 32.44 | 28.99 | 29.01 | 27.63 | NA | 27 | 535 | 176 | 85.91 | 0.09 | NA |
| | | | | | 3 | NA | 32.08 | 29.09 | 29.51 | 27.41 | NA | 35 | 504 | 126 | 88.92 | 0.12 | NA |
| | | BB_2A | 5 | | 1 | 31.73 | 30.89 | 23.01 | 28.07 | 27.48 | 130 | 85 | 18737 | 329 | 99.13 | 0.28 | 2.11 |
| | | | | | 2 | 32.16 | 30.71 | 23.3 | 27.96 | 27.78 | 102 | 97 | 15770 | 354 | 98.89 | 0.32 | 3.31 |
| | | | | | 3 | 31 | 30.52 | 21.8 | 28.02 | 26.83 | 196 | 112 | 38473 | 340 | 99.56 | 0.37 | 1.62 |
| | | | | | 4 | 32.42 | 29.92 | 22.51 | 27.67 | 27.16 | 88 | 174 | 25224 | 429 | 99.16 | 0.57 | 7.19 |
| | | | | | 5 | 31 | 30.47 | 23.34 | 27.95 | 27.49 | 196 | 116 | 15399 | 356 | 98.86 | 0.38 | 1.68 |
| | Negative control | NC | NA | | 1 | NA | 33.5 | NA | 31.24 | 24.51 | NA | 12 | NA | 40 | NA | 0.04 | NA |
| | | | | | 2 | NA | 32.09 | NA | 32.01 | 27.26 | NA | 35 | NA | 24 | NA | 0.12 | NA |
| | | | | | 3 | 28.08 | 32.35 | 34.5 | 31.86 | 27.43 | 1028 | 29 | 20 | 26 | 60.70 | 0.10 | 0.05 |
| | *Gallus gallus domesticus* | NHS_1 | 200 | | 1 | NA | 33.96 | NA | 29.61 | 26.72 | NA | 9 | NA | 118 | NA | 0.03 | NA |
| | | | | | 2 | NA | 32.73 | NA | 29.93 | 26.85 | NA | 22 | NA | 95 | NA | 0.07 | NA |
| | *Sus scrofa domesticus* | NHS_2 | 80 | | 1 | NA | 33.23 | NA | 29.28 | 27.71 | NA | 15 | NA | 147 | NA | 0.05 | NA |
| | | | | | 2 | NA | 34.25 | NA | 28.95 | 27.46 | NA | 7 | NA | 183 | NA | 0.02 | NA |
| | *Felis catus* | NHS_3 | | | 1 | NA | 31.91 | NA | 31.52 | 27.16 | NA | 40 | NA | 33 | NA | 0.13 | NA |
| | | | | | 2 | NA | 32.41 | NA | 29.02 | 27.78 | NA | 28 | NA | 174 | NA | 0.09 | NA |
| | *Canis lupus familiaris* | NHS_4 | | | 1 | NA | 32.62 | NA | 30.81 | 27.5 | NA | 24 | NA | 53 | NA | 0.08 | NA |
| | | | | | 2 | 30.97 | 33.03 | NA | 30.1 | 27.83 | 200 | 18 | NA | 85 | NA | 0.06 | 0.25 |
| | *Rattus* | NHS_5 | | | 1 | NA | 32.11 | NA | 29.42 | 27.72 | NA | 35 | NA | 133 | NA | 0.11 | NA |
| | | | | | 2 | NA | 32.9 | NA | 29.77 | 27.73 | NA | 19 | NA | 106 | NA | 0.06 | NA |
| | *Mus musculus* | NHS_6 | | | 1 | 26.6 | 32.65 | NA | 30.27 | 26.57 | 2378 | 23 | NA | 76 | NA | 0.08 | 0.01 |
| | | | | | 2 | NA | 33.45 | NA | 30.29 | 27.67 | NA | 13 | NA | 75 | NA | 0.04 | NA |
| | *Bos taurus* | NHS_7 | | | 1 | NA | 31.91 | 34.7 | 29.94 | 27.48 | NA | 40 | 18 | 94 | 27.56 | 0.13 | NA |
| | | | | | 2 | NA | 31.56 | 34.8 | 29.75 | 27.2 | NA | 52 | 17 | 107 | 24.00 | 0.17 | NA |
| | *Rhesus* | NHS_8 | | | 1 | NA | 30.37 | 22.99 | 28.59 | 27.58 | NA | 125 | 18961 | 232 | 99.39 | 0.41 | NA |
| | | | | | 2 | NA | 30.23 | 23.07 | 28.18 | 27.59 | NA | 138 | 18081 | 305 | 99.16 | 0.45 | NA |
| | Positive control | PC | NA | | 1 | 27.18 | 30.7 | 17.41 | 30.82 | 27.39 | 1712 | 98 | 523331 | 52 | 99.99 | 0.32 | 0.09 |
| | | | | | 2 | 26.93 | 29.9 | 17.02 | 30.52 | 26.9 | 1972 | 177 | 659911 | 64 | 100.00 | 0.58 | 0.13 |
| | | | | | 3 | 26.93 | 29.33 | 16.26 | 31.55 | 26.65 | 1972 | 269 | 1036901 | 32 | 100.00 | 0.88 | 0.19 |
| | | BB_1A | 50 | 2 | 1 | 26.07 | 26.5 | 26.82 | 28.01 | 21.07 | 2881 | 1833 | 156683 | 8 | 100.00 | 6.02 | 0.84 |
| | | | | | 2 | 26.18 | 26.82 | 26.82 | 27.69 | 21.2 | 2700 | 1466 | 156683 | 2755 | 99.13 | 4.81 | 0.73 |
| | | | | | 3 | 25.33 | 26.86 | 27.04 | 27.74 | 20.99 | 4460 | 1425 | 135842 | 1109 | 99.59 | 4.68 | 0.39 |
| | | BB_1B | | | 1 | 27.29 | 27.65 | 27.93 | 27.96 | 21.57 | 1402 | 820 | 76255 | 20 | 99.99 | 2.69 | 0.88 |
| | | | | | 2 | 27.16 | 27.92 | 27.63 | 27.99 | 21.68 | 1514 | 679 | 92639 | 12 | 99.99 | 2.23 | 0.67 |
| | | | | | 3 | 27.42 | 27.74 | 27.88 | 27.32 | 21.82 | 1298 | 770 | 78769 | 2322138 | 6.35 | 2.53 | 0.91 |
| | | BB_1C | | | 1 | 28.61 | 28.79 | 28.53 | 28.22 | 22.66 | 643 | 369 | 51667 | 0 | 100.00 | 1.21 | 1.00 |
| | | | | | 2 | 28.09 | 28.87 | 28.55 | 28.4 | 22.55 | 874 | 349 | 51001 | 0 | 100.00 | 1.15 | 0.66 |
| | | | | | 3 | 28.56 | 28.7 | 28.49 | 28.08 | 22.38 | 662 | 393 | 53025 | 2 | 100.00 | 1.29 | 1.03 |
| | | BB_1D | | | 1 | 29.38 | 29.64 | 28.12 | 28.12 | 23.23 | 408 | 204 | 67411 | 1 | 100.00 | 0.67 | 0.95 |
| | | | | | 2 | 30.08 | 29.65 | 28.56 | 26.99 | 23.5 | 270 | 202 | 50671 | 944723557 | 0.01 | 0.66 | 1.54 |
| | | | | | 3 | 29.35 | 30.05 | 28.2 | 28.24 | 23.11 | 415 | 153 | 64002 | 0 | 100.00 | 0.50 | 0.70 |
| | | BB_1E | | | 1 | 31.09 | 30.63 | 29.71 | NA | 24.6 | 149 | 102 | 24029 | NA | NA | 0.33 | 1.57 |
| | | | | | 2 | 31.38 | 30.24 | 30.25 | 27.43 | 24.88 | 125 | 134 | 16927 | 313388 | 9.75 | 0.44 | 2.53 |
| | | | | | 3 | 31.12 | 30 | 29.87 | NA | 24.9 | 146 | 158 | 21660 | NA | NA | 0.52 | 2.49 |
| | | BB_1F | | | 1 | 33.57 | 31.2 | 30.79 | 28.11 | 25.93 | 34 | 68 | 11924 | 1 | 99.99 | 0.22 | 5.98 |
| | | | | | 2 | 32.4 | 32.46 | 31.11 | 27.94 | 26.37 | 69 | 28 | 9689 | 29 | 99.85 | 0.09 | 1.09 |
| | | | | | 3 | 33.31 | 32.01 | 31.03 | 27.76 | 26.15 | 40 | 39 | 10205 | 770 | 96.36 | 0.13 | 2.83 |
| | | BB_1G | | | 1 | 33.69 | 32.25 | 32.16 | 27.87 | 27.52 | 32 | 33 | 4903 | 104 | 98.95 | 0.11 | 3.12 |
| | | | | | 2 | NA | 31.97 | 32.56 | 28.24 | 28.56 | NA | 40 | 3782 | 0 | 100.00 | 0.13 | NA |
| | | | | | 3 | NA | 31.86 | 31.99 | 28.17 | 27.92 | NA | 43 | 5474 | 0 | 100.00 | 0.14 | NA |
| | | BB_1H | | | 1 | NA | 32.36 | 31.84 | 27.94 | 30.03 | NA | 30 | 6034 | 29 | 99.76 | 0.10 | NA |
| | | | | | 2 | NA | 33.46 | 32.5 | 27.98 | 30.94 | NA | 14 | 3932 | 14 | 99.82 | 0.05 | NA |
| | | | | | 3 | NA | 32.49 | 32.71 | 28.15 | 30.2 | NA | 28 | 3431 | 1 | 99.99 | 0.09 | NA |
| | | BB_1I | | | 1 | NA | 33.93 | 32.39 | 28.44 | 31.88 | NA | 10 | 4223 | 0 | 100.00 | 0.03 | NA |
| | | | | | 2 | NA | NA | 32.3 | 28.06 | 31.87 | NA | NA | 4477 | 3 | 99.96 | NA | NA |
| | | | | | 3 | NA | 33.81 | 32.28 | 28.07 | 31.25 | NA | 11 | 4535 | 3 | 99.97 | 0.04 | NA |
| | Negative control | NC | NA | | 1 | NA | 33.77 | 33.35 | 28.43 | NA | NA | 11 | 2265 | 0 | 100.00 | 0.04 | NA |
| | Positive control | PC | | | 1 | 27.06 | 30.33 | 34.72 | 27.83 | 18.64 | 1606 | 126 | 931 | 215 | 89.64 | 0.41 | 0.12 |
| | | | | | 2 | 26.8 | 30.54 | 34.46 | 27.47 | 18.02 | 1872 | 109 | 1102 | 151283 | 1.44 | 0.36 | 0.08 |
| | | BB_1A | | | 1 | 27.97 | 28.15 | 24.54 | 27.54 | 27.15 | 1174 | 1189 | 121438 | 2389 | 99.03 | 3.90 | 1.00 |
| | | | | | 2 | 27.79 | 28.23 | 24.62 | 27.77 | 27.19 | 1309 | 1123 | 115928 | 2042 | 99.13 | 3.69 | 0.83 |
| | | | | | 3 | 27.71 | 28.02 | 23.86 | 27.33 | 27.1 | 1374 | 1304 | 180208 | 2758 | 99.24 | 4.28 | 0.91 |
| | | | | | 4 | 27.66 | 28.03 | 24.07 | 27.4 | 27.18 | 1417 | 1295 | 159527 | 2629 | 99.18 | 4.25 | 0.88 |

| | | | | | | | | | | | | | | | | | |
|---|---|---|---|---|---|---|---|---|---|---|---|---|---|---|---|---|---|
| | | BB_1A | | | 5 | 27.35 | 28.08 | 23.98 | 27.53 | 27.04 | 1709 | 1249 | 168083 | 2406 | 99.29 | 4.10 | 0.68 |
| | | | | | 6 | 26.99 | 27.96 | 24.26 | 27.68 | 26.59 | 2124 | 1361 | 142869 | 2171 | 99.25 | 4.47 | 0.57 |
| | | | | | 7 | 27.89 | 27.87 | 24.04 | 27.4 | 26.83 | 1233 | 1451 | 162330 | 2629 | 99.20 | 4.76 | 1.15 |
| | | | | | 8 | 17.66 | 17.56 | 23.18 | 27.12 | 17.18 | 599260 | 2212901 | 267419 | 3184 | 99.41 | NA | NA |
| | | BB_1A_UV120 | | | 1 | NA | 29.25 | 25.2 | 25.48 | 26.94 | NA | 544 | 82790 | 9769 | 94.43 | 1.79 | NA |
| | | | | | 2 | NA | 29.05 | 25.25 | 26.4 | 27.07 | NA | 627 | 80421 | 5209 | 96.86 | 2.06 | NA |
| | | | | | 3 | NA | 29.57 | 25.32 | 26.21 | 27.26 | NA | 433 | 77219 | 5931 | 96.30 | 1.42 | NA |
| | | BB_1A_UV30 | | | 1 | 31.33 | 28.36 | 26.49 | 25.98 | 26.79 | 154 | 1024 | 39155 | 6941 | 91.86 | 3.36 | 9.40 |
| | | | | | 2 | 31.1 | 28.83 | 26.63 | 27.13 | 27.22 | 177 | 733 | 36099 | 3162 | 95.80 | 2.41 | 5.72 |
| | | | | | 3 | NA | 28.72 | 24.74 | 26.43 | 27.17 | NA | 793 | 108128 | 5103 | 97.69 | 2.60 | NA |
| | | BB_1A_UV60 | | | 1 | NA | 28.96 | 29.28 | 25.15 | 27.2 | NA | 668 | 7753 | 12241 | 55.88 | 2.19 | NA |
| | | | | | 2 | NA | 29.35 | 28.27 | 25 | 27.2 | NA | 507 | 13934 | 13563 | 67.26 | 1.66 | NA |
| | | | | | 3 | 21.23 | 21.14 | 28.63 | 25.36 | 20.74 | 69183 | 173610 | 11306 | 10604 | 68.08 | 570.09 | 1.21 |
| | | BB_1B | 50 | 1 | 1 | 29.1 | 29.12 | 25.28 | 28.38 | 27.31 | 593 | 597 | 79033 | 1346 | 99.16 | 1.96 | 1.12 |
| | | | | | 2 | 28.89 | 29.31 | 25.49 | 28.34 | 27.36 | 673 | 521 | 69963 | 1383 | 99.02 | 1.71 | 0.84 |
| | | | | | 3 | 29.13 | 29.74 | 25.69 | 28.66 | 27.24 | 582 | 384 | 62295 | 1111 | 99.12 | 1.26 | 0.74 |
| | | BB_1C | | | 1 | 29.38 | 29.82 | 27.58 | 28.23 | 27.39 | 501 | 363 | 20797 | 1491 | 96.54 | 1.19 | 0.83 |
| | | | | | 2 | NA | 29.91 | 26.53 | 29.03 | 27.24 | NA | 340 | 38256 | 863 | 98.88 | 1.12 | NA |
| | | | | | 3 | 30.3 | 30.13 | 26.67 | 29.02 | 27.1 | 287 | 291 | 35270 | 869 | 98.78 | 0.96 | 1.28 |
| | | BB_1D | | | 1 | NA | NA | NA | NA | 27.17 | NA | NA | NA | NA | NA | NA | NA |
| | | | | | 2 | 33.13 | 31.06 | 27.89 | 29.01 | 27.52 | 52 | 150 | 17372 | 875 | 97.54 | 0.49 | 4.96 |
| | | | | | 3 | 31.69 | 31.18 | 27.94 | 29 | 27.15 | 124 | 138 | 16876 | 881 | 97.46 | 0.45 | 1.64 |
| | | BB_1E | | | 1 | NA | 32.23 | 29.66 | 30.87 | 27.55 | NA | 65 | 6218 | 245 | 98.07 | 0.21 | NA |
| | | | | | 2 | NA | 31.94 | 29.41 | 30.77 | 27.6 | NA | 80 | 7189 | 263 | 98.21 | 0.26 | NA |
| | | | | | 3 | NA | 32 | 29.68 | 31.01 | 27.5 | NA | 77 | 6146 | 223 | 98.22 | 0.25 | NA |
| | | BB_1F | | | 1 | NA | 32.53 | NA | 30.17 | 27.29 | NA | 53 | NA | 396 | NA | 0.17 | NA |
| | | | | | 2 | NA | 33.43 | 31.67 | 32.01 | 27.58 | NA | 28 | 1936 | 113 | 97.18 | 0.09 | NA |
| | | | | | 3 | NA | 33.11 | 32.88 | 31.79 | 27.39 | NA | 35 | 959 | 131 | 93.62 | 0.11 | NA |
| | | BB_1G | | | 1 | NA | 32.93 | NA | 33.51 | 27.47 | NA | 40 | NA | 40 | NA | 0.13 | NA |
| | | | | | 2 | NA | 34.5 | NA | 33.83 | 27.78 | NA | 13 | NA | 32 | NA | 0.04 | NA |
| | | | | | 3 | NA | 33.94 | NA | 33.19 | 26.98 | NA | 19 | NA | 50 | NA | 0.06 | NA |
| | | BB_1H | | | 1 | NA | 33.79 | NA | 33.69 | 27.6 | NA | 22 | NA | 36 | NA | 0.07 | NA |
| | | | | | 2 | NA | 33.09 | NA | 32.99 | 27.5 | NA | 35 | NA | 58 | NA | 0.12 | NA |
| | | | | | 3 | NA | 33.32 | NA | 33.38 | 27.47 | NA | 30 | NA | 44 | NA | 0.10 | NA |
| | | BB_1I | | | 1 | NA | 33.01 | NA | 31.58 | 27.35 | NA | 38 | NA | 151 | NA | 0.12 | NA |
| | | | | | 2 | NA | 33.91 | NA | 32.17 | 27.54 | NA | 20 | NA | 101 | NA | 0.07 | NA |
| | | | | | 3 | NA | 34.31 | NA | 32.04 | 27.28 | NA | 15 | NA | 110 | NA | 0.05 | NA |
| | | BB_2A | 5 | | 1 | NA | 33.79 | 29.37 | 29.51 | 26.97 | NA | 22 | 7358 | 622 | 95.95 | 0.07 | NA |
| | | | | | 2 | NA | 30.74 | 29.55 | 29.26 | 27.42 | NA | 189 | 6628 | 737 | 94.73 | 0.62 | NA |
| | | | | | 3 | NA | 31.69 | 25.91 | 26.28 | 27.53 | NA | 96 | 54827 | 5654 | 95.10 | 0.32 | NA |
| | | | | | 4 | NA | 31.39 | 26.09 | 26.49 | 27.34 | NA | 119 | 49388 | 4898 | 95.28 | 0.39 | NA |
| | | | | | 5 | NA | 31.44 | 25.94 | 26.34 | 27.27 | NA | 115 | 53881 | 5427 | 95.21 | 0.38 | NA |
| | Negative control | NC | NA | | 1 | NA | NA | NA | NA | 27.34 | NA | NA | NA | NA | NA | NA | NA |
| | | | | | 2 | NA | 34.48 | NA | NA | 27.1 | NA | 13 | NA | NA | NA | 0.04 | NA |
| | | | | | 3 | NA | 33.63 | NA | NA | 27.12 | NA | 24 | NA | NA | NA | 0.08 | NA |
| | *Gallus gallus domesticus* | NHS_1 | 200 | | 1 | NA | 32.86 | NA | 33.63 | 26.97 | NA | 42 | NA | 37 | NA | 0.14 | NA |
| | | | | | 2 | NA | 34.18 | NA | 33.22 | 27.5 | NA | 16 | NA | 49 | NA | 0.05 | NA |
| | *Sus scrofa domesticus* | NHS_2 | 80 | | 1 | NA | NA | NA | 32.88 | 27.34 | NA | NA | NA | 62 | NA | NA | NA |
| | | | | | 2 | NA | NA | NA | 32.65 | 27.55 | NA | NA | NA | 73 | NA | NA | NA |
| | *Felis catus* | NHS_3 | | | 1 | NA | 34.19 | NA | 33.97 | 27.5 | NA | 16 | NA | 29 | NA | 0.05 | NA |
| | | | | | 2 | NA | 34.34 | NA | NA | 27.56 | NA | 15 | NA | NA | NA | 0.05 | NA |
| | *Canis lupus familiaris* | NHS_4 | | | 1 | NA | NA | NA | 33.13 | 27.44 | NA | NA | NA | 52 | NA | NA | NA |
| | | | | | 2 | NA | NA | NA | 34.56 | 27.43 | NA | NA | NA | 20 | NA | NA | NA |
| | *Rattus* | NHS_5 | | | 1 | NA | NA | NA | 33.2 | 27.42 | NA | NA | NA | 50 | NA | NA | NA |
| | | | | | 2 | NA | 33.73 | NA | 32.24 | 27.45 | NA | 23 | NA | 96 | NA | 0.07 | NA |
| | *Mus musculus* | NHS_6 | | | 1 | NA | 33.48 | NA | 33.32 | 27.49 | NA | 27 | NA | 46 | NA | 0.09 | NA |
| | | | | | 2 | NA | NA | NA | 33.29 | 27.58 | NA | NA | NA | 47 | NA | NA | NA |
| | *Bos taurus* | NHS_7 | | | 1 | NA | NA | NA | 32.49 | 27.35 | NA | NA | NA | 81 | NA | NA | NA |
| | | | | | 2 | NA | 32.89 | NA | 32.46 | 27.12 | NA | 41 | NA | 83 | NA | 0.13 | NA |
| | *Rhesus* | NHS_8 | | | 1 | NA | 30.65 | NA | 31.38 | 27.2 | NA | 201 | NA | 173 | NA | 0.66 | NA |
| | | | | | 2 | NA | 31.01 | NA | 31.17 | 27.41 | NA | 156 | NA | 200 | NA | 0.51 | NA |
| | Positive control | PC | NA | | 1 | 28.15 | 31.99 | 22.69 | NA | 27.68 | 1053 | 78 | 355399 | NA | NA | 0.25 | 0.07 |
| | | | | | 2 | 28.19 | 32 | 22.51 | NA | 27.81 | 1028 | 77 | 394540 | NA | NA | 0.25 | 0.08 |
| | | | | | 3 | NA | 32.54 | 23.37 | NA | 27.17 | NA | 52 | 239495 | NA | NA | 0.17 | NA |
| | | BB_1A | | | 1 | 26.69 | 27.56 | 24.16 | 25.44 | 27.63 | 2009 | 1385 | 146091 | 3282 | 98.89 | 4.55 | 0.62 |
| | | | | | 2 | 26.68 | 27.78 | 24.02 | 25.48 | 27.53 | 2021 | 1185 | 158277 | 3191 | 99.00 | 3.89 | 0.52 |
| | | | | | 3 | 26.97 | 27.71 | 23.89 | 25.44 | 27.42 | 1703 | 1246 | 170502 | 3282 | 99.05 | 4.09 | 0.67 |
| | | BB_1B | | | 1 | 27.53 | 28.59 | 24.92 | 26.01 | 27.67 | 1223 | 669 | 94565 | 2193 | 98.85 | 2.20 | 0.54 |
| | | | | | 2 | 28.05 | 28.49 | 24.86 | 26.43 | 27.47 | 899 | 718 | 97869 | 1629 | 99.17 | 2.36 | 0.83 |
| | | | | | 3 | 27.76 | 28.64 | 24.86 | 26.27 | 27.43 | 1068 | 645 | 97869 | 1824 | 99.08 | 2.12 | 0.61 |
| | | BB_1C | | | 1 | 28.95 | 29.59 | 26.29 | 27.08 | 27.47 | 528 | 330 | 43175 | 1029 | 98.82 | 1.08 | 0.72 |
| | | | | | 2 | 28.77 | 29.52 | 25.71 | 27.09 | 27.24 | 588 | 347 | 60170 | 1021 | 99.16 | 1.14 | 0.67 |
| | | | | | 3 | 27.96 | 29.66 | 25.92 | 26.95 | 27.38 | 949 | 314 | 53357 | 1128 | 98.95 | 1.03 | 0.34 |
| | | BB_1D | | | 1 | 31.88 | 30.82 | 28.08 | 27.23 | 27.2 | 94 | 138 | 15500 | 925 | 97.10 | 0.45 | 2.41 |
| | | | | | 2 | 30.14 | 30.54 | 27.48 | 26.75 | 27.73 | 262 | 168 | 21851 | 1299 | 97.11 | 0.55 | 0.86 |
| | | | | | 3 | 33.16 | 30.43 | 27.55 | 26.75 | 27.54 | 44 | 182 | 20993 | 1299 | 97.00 | 0.60 | 7.84 |
| | | | | | 1 | 33.02 | 32.58 | 30.03 | 28.67 | 27.32 | 48 | 40 | 5078 | 334 | 96.82 | 0.13 | 1.55 |

| | | | | | | | | | | | | | | | | | |
|---|---|---|---|---|---|---|---|---|---|---|---|---|---|---|---|---|---|
| | Human whole blood | BB_1E | 50 | 2 | 2 | 34.34 | 31.6 | 29.26 | 28.44 | 27.23 | 22 | 80 | 7890 | 393 | 97.57 | 0.26 | 7.90 |
| | | | | | 3 | 32.1 | 31.58 | 29.29 | 28.55 | 27.52 | 82 | 81 | 7755 | 364 | 97.71 | 0.27 | 1.64 |
| | | BB_1F | | | 1 | NA | 33.06 | 32.14 | 29.92 | 27.68 | NA | 28 | 1518 | 138 | 95.65 | 0.09 | NA |
| | | | | | 2 | NA | 33.28 | 32.03 | 29.94 | 27.73 | NA | 24 | 1617 | 136 | 95.96 | 0.08 | NA |
| | | | | | 3 | 34.95 | 32.01 | 31.38 | 30.13 | 27.65 | 15 | 60 | 2345 | 119 | 97.53 | 0.20 | 9.10 |
| | | BB_1G | | | 1 | NA | 33.2 | NA | 30.52 | 27.27 | NA | 26 | NA | 90 | NA | 0.08 | NA |
| | | | | | 2 | NA | NA | NA | 31.28 | 28.14 | NA | NA | NA | 53 | NA | NA | NA |
| | | | | | 3 | 17.94 | 32.62 | NA | 30.95 | 27.41 | 353720 | 39 | NA | 67 | NA | 0.13 | 0.00 |
| | | BB_1H | | | 1 | NA | NA | NA | 31.4 | 28.06 | NA | NA | NA | 48 | NA | NA | NA |
| | | | | | 2 | NA | 34.1 | NA | 31.03 | 27.74 | NA | 14 | NA | 63 | NA | 0.04 | NA |
| | | | | | 3 | NA | 33.68 | NA | 31.18 | 27.86 | NA | 18 | NA | 57 | NA | 0.06 | NA |
| | | BB_1I | | | 1 | 29.75 | 33.63 | NA | 29.45 | 27.51 | 329 | 19 | NA | 192 | NA | 0.06 | 0.07 |
| | | | | | 2 | 21.4 | NA | NA | 30.66 | 27.66 | 45778 | NA | NA | 82 | NA | NA | NA |
| | | | | | 3 | NA | NA | NA | 30.57 | 27.94 | NA | NA | NA | 87 | NA | NA | NA |
| | Negative control | NC | NA | | 1 | 24.45 | 34.67 | NA | 32.36 | 28.48 | 7549 | 9 | NA | 25 | NA | 0.03 | 0.00 |
| | | | | | 2 | 25.79 | 31.72 | NA | 31.97 | 28.11 | 3420 | 73 | NA | 32 | NA | 0.24 | 0.02 |
| | | | | | 3 | NA | NA | 18.7 | NA | NA | NA | NA | 3324003 | NA | NA | NA | NA |
| | Positive control | PC | | | 1 | NA | 34.26 | NA | 31.06 | 27.26 | NA | 12 | NA | 62 | NA | 0.04 | NA |
| | | | | | 2 | 26.55 | 31.55 | 18.64 | 32.42 | 28.2 | 2182 | 83 | 3440122 | 24 | 100.00 | 0.27 | 0.03 |
| | | | | | 3 | NA | NA | NA | NA | NA | NA | NA | NA | NA | NA | NA | NA |
| **7500 Fast** | Human whole blood | BB_1A | 50 | 1 | 1 | 27.29 | 27.62 | 19.32 | 24.96 | 28.35 | 1404 | 2838 | 164410 | 4057 | 98.78 | 9.32 | 0.88 |
| | | | | | 2 | 27.46 | 27.79 | 19.86 | 25.26 | 28.76 | 1270 | 2480 | 116285 | 3281 | 98.61 | 8.14 | 0.88 |
| | | | | | 3 | 27.33 | 27.84 | 19.75 | 25.35 | 28.85 | 1371 | 2384 | 124785 | 3079 | 98.78 | 7.83 | 0.76 |
| | | | | | 4 | 27.43 | 26.71 | 19.07 | 25.3 | NA | 1293 | 5840 | 193001 | 3190 | 99.18 | 19.18 | 2.01 |
| | | | | | 5 | 27.22 | 27.32 | 19.33 | 25.12 | 28.07 | 1463 | 3600 | 163359 | 3623 | 98.90 | 11.82 | 1.05 |
| | | BB_1A_UV120 | | | 1 | NA | 30.38 | 26.7 | 27.12 | 28.56 | NA | 318 | 1447 | 880 | 76.68 | 1.04 | NA |
| | | | | | 2 | NA | 30.19 | 26.3 | 27.05 | 29.06 | NA | 370 | 1870 | 925 | 80.18 | 1.21 | NA |
| | | | | | 3 | NA | 30.54 | 26.76 | 27.25 | 28.72 | NA | 280 | 1392 | 803 | 77.63 | 0.92 | NA |
| | | BB_1A_UV30 | | | 1 | NA | 28.23 | 20.77 | 25.1 | 29.25 | NA | 1750 | 64873 | 3675 | 97.25 | 5.75 | NA |
| | | | | | 2 | NA | 28.25 | 20.37 | 24.96 | 29.12 | NA | 1722 | 83845 | 4057 | 97.64 | 5.65 | NA |
| | | | | | 3 | NA | 28.4 | 20.34 | 24.96 | 28.75 | NA | 1529 | 85473 | 4057 | 97.68 | 5.02 | NA |
| | | BB_1A_UV60 | | | 1 | NA | 28.65 | 22.31 | 25.98 | 28.89 | NA | 1254 | 24162 | 1971 | 96.08 | 4.12 | NA |
| | | | | | 2 | NA | 29.08 | 22.22 | 26.35 | 28.95 | NA | 892 | 25598 | 1517 | 97.12 | 2.93 | NA |
| | | | | | 3 | NA | 30.09 | 23.46 | 26.95 | 28.84 | NA | 400 | 11556 | 992 | 95.88 | 1.31 | NA |
| | | BB_1B | | | 1 | 28.07 | 28.05 | 19.84 | 25.46 | 28.26 | 887 | 2018 | 117786 | 2848 | 98.81 | 6.63 | 1.16 |
| | | | | | 2 | 28.09 | 28.42 | 20.13 | 25.34 | 28.5 | 877 | 1505 | 97796 | 3101 | 98.44 | 4.94 | 0.88 |
| | | | | | 3 | 28.33 | 28.4 | 20.21 | 26.38 | 28.28 | 761 | 1529 | 92905 | 1485 | 99.21 | 5.02 | 1.08 |
| | | BB_1C | | | 1 | 29.03 | 29.33 | 20.43 | 25.97 | 28.71 | 504 | 731 | 80679 | 1985 | 98.78 | 2.40 | 0.90 |
| | | | | | 2 | 30.01 | 28.59 | 21.75 | 27.77 | 28.7 | 283 | 1315 | 34602 | 555 | 99.20 | 4.32 | 3.51 |
| | | | | | 3 | 29.11 | 29.2 | 21.11 | 27.37 | 28.58 | 481 | 811 | 52163 | 737 | 99.30 | 2.66 | 1.06 |
| | | BB_1D | | | 1 | 30.21 | 29.65 | 21.28 | 26.84 | 28.26 | 252 | 567 | 46775 | 1073 | 98.87 | 1.86 | 1.77 |
| | | | | | 2 | 30.7 | 29.83 | 22.19 | 28.07 | 28.12 | 189 | 492 | 26095 | 449 | 99.15 | 1.62 | 2.27 |
| | | | | | 3 | 30.35 | 29.95 | 21.94 | 27.65 | 28.09 | 232 | 447 | 30633 | 605 | 99.02 | 1.47 | 1.56 |
| | | BB_1E | | | 1 | 32.2 | 30.73 | 22.62 | 27.98 | 28.74 | 78 | 241 | 19806 | 479 | 98.81 | 0.79 | 3.65 |
| | | | | | 2 | 31.88 | 30.01 | 22.6 | 28.27 | 28.18 | 94 | 426 | 20061 | 390 | 99.04 | 1.40 | 5.02 |
| | | | | | 3 | 30.84 | 30.53 | 23.03 | 28.84 | 28.14 | 174 | 282 | 15226 | 261 | 99.15 | 0.93 | 1.46 |
| | | BB_1F | | | 1 | 30.83 | 30.33 | 23.97 | 29.08 | 28.05 | 175 | 331 | 8333 | 220 | 98.70 | 1.09 | 1.69 |
| | | | | | 2 | 32.46 | 30.74 | 24.21 | 29.38 | 27.35 | 67 | 239 | 7144 | 178 | 98.77 | 0.79 | 4.45 |
| | | | | | 3 | 33.87 | 31.08 | 24.11 | 29.61 | 27.81 | 29 | 183 | 7617 | 151 | 99.02 | 0.60 | 10.40 |
| | | BB_1G | | | 1 | NA | 31.92 | 25.1 | 29.84 | 29.01 | NA | 94 | 4037 | 128 | 98.43 | 0.31 | NA |
| | | | | | 2 | NA | 32.09 | 25.43 | 30 | 29.02 | NA | 82 | 3267 | 115 | 98.28 | 0.27 | NA |
| | | | | | 3 | NA | 31.73 | 25.42 | 29.98 | 28.83 | NA | 109 | 3288 | 116 | 98.26 | 0.36 | NA |
| | | BB_1H | | | 1 | NA | 32.87 | 26.56 | 30.63 | 29.14 | NA | 44 | 1583 | 73 | 97.73 | 0.14 | NA |
| | | | | | 2 | NA | 32.93 | 27.14 | 30.66 | 29.13 | NA | 42 | 1091 | 72 | 96.81 | 0.14 | NA |
| | | | | | 3 | NA | 33.35 | 27.03 | 30.53 | 29.23 | NA | 30 | 1171 | 79 | 96.75 | 0.10 | NA |
| | | BB_1I | | | 1 | NA | 32.7 | 27.63 | 29.82 | 29.11 | NA | 51 | 797 | 130 | 92.45 | 0.17 | NA |
| | | | | | 2 | NA | 32.15 | 28.35 | 30.21 | 28.84 | NA | 78 | 502 | 99 | 91.04 | 0.26 | NA |
| | | | | | 3 | NA | 32.69 | 27.88 | 29.72 | 28.45 | NA | 51 | 679 | 140 | 90.67 | 0.17 | NA |
| | | BB_2A | 5 | | 1 | NA | 30.7 | 23.75 | 28.42 | 28.26 | NA | 247 | 9595 | 351 | 98.21 | 0.81 | NA |
| | | | | | 2 | NA | 30.47 | 23.72 | 28.42 | 28.34 | NA | 296 | 9782 | 351 | 98.24 | 0.97 | NA |
| | | | | | 3 | 32.94 | 30.76 | 23.79 | 28.61 | 28.39 | 50 | 235 | 9352 | 307 | 98.39 | 0.77 | 6.41 |
| | | | | | 4 | NA | 31.35 | 23.84 | 28.56 | 28.67 | NA | 147 | 9057 | 318 | 98.28 | 0.48 | NA |
| | | | | | 5 | NA | 31.35 | 24.11 | 28.49 | 29.1 | NA | 147 | 7617 | 334 | 97.86 | 0.48 | NA |
| | Negative control | NC | NA | | 1 | NA | 33.02 | NA | 31.62 | 29.48 | NA | 39 | NA | 36 | NA | 0.13 | NA |
| | | | | | 2 | NA | 32.21 | NA | 31.28 | 29.51 | NA | 75 | NA | 46 | NA | 0.24 | NA |
| | | | | | 3 | NA | 33.19 | NA | 24.61 | 29.12 | NA | 34 | NA | 5198 | NA | 0.11 | NA |
| | *Gallus gallus domesticus* | NHS_1 | 200 | | 1 | NA | 33.18 | NA | 31 | 29.18 | NA | 35 | NA | 56 | NA | 0.11 | NA |
| | | | | | 2 | NA | 32.53 | NA | 30.65 | 29.45 | NA | 58 | NA | 72 | NA | 0.19 | NA |
| | *Sus scrofa domesticus* | NHS_2 | 80 | | 1 | NA | 34.68 | 33.26 | 29.49 | 29.12 | NA | 11 | 22 | 164 | 20.76 | 0.03 | NA |
| | | | | | 2 | NA | 32.83 | NA | 30.15 | 29.01 | NA | 46 | NA | 103 | NA | 0.15 | NA |
| | *Felis catus* | NHS_3 | | | 1 | NA | 32.94 | NA | 31.04 | 29.26 | NA | 42 | NA | 55 | NA | 0.14 | NA |
| | | | | | 2 | NA | 32.08 | NA | 31.45 | 28.79 | NA | 83 | NA | 41 | NA | 0.27 | NA |
| | *Canis lupus familiaris* | NHS_4 | | | 1 | NA | 32.38 | NA | 31.17 | 28.95 | NA | 65 | NA | 50 | NA | 0.21 | NA |
| | | | | | 2 | NA | 32.72 | NA | 31.18 | 29.23 | NA | 50 | NA | 50 | NA | 0.16 | NA |
| | *Rattus* | NHS_5 | | | 1 | NA | 32.52 | NA | 30.5 | 28.75 | NA | 58 | NA | 80 | NA | 0.19 | NA |
| | | | | | 2 | NA | 32.43 | NA | 30.11 | 28.91 | NA | 63 | NA | 106 | NA | 0.21 | NA |
| | *Mus musculus* | NHS_6 | | | 1 | NA | 32.07 | NA | 30.77 | 28.76 | NA | 83 | NA | 66 | NA | 0.27 | NA |

| | | | | | | | | | | | | | | | | | |
|---|---|---|---|---|---|---|---|---|---|---|---|---|---|---|---|---|---|
| | *Mus musculus* | NHS_[illegible] | | | 2 | NA | 32.27 | NA | 30.52 | 29.13 | NA | 71 | NA | 79 | NA | 0.23 | NA |
| | *Bos taurus* | NHS_7 | | | 1 | NA | 32.55 | NA | 30.53 | 28.86 | NA | 57 | NA | 79 | NA | 0.19 | NA |
| | | | | | 2 | NA | 32.26 | NA | 30.26 | 29.09 | NA | 72 | NA | 95 | NA | 0.24 | NA |
| | *Rhesus* | NHS_8 | | | 1 | NA | 30.52 | 23.41 | 28.62 | 29.15 | NA | 285 | 11933 | 304 | 98.74 | 0.93 | NA |
| | | | | | 2 | NA | 30.44 | 23.15 | 28.6 | 29.1 | NA | 303 | 14098 | 309 | 98.92 | 1.00 | NA |
| | Positive control | PC | NA | | 1 | 27.73 | 30.68 | 17.34 | 31.6 | 29.05 | 1084 | 251 | 585345 | 37 | 100.00 | 0.82 | 0.11 |
| | | | | | 2 | 28.2 | 30.55 | 17.15 | 31.46 | 29.22 | 822 | 278 | 661199 | 41 | 100.00 | 0.91 | 0.18 |
| | | | | | 3 | 27.86 | 30.28 | 17.06 | 31.08 | 29.15 | 1004 | 344 | 700486 | 53 | 100.00 | 1.13 | 0.17 |
| | | BB_1A | 50 | 2 | 1 | 27.53 | 27.74 | 20.13 | 25.87 | 28.11 | 5079 | 2789 | 233686 | 6261 | 98.68 | 9.16 | 0.97 |
| | | | | | 2 | 27.56 | 27.91 | 20.34 | 25.94 | 27.93 | 4974 | 2459 | 202679 | 5979 | 98.55 | 8.08 | 0.88 |
| | | | | | 3 | 27.48 | 27.72 | 20.36 | 25.59 | 27.96 | 5261 | 2831 | 199950 | 7529 | 98.15 | 9.30 | 0.95 |
| | | BB_1B | | | 1 | 27.97 | 28.39 | 20.65 | 26.61 | 27.95 | 3729 | 1724 | 164266 | 3846 | 98.84 | 5.66 | 0.83 |
| | | | | | 2 | 28.57 | 28.1 | 20.9 | 26.7 | 28.11 | 2447 | 2136 | 138659 | 3625 | 98.71 | 7.02 | 1.61 |
| | | | | | 3 | 28.02 | 25.44 | 21.09 | 26.47 | 25.67 | 3600 | 15313 | 121903 | 4218 | 98.30 | 50.28 | 7.69 |
| | | BB_1C | | | 1 | 29.46 | 28.94 | 21.65 | 27.55 | 28.02 | 1309 | 1147 | 83397 | 2071 | 98.77 | 3.77 | 1.67 |
| | | | | | 2 | 27.58 | 27.65 | 21.6 | 27.39 | 25.51 | 4904 | 2981 | 86272 | 2301 | 98.68 | 9.79 | 1.08 |
| | | | | | 3 | 29.21 | 28.38 | 21.82 | 27.44 | 27.56 | 1561 | 1736 | 74320 | 2227 | 98.52 | 5.70 | 2.10 |
| | | BB_1D | | | 1 | 30.83 | 30.17 | 22.28 | 28.07 | 27.54 | 500 | 461 | 54410 | 1470 | 98.67 | 1.51 | 1.86 |
| | | | | | 2 | 30.66 | 28.76 | 22.38 | 28.09 | 27.84 | 564 | 1311 | 50844 | 1451 | 98.59 | 4.30 | 4.65 |
| | | | | | 3 | 30.24 | 26.55 | 22.03 | 28.05 | 27.36 | 757 | 6732 | 64459 | 1490 | 98.86 | 22.10 | 17.49 |
| | | BB_1E | | | 1 | 31.17 | 27.9 | 22.91 | 28.48 | 26.59 | 394 | 2477 | 35499 | 1122 | 98.44 | 8.14 | 12.81 |
| | | | | | 2 | 32.37 | 31 | 23.68 | 28.4 | 27.65 | 170 | 250 | 21063 | 1183 | 97.27 | 0.82 | 3.14 |
| | | | | | 3 | 31.07 | 31.49 | 23.87 | 28.88 | 26.97 | 423 | 174 | 18518 | 863 | 97.72 | 0.57 | 0.83 |
| | | BB_1F | | | 1 | 31.53 | 30.84 | 24.01 | 29.48 | 26.52 | 306 | 281 | 16841 | 581 | 98.30 | 0.92 | 1.90 |
| | | | | | 2 | NA | 31.26 | 24.61 | 29.86 | 27.61 | NA | 206 | 11214 | 452 | 98.02 | 0.68 | NA |
| | | | | | 3 | 33.69 | 31.6 | 24.61 | 30.36 | 27.49 | 67 | 160 | 11214 | 325 | 98.57 | 0.53 | 5.35 |
| | | BB_1G | | | 1 | NA | 32.37 | 25.28 | 26.85 | 26.06 | NA | 90 | 7120 | 3284 | 81.26 | 0.30 | NA |
| | | | | | 2 | NA | 31.44 | 25.66 | 29.97 | 27.7 | NA | 180 | 5503 | 421 | 96.32 | 0.59 | NA |
| | | | | | 3 | NA | 31.29 | 25.27 | 30.27 | 27.21 | NA | 201 | 7169 | 345 | 97.65 | 0.66 | NA |
| | | BB_1H | | | 1 | NA | 32.66 | 27.09 | 30.43 | 28.22 | NA | 73 | 2088 | 311 | 93.07 | 0.24 | NA |
| | | | | | 2 | NA | 32.56 | 27.66 | 30.41 | 28.17 | NA | 79 | 1419 | 315 | 90.01 | 0.26 | NA |
| | | | | | 3 | NA | 33.43 | 27.66 | 30.65 | 28.23 | NA | 41 | 1419 | 269 | 91.34 | 0.14 | NA |
| | | BB_1I | | | 1 | NA | 32.73 | NA | 32.09 | 26.3 | NA | 69 | NA | 104 | NA | 0.23 | NA |
| | | | | | 2 | NA | 32.46 | 27.79 | 29.9 | 27.65 | NA | 85 | 1299 | 441 | 85.50 | 0.28 | NA |
| | | | | | 3 | NA | 32.57 | 27.65 | 29.1 | 28.51 | NA | 78 | 1428 | 746 | 79.29 | 0.26 | NA |
| | Negative control | NC | NA | | 1 | NA | 32.91 | NA | 32.71 | 28.05 | NA | 61 | NA | 69 | NA | 0.20 | NA |
| | | | | | 2 | NA | 23.26 | NA | 33.34 | 20.05 | NA | 76928 | NA | 46 | NA | 252.61 | NA |
| | | | | | 3 | NA | 33.11 | NA | 32.85 | 27.89 | NA | 52 | NA | 63 | NA | 0.17 | NA |
| | | BB_1A | 50 | 1 | 1 | 27.41 | 26.48 | 17.58 | 23.2 | 28.42 | 979 | 948 | 754167 | 2292 | 99.85 | 3.11 | 2.16 |
| | | | | | 2 | 27.6 | 26.69 | 17.69 | 23.46 | 28.57 | 874 | 821 | 699956 | 1909 | 99.86 | 2.70 | 2.13 |
| | | | | | 3 | 26.87 | 26.95 | 18.04 | 23.63 | 28.22 | 1351 | 686 | 552065 | 1694 | 99.85 | 2.25 | 1.08 |
| | | BB_1B | | | 1 | 28.32 | 27.64 | 19.01 | 24.24 | 27.92 | 569 | 427 | 285963 | 1103 | 99.81 | 1.40 | 1.82 |
| | | | | | 2 | 28.12 | 27.43 | 18.85 | 24.15 | 28.35 | 641 | 493 | 318737 | 1176 | 99.82 | 1.62 | 1.83 |
| | | | | | 3 | 27.68 | 27.33 | 18.75 | 24.16 | 28.38 | 833 | 528 | 341101 | 1167 | 99.83 | 1.74 | 1.45 |
| | | BB_1C | | | 1 | 29.47 | 28.54 | 20.02 | 25.02 | 27.92 | 286 | 230 | 144161 | 638 | 99.78 | 0.75 | 2.16 |
| | | | | | 2 | 29.14 | 28.45 | 19.94 | 25.02 | 28.32 | 349 | 244 | 152198 | 638 | 99.79 | 0.80 | 1.83 |
| | | | | | 3 | 28.45 | 27.98 | 19.76 | 24.71 | 27.5 | 526 | 338 | 171958 | 793 | 99.77 | 1.11 | 1.57 |
| | | BB_1D | | | 1 | 29.8 | 29.02 | 20.89 | 25.57 | 27.57 | 235 | 165 | 79913 | 433 | 99.73 | 0.54 | 1.95 |
| | | | | | 2 | 30.44 | 28.76 | 21.37 | 25.75 | 28.07 | 161 | 197 | 57710 | 382 | 99.67 | 0.65 | 3.62 |
| | | | | | 3 | 30.07 | 29.45 | 21.17 | 25.88 | 28.24 | 200 | 123 | 66093 | 349 | 99.74 | 0.40 | 1.74 |
| | | BB_1E | | | 1 | 29.96 | 29.13 | 21.28 | 26.14 | 28.2 | 214 | 153 | 61342 | 290 | 99.76 | 0.50 | 2.02 |
| | | | | | 2 | 30.69 | 29.72 | 21.58 | 26.32 | 28.37 | 138 | 102 | 50050 | 256 | 99.75 | 0.33 | 2.22 |
| | | | | | 3 | 31.87 | 30.09 | 21.51 | 26.23 | 27.42 | 68 | 79 | 52483 | 273 | 99.74 | 0.26 | 3.88 |
| | | BB_1F | | | 1 | 31.34 | 30.37 | 22.7 | 26.56 | 28.3 | 94 | 65 | 23418 | 216 | 99.54 | 0.21 | 2.22 |
| | | | | | 2 | 30.71 | 30.61 | 22.86 | 26.69 | 27.42 | 137 | 55 | 21010 | 197 | 99.53 | 0.18 | 1.22 |
| | | | | | 3 | 32.29 | 30.63 | 23.04 | 26.71 | 27.78 | 53 | 55 | 18596 | 194 | 99.48 | 0.18 | 3.57 |
| | | BB_1G | | | 1 | 34.21 | 30.28 | 23.67 | 27.26 | 28.47 | 17 | 69 | 12130 | 132 | 99.46 | 0.23 | 17.02 |
| | | | | | 2 | 34.67 | 30.39 | 24.19 | 27.61 | 28.18 | 13 | 64 | 8525 | 103 | 99.40 | 0.21 | 21.66 |
| | | | | | 3 | 34.03 | 31.53 | 23.86 | 27.38 | 28.79 | 19 | 29 | 10664 | 121 | 99.43 | 0.10 | 6.36 |
| | | BB_1H | | | 1 | 36.57 | 31.3 | 25.38 | 27.57 | 28.75 | 4 | 34 | 3804 | 106 | 98.62 | 0.11 | 42.81 |
| | | | | | 2 | 32.63 | 31.57 | 25.16 | 27.55 | 27.77 | 43 | 29 | 4416 | 108 | 98.79 | 0.09 | 2.36 |
| | | | | | 3 | 38.02 | 31.36 | 26.18 | 27.76 | 28.21 | 2 | 33 | 2211 | 93 | 97.94 | 0.11 | 111.43 |
| | | BB_1I | | | 1 | 36.6 | 31.67 | 27.05 | 27.99 | 28.02 | 4 | 27 | 1226 | 79 | 96.87 | 0.09 | 33.88 |
| | | | | | 2 | NA | 30.96 | 26.77 | 28.09 | 27.92 | NA | 43 | 1482 | 74 | 97.57 | 0.14 | NA |
| | | | | | 3 | 38.94 | 31.76 | 27.19 | 27.74 | 27.48 | 1 | 25 | 1115 | 94 | 95.94 | 0.08 | 159.38 |
| | Negative control | NC | NA | | 1 | 37.97 | 32.29 | 34.49 | 28.27 | 27.94 | 2 | 17 | 8 | 65 | 19.55 | 0.06 | 56.77 |
| | | | | | 2 | NA | 30.97 | 36.5 | 26.99 | 27.92 | NA | 43 | 2 | 160 | 2.47 | 0.14 | NA |
| | | | | | 3 | NA | 31.93 | 34.93 | 27.97 | 28.14 | NA | 22 | 6 | 80 | 12.74 | 0.07 | NA |
| | Positive control | PC | | | 1 | 27.16 | 29.55 | 15.53 | 27.01 | 28.78 | 1136 | 115 | 3028491 | 158 | 100.00 | 0.38 | 0.22 |
| | | | | | 2 | 27.69 | 29.9 | 15.98 | 28.05 | 28.26 | 828 | 90 | 2231999 | 76 | 100.00 | 0.30 | 0.25 |
| | | | | | 3 | 27.4 | 29.55 | 15.5 | 28.01 | 28.09 | 984 | 115 | 3090735 | 78 | 100.00 | 0.38 | 0.26 |
| | | BB_1A | | | 1 | 27.32 | 26.35 | 17.22 | 23.07 | 28.24 | 1033 | 1037 | 962706 | 2511 | 99.87 | 3.41 | 2.22 |
| | | | | | 2 | 26.98 | 26.61 | 16.99 | 23.38 | 28.39 | 1265 | 867 | 1125207 | 2020 | 99.91 | 2.85 | 1.47 |
| | | | | | 3 | 26.7 | 26.46 | 17.45 | 23.18 | 27.84 | 1495 | 962 | 823673 | 2324 | 99.86 | 3.16 | 1.34 |
| | | BB_1B | | | 1 | 28.56 | 27.34 | 18.16 | 23.91 | 28.47 | 493 | 525 | 508919 | 1392 | 99.86 | 1.72 | 2.64 |
| | | | | | 2 | 28.09 | 27.49 | 18.56 | 24.04 | 28.19 | 652 | 473 | 388009 | 1270 | 99.84 | 1.55 | 1.72 |
| | | | | | 3 | 27.74 | 27.23 | 18.36 | 23.91 | 26.57 | 804 | 566 | 444370 | 1392 | 99.84 | 1.86 | 1.62 |

| | | | | | | | | | | | | | | | | | |
|---|---|---|---|---|---|---|---|---|---|---|---|---|---|---|---|---|---|
| **Rotorgene Q** | Human whole blood | BB_1C | 50 | 2 | 1 | 29.47 | 28.32 | 19.51 | 24.59 | 27.35 | 286 | 267 | 203728 | 863 | 99.79 | 0.88 | 2.51 |
| | | | | | 2 | 28.72 | 28.13 | 19.17 | 24.56 | 27.27 | 448 | 305 | 256559 | 881 | 99.83 | 1.00 | 1.71 |
| | | | | | 3 | 28.69 | 27.99 | 19.29 | 24.48 | 26.23 | 456 | 335 | 236507 | 932 | 99.80 | 1.10 | 1.84 |
| | | BB_1D | | | 1 | 30.36 | 29.02 | 20.81 | 25.53 | 27.52 | 168 | 165 | 84368 | 446 | 99.74 | 0.54 | 2.86 |
| | | | | | 2 | 30.57 | 29 | 20.56 | 25.53 | 27.96 | 149 | 167 | 99956 | 446 | 99.78 | 0.55 | 3.35 |
| | | | | | 3 | 29.88 | 29.23 | 20.31 | 25.56 | 28.11 | 224 | 143 | 118424 | 436 | 99.82 | 0.47 | 1.78 |
| | | BB_1E | | | 1 | 30.86 | 29.97 | 21.25 | 26.71 | 28.46 | 125 | 86 | 62603 | 194 | 99.84 | 0.28 | 2.10 |
| | | | | | 2 | 31.33 | 30.36 | 21.66 | 26.59 | 27.63 | 94 | 66 | 47407 | 212 | 99.78 | 0.22 | 2.22 |
| | | | | | 3 | 31.87 | 29.90 | 21.63 | 26.75 | 28.54 | 68 | 90 | 48381 | 189 | 99.80 | 0.30 | 4.42 |
| | | BB_1F | | | 1 | 32.77 | 30.50 | 23.09 | 27.01 | 28.14 | 40 | 60 | 17976 | 158 | 99.56 | 0.20 | 5.43 |
| | | | | | 2 | 32.29 | 30.74 | 22.60 | 26.97 | 27.69 | 53 | 51 | 25061 | 162 | 99.68 | 0.17 | 3.31 |
| | | | | | 3 | 32.36 | 31.10 | 22.96 | 27.10 | 27.95 | 51 | 39 | 19632 | 148 | 99.62 | 0.13 | 2.71 |
| | | BB_1G | | | 1 | 33.85 | 32.49 | 24.73 | 27.56 | 26.74 | 21 | 15 | 5911 | 107 | 99.10 | 0.05 | 2.90 |
| | | | | | 2 | 34.42 | 30.82 | 24.08 | 27.44 | 27.45 | 15 | 48 | 9186 | 116 | 99.37 | 0.16 | 13.56 |
| | | | | | 3 | 33.59 | 30.95 | 24.03 | 27.47 | 27.58 | 25 | 44 | 9502 | 114 | 99.40 | 0.14 | 7.01 |
| | | BB_1H | | | 1 | NA | 31.49 | 25.15 | 27.92 | 26.88 | NA | 30 | 4446 | 83 | 99.07 | 0.10 | NA |
| | | | | | 2 | 35.98 | 30.88 | 25.25 | 27.85 | 27.37 | 6 | 46 | 4155 | 87 | 98.96 | 0.15 | 38.08 |
| | | | | | 3 | NA | 30.97 | 25.48 | 27.96 | 27.50 | NA | 43 | 3555 | 81 | 98.88 | 0.14 | NA |
| | | BB_1I | | | 1 | NA | 31.84 | 27.02 | 28.41 | 27.4 | NA | 24 | 1251 | 59 | 97.70 | 0.08 | NA |
| | | | | | 2 | NA | 31.9 | 27.64 | 28.53 | 27.77 | NA | 23 | 822 | 54 | 96.81 | 0.07 | NA |
| | | | | | 3 | NA | 31.8 | 27.32 | 28.41 | 27.92 | NA | 24 | 1021 | 59 | 97.20 | 0.08 | NA |
| | *Sus scrofa domesticus* | NHS_2 | 80 | | 1 | NA | 31.33 | NA | 28.66 | 29.21 | NA | 34 | NA | 49 | NA | 0.11 | NA |
| | | | | | 2 | NA | 32.80 | NA | 29.09 | 29.67 | NA | 12 | NA | 37 | NA | 0.04 | NA |
| | *Rattus* | NHS_5 | | | 1 | NA | 31.78 | 38.02 | 25.79 | 27.88 | NA | 25 | 1 | 371 | 0.39 | 0.08 | NA |
| | | | | | 2 | NA | 31.41 | 34.88 | 28.42 | 27.82 | NA | 32 | 6 | 58 | 17.16 | 0.10 | NA |
| | *Mus musculus* | NHS_6 | | | 1 | NA | 32.03 | 33.98 | 28.63 | 27.48 | NA | 21 | 11 | 50 | 30.66 | 0.07 | NA |
| | | | | | 2 | NA | 31.62 | 33.63 | 28.49 | 28.15 | NA | 28 | 14 | 56 | 33.69 | 0.09 | NA |
| | Human whole blood | BB_1A | 50 | 3 | 1 | 27.30 | 27.04 | 17.96 | 23.54 | 28.47 | 1087 | 618 | 562317 | 1771 | 99.84 | 2.03 | 1.36 |
| | | | | | 2 | 27.09 | 26.78 | 17.67 | 23.49 | 28.07 | 1229 | 740 | 686340 | 1834 | 99.87 | 2.43 | 1.41 |
| | | | | | 3 | 27.15 | 26.73 | 17.88 | 23.51 | 28.32 | 1186 | 766 | 594100 | 1809 | 99.85 | 2.51 | 1.52 |
| | | | | | 4 | 27.15 | 26.88 | 18.24 | 23.48 | 28.99 | 1186 | 690 | 463882 | 1847 | 99.80 | 2.27 | 1.37 |
| | | | | | 5 | 27.16 | 26.79 | 17.29 | 23.56 | 28.68 | 1179 | 734 | 891171 | 1747 | 99.90 | 2.41 | 1.47 |
| | | BB_1A_UV120 | | | 1 | 32.92 | 28.09 | 20.69 | 21.53 | 28.57 | 40 | 298 | 86127 | 7200 | 95.99 | 0.98 | 32.44 |
| | | | | | 2 | 32.56 | 28.12 | 20.69 | 21.53 | 28.41 | 50 | 292 | 86127 | 7200 | 95.99 | 0.96 | 24.75 |
| | | | | | 3 | 32.85 | 28.09 | 20.92 | 21.81 | 27.73 | 42 | 298 | 73534 | 5922 | 96.13 | 0.98 | 30.90 |
| | | BB_1A_UV30 | | | 1 | 28.48 | 26.92 | 18.61 | 21.62 | 28.15 | 544 | 671 | 359725 | 6762 | 99.07 | 2.20 | 3.36 |
| | | | | | 2 | 28.38 | 27.22 | 18.57 | 21.99 | 28.60 | 577 | 545 | 369751 | 5223 | 99.30 | 1.79 | 2.55 |
| | | | | | 3 | 28.27 | 26.69 | 18.33 | 21.24 | 28.54 | 616 | 787 | 436058 | 8815 | 99.00 | 2.58 | 3.41 |
| | | BB_1A_UV60 | | | 1 | 30.50 | 28.00 | 19.89 | 23.07 | 28.58 | 167 | 317 | 149252 | 2459 | 99.18 | 1.04 | 6.45 |
| | | | | | 2 | 29.90 | 27.34 | 19.63 | 22.27 | 27.95 | 237 | 502 | 178453 | 4296 | 98.81 | 1.65 | 6.72 |
| | | | | | 3 | 30.82 | 27.58 | 19.65 | 22.26 | 28.36 | 138 | 425 | 176017 | 4326 | 98.79 | 1.39 | 10.77 |
| | | BB_2A | 5 | | 1 | 31.97 | 30.11 | 21.87 | 25.76 | 28.69 | 71 | 73 | 38277 | 376 | 99.51 | 0.24 | 4.14 |
| | | | | | 2 | 32.19 | 30.61 | 22.18 | 25.66 | 27.84 | 62 | 52 | 30932 | 403 | 99.35 | 0.17 | 3.41 |
| | | | | | 3 | 31.77 | 30.25 | 22.36 | 25.65 | 28.40 | 79 | 67 | 27332 | 406 | 99.26 | 0.22 | 3.27 |
| | | | | | 4 | 32.43 | 30.16 | 22.30 | 25.64 | 28.04 | 54 | 71 | 28483 | 409 | 99.29 | 0.23 | 5.50 |
| | | | | | 5 | 32.26 | 30.10 | 21.92 | 25.67 | 28.59 | 60 | 74 | 36984 | 401 | 99.46 | 0.24 | 5.09 |
| | *Gallus gallus domesticus* | NHS_1 | 200 | | 1 | NA | 32.68 | 32.98 | 29.21 | 30.83 | NA | 12 | 18 | 34 | 52.17 | 0.04 | NA |
| | | | | | 2 | NA | 32.62 | 33.24 | 28.84 | 31.19 | NA | 13 | 15 | 44 | 41.34 | 0.04 | NA |
| | *Felis catus* | NHS_3 | 80 | | 1 | NA | 32.60 | 34.58 | 29.07 | 28.74 | NA | 13 | 6 | 37 | 24.78 | 0.04 | NA |
| | | | | | 2 | NA | 31.17 | 36.18 | 29.01 | 28.23 | NA | 35 | 2 | 39 | 9.52 | 0.12 | NA |
| | *Canis lupus familiaris* | NHS_4 | | | 1 | NA | 32.67 | 35.76 | 28.45 | 27.18 | NA | 12 | 3 | 58 | 8.68 | 0.04 | NA |
| | | | | | 2 | NA | 33.37 | 33.90 | 28.70 | 28.79 | NA | 8 | 10 | 48 | 28.88 | 0.03 | NA |
| | *Bos taurus* | NHS_7 | | | 1 | NA | 32.69 | 35.60 | 28.95 | 28.21 | NA | 12 | 3 | 41 | 13.07 | 0.04 | NA |
| | | | | | 2 | NA | 32.90 | 34.29 | 29.27 | 29.21 | NA | 11 | 8 | 33 | 31.62 | 0.03 | NA |
| | *Rhesus* | NHS_8 | | | 1 | 35.63 | 26.70 | 22.90 | 27.86 | 29.20 | 8 | 782 | 18858 | 87 | 99.77 | 2.57 | 557.23 |
| | | | | | 2 | 33.41 | 26.26 | 22.55 | 27.76 | 28.61 | 30 | 1061 | 23987 | 93 | 99.81 | 3.48 | 162.14 |

**S9 Table. Statistical evaluation per performance parameter during the instrument-specific validation of qBiCo-v2.** BC: Bisulfite conversion; p: p-value; p.adj: corrected p-value (Bonferroni correction); p.signif: significance level (*: p < 0.05, **: p < 0.01, ***: p < 0.001, ****: p < 0.0001)

| qBiCo parameter | Instrument | Statistics | Repeatability | Sensitivity - Total | Sensitivity - Pairwise | | | | | | | | Reproducibility - Total | Reproducibility - Pairwise | | | | | | Robustness - Total | Robustness - Pairwise | | |
|---|---|---|---|---|---|---|---|---|---|---|---|---|---|---|---|---|---|---|---|---|---|---|---|
| | | *Method* | *Wilcoxon* | *Kruskal-Wallis* | *T-test* | | | | | | | | *Kruskal-Wallis* | *T-test* | | | | | | *Kruskal-Wallis* | *T-test* | | |
| | | *Group 1* | *50ng* | | *5ng* | *5ng* | *5ng* | *5ng* | *5ng* | *5ng* | *5ng* | *5ng* | | *5ng* | *2.5ng* | *1.25ng* | *0.625ng* | *0.3125ng* | *0.1563ng* | | *0sec* | *0sec* | *0sec* |
| | | *Group 2* | *5ng* | | *2.5ng* | *1.25ng* | *0.625ng* | *0.3125ng* | *0.1563ng* | *0.0781ng* | *0.0391ng* | *0.0195ng* | | *5ng* | *2.5ng* | *1.25ng* | *0.625ng* | *0.3125ng* | *0.1563ng* | | *30sec* | *60sec* | *120sec* |
| **BC efficiency** | **CFX96** | p | 0.0079 | 0.0018 | 0.0861 | 0.0532 | 0.6479 | 0.2665 | 0.0212 | 0.0003 | 0.0579 | 0.0073 | 2.83E-06 | 0.0409 | 0.1075 | 0.0039 | 0.0016 | 0.0729 | 0.0129 | 0.0156 | 0.0443 | 0.0044 | 0.0295 |
| | | p.adj | 0.0079 | 0.0018 | 0.27 | 0.27 | 0.65 | 0.53 | 0.13 | 0.002 | 0.27 | 0.051 | 2.80E-06 | 0.120 | 0.15 | 0.02 | 0.009 | 0.15 | 0.052 | 0.016 | 0.059 | 0.013 | 0.059 |
| | | p.signif | ** | ** | ns | ns | ns | ns | * | *** | ns | ** | **** | * | ns | ** | ** | ns | * | * | * | ** | * |
| | **CFX384** | p | 0.0079 | 0.0072 | 0.394 | 0.192 | 0.258 | 0.254 | 0.369 | 0.001 | 0.018 | 0.058 | 0.593 | 0.5675 | 0.1335 | 0.5253 | 0.731 | 0.9449 | 0.5008 | 0.0243 | 0.0051 | 0.012 | 0.0588 |
| | | p.adj | 0.008 | 0.007 | 1 | 0.96 | 1 | 1 | 1 | 0.005 | 0.12 | 0.35 | 0.59 | 1 | 0.8 | 1 | 1 | 1 | 1 | 0.024 | 0.015 | 0.024 | 0.059 |
| | | p.signif | ** | ** | ns | ns | ns | ns | ns | *** | * | ns | ns | ns | ns | ns | ns | ns | ns | * | ** | * | ns |
| | **QS5** | p | 0.0556 | 0.003 | 0.462 | 0.4095 | 0.0038 | 0.0804 | 0.1877 | 0.0195 | 0.1219 | 0.0061 | 0.03 | 0.475 | 0.4295 | 0.021 | 0.4397 | NA | 0.781 | 0.0273 | 0.1482 | 0.0066 | 0.2157 |
| | | p.adj | 0.056 | 0.003 | 0.82 | 0.82 | 0.03 | 0.4 | 0.560 | 0.12 | 0.49 | 0.042 | 0.03 | 1 | 1 | 0 | 1 | NA | 1 | 0.027 | 0.3 | 0.02 | 0.3 |
| | | p.signif | ns | ** | ns | ns | ** | ns | ns | * | ns | ** | * | ns | ns | * | ns | NA | ns | * | ns | ** | ns |
| | **QS7 Flex** | p | 0.0079 | 0.029 | 0.678 | 0.299 | 0.0002 | 0.0003 | 0.283 | NA | NA | NA | 0.5346 | 0.1279 | 0.5895 | 0.3566 | 0.0112 | 0.0984 | 0.6784 | 0.0249 | 0.0463 | 0.0121 | 0.1446 |
| | | p.adj | 0.008 | 0.029 | 0.85 | 0.85 | 0.001 | 0.001 | 0.85 | NA | NA | NA | 0.53 | 0.51 | 1 | 1 | 0.067 | 0.49 | 1 | 0.025 | 0.093 | 0.036 | 0.14 |
| | | p.signif | ** | * | ns | ns | *** | *** | ns | NA | NA | NA | ns | ns | ns | ns | * | ns | ns | * | * | * | ns |
| | **7500 Fast** | p | 0.0079 | 0.0057 | 0.7184 | 0.1306 | 0.0501 | 0.096 | 0.413 | 0.0075 | 0.0329 | 0.005 | 6.71E-05 | 0.23 | 0.512 | 0.093 | 0.054 | 0.064 | 0.057 | 0.0156 | 0.0061 | 0.0234 | 0.0025 |
| | | p.adj | 0.008 | 0.006 | 0.83 | 0.39 | 0.25 | 0.38 | 0.83 | 0.052 | 0.2 | 0.04 | 6.70E-05 | 0.46 | 0.510 | 0.32 | 0.32 | 0.32 | 0.32 | 0.016 | 0.012 | 0.023 | 0.008 |
| | | p.signif | ** | ** | ns | ns | ns | ns | ns | ** | * | ** | **** | ns | ns | ns | ns | ns | ns | * | ** | * | ** |
| | **RotorgeneQ** | p | 0.0079 | 0.0012 | 0.0138 | 0.0009 | 0.0169 | 0.0004 | 0.0018 | 0.0008 | 0.0329 | 0.023 | 0.1066 | 0.2225 | 0.044 | 0.1335 | 0.1083 | 0.0814 | 0.0662 | 0.0213 | 0.0145 | 0.0001 | 0.0185 |
| | | p.adj | 0.008 | 0.001 | 0.055 | 0.006 | 0.055 | 0.004 | 0.009 | 0.005 | 0.055 | 0.055 | 0.11 | 0.33 | 0.260 | 0.33 | 0.33 | 0.33 | 0.33 | 0.021 | 0.029 | 0.0003 | 0.029 |
| | | p.signif | ** | ** | * | *** | * | *** | ** | *** | * | * | ns | ns | * | ns | ns | ns | ns | * | * | *** | * |
| **BC-DNA concentration** | **CFX96** | p | 0.0014 | 0.0068 | 0.0719 | 0.0991 | 0.2346 | 0.1258 | 0.2156 | 0.0927 | 0.0371 | 0.3683 | 0.0002 | 0.0019 | 0.0006 | 0.0792 | 0.1315 | 0.1243 | 0.1886 | 0.0139 | 0.3195 | 5.07E-05 | 0.2133 |
| | | p.adj | 0.001 | 0.007 | 0.5 | 0.56 | 0.65 | 0.56 | 0.65 | 0.56 | 0.3 | 0.65 | 0.0002 | 0.009 | 0.004 | 0.320 | 0.370 | 0.370 | 0.370 | 0.014 | 0.430 | 0.0002 | 0.430 |
| | | p.signif | ** | ** | ns | ns | ns | ns | ns | ns | * | ns | *** | ** | *** | ns | ns | ns | ns | * | ns | **** | ns |
| | **CFX384** | p | 0.0185 | 9.97E-05 | 0.2467 | 0.312 | 0.0144 | 0.0204 | 0.0002 | 0.6167 | 0.0147 | 0.0682 | 0.9512 | 0.8356 | 0.1939 | 0.7804 | 0.9255 | 0.9439 | 0.8595 | 0.0039 | 0.004 | 0.0451 | 0.0307 |
| | | p.adj | 0.018 | 1.00E-04 | 0.74 | 0.74 | 0.1 | 0.1 | 0.002 | 0.74 | 0.1 | 0.27 | 0.95 | 1 | 1 | 1 | 1 | 1 | 1 | 0.0039 | 0.012 | 0.061 | 0.061 |
| | | p.signif | * | **** | ns | ns | * | * | *** | ns | * | ns | ns | ns | ns | ns | ns | ns | ns | ** | ** | * | * |
| | **QS5** | p | 0.004 | 4.90E-10 | 0.0064 | 0.1081 | 0.5878 | 0.1822 | 0.0763 | 0.0585 | 0.0364 | 0.0534 | 0.5279 | 0.1147 | 0.8296 | 0.254 | 0.3083 | 0.1251 | 0.8971 | 0.005 | 0.081 | 0.0081 | 0.03 |
| | | p.adj | 0.004 | 4.90E-10 | 0.051 | 0.320 | 0.59 | 0.36 | 0.32 | 0.32 | 0.25 | 0.32 | 0.53 | 0.69 | 1 | 1 | 1 | 0.690 | 1 | 0.005 | 0.081 | 0.024 | 0.06 |
| | | p.signif | ** | **** | ** | ns | ns | ns | ns | ns | * | ns | ns | ns | ns | ns | ns | ns | ns | ** | ns | ** | * |
| | **QS7 Flex** | p | 0.3885 | 0.0004 | 0.2378 | 0.3086 | 0.5196 | 0.8542 | 0.9179 | 0.5847 | 0.0388 | 0.1062 | 0.4298 | 0.4483 | 0.0923 | 0.9628 | 0.2872 | 0.643 | 0.9383 | 0.4448 | 0.001 | 0.4268 | 0.0362 |
| | | p.adj | 0.390 | 0.0004 | 1 | 1 | 1 | 1 | 1 | 1 | 0.31 | 0.74 | 0.43 | 1 | 0.55 | 1 | 1 | 1 | 1 | 0.44 | 0.003 | 0.43 | 0.072 |
| | | p.signif | ns | *** | ns | ns | ns | ns | ns | ns | * | ns | ns | ns | ns | ns | ns | ns | ns | ns | *** | ns | * |
| | **7500 Fast** | p | 0.1216 | 2.43E-06 | 0.0317 | 0.125 | 0.2287 | 0.0201 | 0.1056 | 0.0573 | 0.0146 | 0.0431 | 0.0552 | 0.5275 | 0.4021 | 0.1974 | 0.3569 | 0.4804 | 0.563 | 2.38E-05 | 0.0106 | 0.0079 | 0.0029 |
| | | p.adj | 0.120 | 2.40E-06 | 0.19 | 0.32 | 0.32 | 0.14 | 0.32 | 0.23 | 0.12 | 0.22 | 0.055 | 1 | 1 | 1 | 1 | 1 | 1 | 2.40E-05 | 0.016 | 0.016 | 0.0087 |
| | | p.signif | ns | **** | * | ns | ns | * | ns | ns | * | * | ns | ns | ns | ns | ns | ns | ns | **** | * | ** | ** |
| | **Rotorgene Q** | p | 0.4862 | 1.09E-06 | 0.0397 | 0.2051 | 0.183 | 0.0941 | 0.1045 | 0.0023 | 0.0795 | 0.0022 | 0.566 | 0.2154 | 0.3913 | 0.4722 | 0.8956 | 0.2923 | 0.2942 | 0.001 | 0.6623 | 0.012 | 0.0147 |
| | | p.adj | 0.490 | 1.10E-06 | 0.24 | 0.4 | 0.4 | 0.4 | 0.4 | 0.018 | 0.400 | 0.018 | 0.570 | 1 | 1 | 1 | 1 | 1 | 1 | 0.001 | 0.660 | 0.036 | 0.036 |
| | | p.signif | ns | **** | * | ns | ns | ns | ns | ** | ns | ** | ns | ns | ns | ns | ns | ns | ns | *** | ns | * | * |
| **BC-DNA fragmentation** | **CFX96** | p | 0.0079 | 0.2934 | 0.5978 | 0.2647 | 0.2136 | 0.043 | 0.462 | 0.4788 | NA | NA | 6.71E-05 | 0.1117 | 0.0109 | 0.1896 | 0.1647 | 0.0258 | 0.2268 | | | | |
| | | p.adj | 0.0079 | 0.29 | 1 | 1 | 1 | 0.26 | 1 | 1 | NA | NA | 6.70E-05 | 0.45 | 0.065 | 0.49 | 0.49 | 0.13 | 0.49 | | | | |
| | | p.signif | ** | ns | ns | ns | ns | * | ns | ns | NA | NA | **** | ns | * | ns | ns | * | ns | | | | |
| | **CFX384** | p | 1 | 0.0396 | 0.2444 | 0.2447 | 0.0010 | 0.1648 | 0.5196 | 0.0854 | NA | NA | 0.468 | 0.5003 | 0.3175 | 0.8842 | 0.0767 | 0.3453 | 0.976 | | | | |
| | | p.adj | 1 | 0.04 | 0.73 | 0.73 | 0.0057 | 0.66 | 0.73 | 0.43 | NA | NA | 0.47 | 1 | 1 | 1 | 0.46 | 1 | 1 | | | | |
| | | p.signif | ns | * | ns | ns | *** | ns | ns | ns | NA | NA | ns | ns | ns | ns | ns | ns | ns | | | | |
| | **QS5** | p | 0.0079 | 0.0111 | 0.1022 | 0.0316 | 0.0008 | 0.2210 | 0.0405 | NA | NA | NA | 0.0861 | 0.0934 | 0.2418 | 0.0771 | 0.1825 | 0.442 | 0.428 | | | | |
| | | p.adj | 0.0079 | 0.011 | 0.2 | 0.13 | 0.0038 | 0.22 | 0.13 | NA | NA | NA | 0.086 | 0.47 | 0.73 | 0.46 | 0.73 | 0.86 | 0.86 | | | | |
| | | p.signif | ** | * | ns | * | *** | ns | * | NA | NA | NA | ns | ns | ns | ns | ns | ns | ns | | | | |
| | **QS7 Flex** | p | NA | 0.1974 | 0.9168 | 0.6404 | 0.3876 | NA | NA | NA | NA | NA | 0.0250 | 0.0093 | 0.1772 | 0.2344 | 0.8901 | NA | NA | | | | |
| | | p.adj | NA | 0.2 | 1 | 1 | 1 | NA | NA | NA | NA | NA | 0.025 | 0.037 | 0.53 | 0.53 | 0.89 | NA | NA | | | | |
| | | p.signif | NA | ns | ns | ns | ns | NA | NA | NA | NA | NA | * | ** | ns | ns | ns | NA | NA | | | | |
| | **7500 Fast** | p | 0.3333 | 0.0333 | 0.1245 | 0.3640 | 0.0350 | 0.1340 | 0.2103 | NA | NA | NA | 0.409 | 0.1194 | 0.3928 | 0.838 | 0.3309 | 0.612 | 0.5841 | | | | |
| | | p.adj | 0.33 | 0.033 | 0.5 | 0.5 | 0.17 | 0.5 | 0.5 | NA | NA | NA | 0.41 | 0.72 | 1 | 1 | 1 | 1 | 1 | | | | |
| | | p.signif | ns | * | ns | ns | * | ns | ns | NA | NA | NA | ns | ns | ns | ns | ns | ns | ns | | | | |
| | **Rotorgene Q** | p | 0.0079 | 0.0256 | 0.83 | 0.879 | 0.413 | 0.2684 | 0.5274 | 0.0991 | 0.2541 | 0.3721 | 0.3267 | 0.8164 | 0.4705 | 0.6120 | 0.7769 | 0.8384 | 0.2411 | | | | |
| | | p.adj | 0.0079 | 0.026 | 1 | 1 | 1 | 1 | 1 | 0.79 | 1 | 1 | 0.33 | 1 | 1 | 1 | 1 | 1 | 1 | | | | |
| | | p.signif | ** | * | ns | ns | ns | ns | ns | ns | ns | ns | ns | ns | ns | ns | ns | ns | ns | | | | |
